# Overcoming the accuracy–generalization tradeoff in docking and scoring for prospective virtual screening

**DOI:** 10.64898/2026.08.03.742480

**Authors:** Garik Petrosyan, Vahagn Altunyan, Tsolak Ghukasyan, Tigran M. Abramyan, Grigor Arakelov, Aram Davtyan, Tigran Aghajanyan, Iskander Nakipov, Gagik Navasardyan, Arman Fahradyan, Hazarapet Tunanyan, Arman Simonyan, Paweł Ł Janczyk, Das De Silva, Hayk Saribekyan, Lev Tsidilkovski, Vahram Arakelov, Narek Ginoyan, Hasmik Mnatsakanyan, Max Ratnikov, Khachik Smbatyan, Ashot Papoyan, Garegin A. Papoian

## Abstract

Virtual screening promises access to tens of billions of synthetically accessible, diverse compounds, yet it is rarely used as a primary hit-discovery strategy in contemporary drug-discovery campaigns. We argue that this gap reflects the real-world underperformance of the underlying docking and scoring methods: classical docking is generalizable but limited in accuracy by simple functional forms and parsimonious parameterization, whereas recent machine-learning approaches are highly expressive but do not generalize well to novel molecules and pockets, their reported accuracy often inflated by train–test leakage. To address these challenges, we introduce DODock and DOScore, docking and scoring ML/physics hybrid frameworks that are also highly expressive, yet generalize much better out of distribution compared with the prior ML approaches. This generalization has been prospectively tested in several ways. First, DODock’s blind prediction of a drug candidate binding to PCSK9 was compared to the crystal structure that was subsequently solved, recovering the binding pose to 1.2 Å RMSD. We also used DODock and DOScore in prospective virtual screening campaigns against four therapeutic targets, spanning an ectoenzyme (CD73), a kinase (IRAK4), an extended-substrate protease (FXI), and an allosteric inhibition of protein–protein interface (IL17). These screens yielded many chemically novel, biochemically and cellularly active inhibitors. In the case of CD73, which is a historically challenging target for virtual screening, our screen resulted in a roughly hundredfold improvement in hit rate over a recent machine-learning screen. Our results indicate that the apparent ceiling in the accuracy of virtual screening that seemed to have somewhat plateaued in the last two decades is not fundamental, and that structure-based interrogation of ultralarge chemical space may eventually become a credible primary route to novel chemical matter.

## Introduction

Despite decades of academic research and substantial investment by the pharmaceutical industry, virtual screening is rarely reported as the primary hit-discovery strategy behind clinical candidates (recently estimated at ∼1% of hit-finding campaigns (*1*)). Indeed, remarkably few FDA-approved drugs can be unambiguously attributed to virtual screening as the original hit-finding source (*2*) (many of the small number of frequently cited examples may be more generally described as structure-based drug design). This is surprising because, if sufficiently accurate, virtual screening should in principle offer major advantages over experimental screening: ultralarge virtual chemical spaces (*3*) now span tens of billions of synthetically accessible compounds, far exceeding physical screening libraries in both size and structural diversity (*4*). We hypothesize that the limited adoption of virtual screening reflects persistent inaccuracies in its core technologies: docking, which predicts protein–ligand structure, and scoring, which estimates binding affinity. Here we first briefly review both classical and modern machine-learning-based approaches to docking and scoring. We then introduce new docking and scoring methods that substantially reduce these inaccuracies, and evaluate them on a range of retrospective benchmarks as well as in several prospective experimental campaigns.

To a reader unaware of its history, the idea of traditional docking would seem implausible on fundamental physicochemical grounds. Billions or trillions of chemically diverse molecules have complex, idiosyncratic electronic structures that govern their conformational preferences. These conformational preferences, together with many-body, conformer-dependent, and solvent-mediated nonbonded interactions, are unlikely to be accurately captured by the low-dimensional, additive, mostly pairwise functional forms on which classical docking scores rely (*5*). Nevertheless, the theory of simple liquids shows that short-range repulsion (the excluded volume of the molecules) sets liquid structure, while attractive interactions act as a weaker perturbing mean field (*6*, *7*). If one were to extend this concept to biomolecules and hypothesize that molecular recognition is also repulsion-dominated to a substantial degree, then the reliance of early docking algorithms on geometric shape complementarity alone (*8*) would have been well anticipated. Indeed, the native poses were correctly recovered in some cases despite neglecting everything else. Over the subsequent decades, these still very simple scoring functions were greatly elaborated to include hydrogen bonding and hydrophobic interactions, among others, and were paired with a variety of search algorithms, from Monte Carlo and simulated annealing to genetic algorithms (*9*). These elaborations broadened the coverage of docking physics, but largely preserved the additive, pairwise form and parsimonious parameterization. In summary, theoretical arguments anticipate that traditional docking should generalize well but may be fundamentally limited to a relatively low accuracy.

Most modern, pre-machine-learning docking algorithms are reasonably fast, taking seconds to minutes and enabling the screening of billions of compounds with the additional help of activelearning methods. They allow for the internal flexibility of the docked molecule, as well as its translation and rotation, with some methods additionally exploring side-chain motions in the receptor pocket. Using the common definition of successful self-docking, namely a ligand pose predicted within 2 Å heavy-atom RMSD of the crystallographic pose, the most commonly used academic and commercial docking methods achieve success rates of about 40–55% (*10*, *11*). Overcoming this limited performance in pose prediction, together with addressing even larger errors in binding-affinity prediction (scoring is discussed further below), should provide the groundwork for adopting virtual screening as a primary hit-discovery strategy in drug discovery.

The recent wave of machine-learning methods might have been expected to break the accuracy ceiling. Indeed, in the past five years, diffusion-based and cofolding approaches to docking have proliferated, following the success of AlphaFold2 (*12*). Early diffusion-docking methods such as DiffDock (*13*) reported high blind-docking success rates, but independent re-evaluation under stricter controls showed that much of this performance does not carry over to proteins dissimilar to the training set (Figure 1d). The same pattern recurs in more recent cofolding models (AlphaFold3 (*14*), Boltz-1 (*15*), Chai-1 (*16*)), which predict protein and ligand structures jointly. Several of these are evaluated using time-based splits, which exclude structures deposited after a cutoff but not earlier structures that are similar in sequence, fold, or ligand chemistry. Reported accuracy, then might not be reflective of robust generalization. Indeed, when evaluated under explicit structural and chemical similarity filters, the success rates of these models fall steeply with decreasing similarity to the training set, from roughly 80% to below 25% (Fig. 1b). This is unfortunate, because such models are, in principle, well positioned to capture the flexibility and reorganization of a protein pocket upon binding of a bespoke ligand, a long-standing challenge in molecular docking.

**Figure 1:**
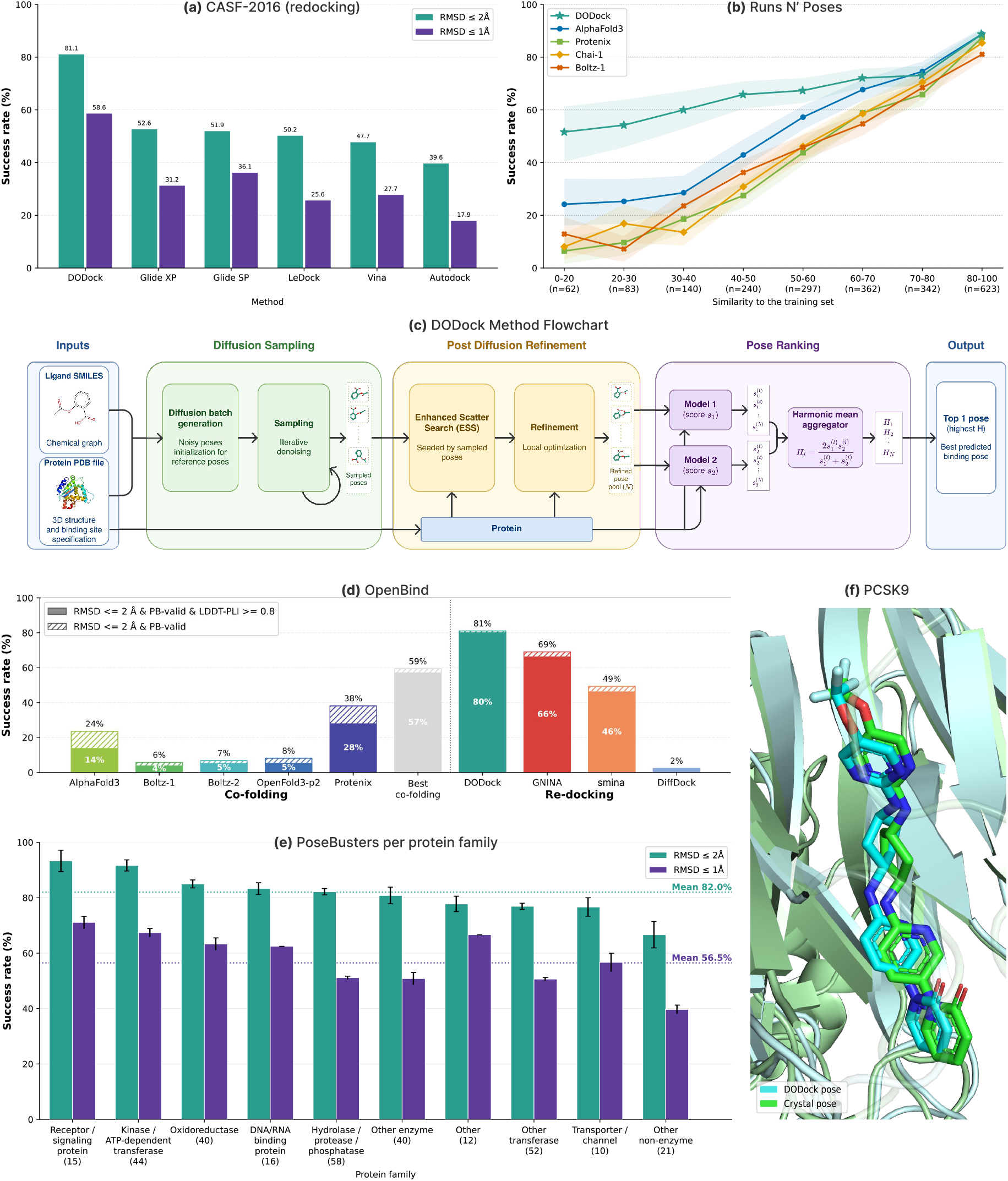
DODock benchmark performance and workflow. **(a)** Success rates on CASF-2016 (redocking), reported at 2 Å and 1 Å RMSD thresholds (*10*, *11*). **(b)** Success rates on the Runs N’ Poses benchmark, shown across similarity bins; success requires RMSD < 2 Å and LDDT-PLI > 0.8. Shaded regions correspond to the 90% confidence interval, calculated from 1,000 bootstrap samples for each bin and method. **(c)** DODock pipeline, comprising diffusion-based pose generation, post-diffusion refinement, and learned pose ranking. **(d)** Success rates on OpenBind benchmark, reported using PB-valid RMSD success and the stricter combined RMSD, PB-validity, and LDDT-PLI criterion. **(e)** DODock performance on PoseBusters benchmark binned by protein family, reported at 2 Å and 1 Å RMSD thresholds. The error bars indicate the standard error of the mean across the three repeat DODock runs. **(f)** Representative PCSK9 docking case showing the crystallographic AZD-0780 pose and the DODock-predicted pose, with a heavy-atom RMSD of ≈ 1.2 Å, in the corresponding pocket.

## Results

### Machine-learning exploration with a physics-based energy landscape

Here, we sketch out the general strategy for DODock, which combines elements of traditional docking with a machine learning ranking model and a diffusion model (Fig. 1c). The latter first proposes plausible candidate poses, supplying broad exploration of pose space. A physics-based search then refines them against an interpretable energy function that is not expected to overfit to training-set similarity because its parameter count is limited to less than 100. The diffusion sampler treats the protein structure as rigid and denoises a ligand pose over its natural degrees of freedom (translation, rotation, and torsional angles) using a score network built on atomic environment vectors (*17*). Because diffusion only proposes initial poses, this diminishes the opportunity for overfitting.

In the subsequent search stage we apply the energy landscape theory of molecular recognition (*18*). We supposed that a well-behaved binding site is characterized by a binding mini-funnel, where, despite local ruggedness, diverse starting poses follow a free energy gradient towards the native pose (*18*, *19*). Next, we relied on the general functional form of the AutoDock Vina potential (*20*), however, increasing its expressive power by expanding its parameter count to 80 by making each coefficient specific to interacting atom-type pairs. Because this is still highly parsimonious, this choice helps to avoid overfitting. We optimized the new docking potential in light of the theories of protein folding and binding – with a funnel objective that drives near-native poses to low energy while suppressing competing decoy minima. To traverse this funnel we adapted and improved upon enhanced scatter search (*21*), a powerful global optimizer, combined with local refinement. A separate ranking neural network, trained on the same funnel logic to score how faithfully a pose reproduces native protein–ligand contacts, then selects the final structure. Together these choices were designed to largely preserve the generalizability of a low-parameter physical model, while providing new avenues for enhancing the overall accuracy. DODock architecture and training are described in detail in SI’s Materials and Methods Section 7.

### DODock generalizes across protein and chemical space

The CASF-2016 benchmark set (*22*) (285 high-resolution protein-ligand crystal structures grouped into 57 sequence-clustered targets, each represented by ligands spanning a wide affinity range) has been widely used for over the last decade to assess various docking and scoring algorithms, evaluated most often through the official CASF rescoring protocols (*22*). Here, in what we refer to as CASF-2016 (redocking), we evaluated the harder task of docking from scratch, requiring each method to both sample and identify the crystallographic pose as its top-ranked solution, and applied a strict protein- and ligand-similarity split to DODock’s training data. Under these conditions, DODock predicted the native pose to within 2 Å for 81.1% of complexes, exceeding widely used commercial and academic programs by more than 25 percentage points (Fig. 1a). A substantial gap remains at near-crystallographic resolution: DODock reached 58.6% within 1 Å, where these methods fell below 37%. The details of the similarity-based train-test split are provided in SI’s Materials and Methods Section 5.

Runs N’ Poses (*23*) comprises 2,600 high-resolution protein-ligand complexes deposited in the Protein Data Bank after the 30 September 2021, the cutoff date shared by many cofolding methods. Each complex is binned by the similarity of the ligand and pocket with that of the closest training example, facilitating the assessment of method’s generalizability. Because success on this benchmark requires both an accurate ligand pose (RMSD < 2 Å) and an accurate protein-ligand interface (LDDT-PLI > 0.8), it tests structure prediction more stringently than ligand-RMSD alone. Fig. 1b demonstrates that the cofolding methods AlphaFold3, Boltz-1, Protenix, and Chai-1 do not generalize well, succeeding on 75-88% of the most similar complexes but falling to 25% or below on the most dissimilar. Interestingly, Fig. S3 suggests that across most of the similarity range co-folding methods maintain the quality of the protein pocket prediction well, while it is specifically the docking outcomes that deteriorate. DODock matches the performance of the cofolding methods on the most similar complexes, at 89%, while showing a much gentler decline as complexes become more novel, exceeding 50% even in the least-similar bin (Fig. 1b). In virtual screening discovering novel chemical matter is the main objective, hence, the performance of docking methods on the novel molecules and pockets (left side) of the Runs N’ Poses plot has the most practical impact.

OpenBind (*24*) is a recent open-science effort to generate dense, experimentally determined protein-ligand data on targets deliberately chosen to be dissimilar from existing structures. Its inaugural release comprises 925 crystallographic binding events for the Enterovirus A71 2A protease, a viral target whose complexes fall far from the pre-2021 structures on which current cofolding models were trained (Fig. 1d). This novelty provided a stringent prospective test of generalization, where the cofolding models reached top-1 success rates of only 4-28% (AlphaFold3, 14%; Protenix, 28%; Boltz-1, 4%), and the diffusion-docking method DiffDock essentially fails (2%). DODock, when trained under a strict protein- and ligand-similarity split specifically with respect to the OpenBind test set, performs best of all methods evaluated, reaching 80% top-1 success for self docking (exceeding also the performance of the CNN-based GNINA, 66%, and the conventional docking program SMINA, 46%). Interestingly, even though DODock is also a highly expressive machine-learning method, it successfully generalizes to unfamiliar complexes.

To assess whether DODock may excel on a few well-represented protein classes, such as kinases, but fail on other protein classes, we evaluated it on the PoseBusters (*25*) benchmark set under the same strict similarity split and stratified the results by protein family (Fig. 1e). Performance was consistently high across diverse families: success rates within 2 Å ranged from roughly 76% to 93% across receptor and signaling proteins, oxidoreductases, hydrolases and proteases, kinases, transporters and channels, and nucleic-acid-binding proteins, with a mean of 82%. Notably, kinases and other ATP dependent transferases fell squarely within this band (91.6%). The same evenness held at near-crystallographic resolution, where success within 1 Å averaged 56.5% across families. DODock’s accuracy is therefore not concentrated in a few data-rich target classes but extends across the structural and functional diversity of the proteome, a prerequisite for general-purpose application in drug discovery.

We note that in real-world virtual screening the pocket is most often well established from prior structural and mechanistic work. Most results in this work therefore assume that the identity of the pocket is known. Blind docking against the whole protein, however, degrades performance only modestly (see SI Section 7.7).

Full benchmark results and analyses are provided in the SI’s Supplementary Text Section 1.

### Prospective application to PCSK9

As a stringent prospective test, we applied DODock to PCSK9, a central regulator of plasma LDL-cholesterol and one of the most actively-pursued targets in cardiovascular drug discovery. Because PCSK9 lowers LDL-receptor levels through a protein–protein interaction, it has long resisted small-molecule inhibition. The recent emergence of orally bioavailable agents therefore represents a significant medicinal-chemistry advance (*26*). The most prominent, AstraZeneca’s AZD0780 (laroprovstat), does not act by blocking PCSK9 binding to the LDL receptor, instead it stabilizes the PCSK9 C-terminal domain, preventing lysosomal trafficking and degradation of the receptor (*26*). AZD0780 is now in a Phase III cardiovascular outcomes trial.

At the time of our prediction, no experimental coordinates for the PCSK9–AZD0780 complex were available in the Protein Data Bank or any other public repository. We generated a blind pose prediction with DODock, timestamped February 24, 2025, followed by an effort to determine the complex’s crystal structure experimentally. The structure was successfully solved on October 20, 2025, at 2.07 Å resolution (we have deposited the corresponding data in the Protein Data Bank under PDB ID 37HF). The prediction therefore constitutes a carefully documented prospective test against a subsequently determined experimental structure. We note that AstraZeneca’s structure of this complex (PDB ID 9HZ3) was deposited on 13 January 2025 but held from public release until 27 May 2026 (*27*).

Using only the protein structure and the chemical identity of AZD0780 as input, DODock placed the ligand in the C-terminal cysteine-rich domain of PCSK9. This site is distinct from the well-characterized catalytic and LDLR-binding surfaces and is sparsely represented among known small-molecule complexes. The predicted pose reproduced the experimentally observed binding mode to within approximately 1.2 Å heavy-atom RMSD (Fig. 1f), recovering both the ligand’s orientation and its key protein contacts. Even though this is only one experiment, it still provides valuable insights. Achieving near-crystallographic accuracy on an unusual binding site, without even distant analogues to guide the placement, suggests that DODock’s generalization extends to genuine prospective cross-docking.

PCSK9 experimental methodology and analysis are described in SI’s Material and Methods Section 9.1.

### Docking-derived data enables generalizable scoring

Accurate pose prediction is important, but only the first step of virtual screening. It is followed by scoring, which is supposed to correlate with thermodynamic binding affinity from a docked complex. Scoring has historically been intractable, showing poor correlation with experimental measurements in prospective testing. Classical scoring functions inherit the same additive, pairwise functional forms as docking, limiting their predictive power. On the other hand, machine-learning scoring functions, though more expressive, are constrained by data. There are relatively abundant data on various affinity measurements; however, reliable associated three-dimensional binding-pose data are lacking.

Consequently, models are often trained on very limited structural data. Exacerbating this challenge, leakage between training and test sets inflates results on enrichment benchmarks such as DUD-E (*28*) and DEKOIS 2.0 (*29*), where train–test separations that filter on protein sequence or ligand identity alone frequently fail to remove near-duplicate proteins, folds, or related chemotypes (*30*). Apparent success thus partly reflects memorization rather than generalization (the same pattern we observed for docking, Fig. 1b). Realistic estimates of screening performance therefore require stringent, simultaneous filtering on both protein and ligand (*30*).

We reasoned that if large affinity datasets could be paired with accurate three-dimensional structures of protein-ligand complexes, this would significantly improve the accuracy and generalizability of the scoring model. On this view, accurate docking is a prerequisite for training an accurate scoring model. Following this strategy, we used DODock to build structural training data at scale (Fig. 2a). Starting from assays with experimental affinities and defined binding sites, we processed the corresponding full protein structures from the PDB, docked each annotated ligand into its assay-specific pocket, and retained the top-ranked pose. To support hit identification, we additionally generated property-matched but topologically dissimilar decoys and docked them into the same pockets. This program generated a large corpus of affinity-labelled three-dimensional complexes.

**Figure 2:**
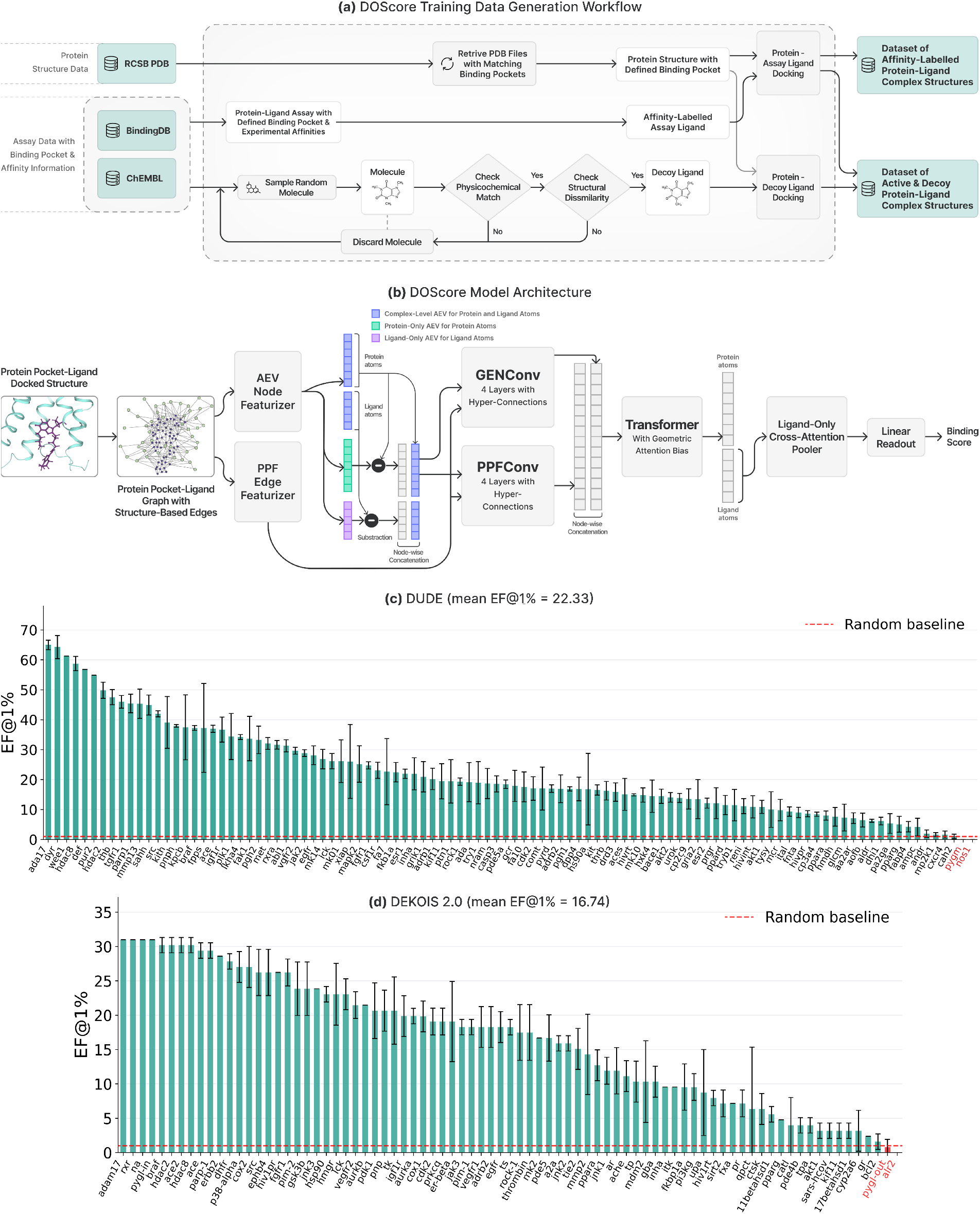
DOScore overview and results. **(a)** The scoring data generation method for training DOScore. **(b)** The architecture of DOScore model. **(c)** DOScore’s per-target enrichment factors for DUD-E benchmark. **(d)** DOScore’s per-target enrichment factors for DEKOIS 2.0 benchmark. In sub-figures c) and d) EF@1% quantifies how many times more true actives are within the top 1% DOScore’s ranking of the ligands compared to a completely random selection from the target’s entire ligand set; results are reported as mean ± standard deviation across three independent runs; red highlights targets with mean EF@1% < 1 (i.e. worse than random).

On these data, we trained DOScore, a neural network scoring model that represents each complex as an atomic graph whose edges are defined by spatial proximity rather than covalent bonds, capturing non-bonded and long-range interactions (Fig. 2b). Local atomic environments are encoded with atomic environment vectors (*17*) and processed by a graph neural network, augmented with hyper-connections to counter over-smoothing, before a transformer with a distance-dependent attention bias aggregates global interaction context. Ligand-focused attention pooling then reads out a binding score. Decoys are handled by an asymmetric loss that penalizes only false positives, preparing the model for the ranking task that screening demands. Scoring data generation, DOScore model architecture, and training are described in Supplementary Information, Materials and Methods, Section 8.

We evaluated DOScore on two established virtual-screening benchmarks, DUD-E and DEKOIS 2.0, under a deliberately stringent split that removes from training every protein similar to a test protein (structural alignment with sequence identity above 0.3) and every ligand similar to a test ligand (Tanimoto similarity above 0.4 on Morgan fingerprints). Even under this strict separation, DOScore enriched actives strongly and consistently across targets: it exceeded an EF@1% of 10 for a large majority of targets in both benchmarks, with only two targets in each falling below the random baseline (Fig. 2c, d).

Performance was broadly distributed rather than concentrated in a few well-studied protein classes, mirroring the cross-family evenness of the docking results. We note that many published methods report higher mean enrichments on these benchmarks, but under similarity filters as strict as ours, such figures typically fall. When we relax our cutoffs to match the protein-only splits used elsewhere, DOScore’s enrichment rises accordingly and exceeds that of recent methods evaluated under the same conditions. We regard the strict-split numbers as the more realistic estimate of how DOScore will perform on genuinely novel chemical space, the regime that matters most for prospective screening. Beyond enrichment, DOScore also correlates well with binding affinity itself: on the JACS, Merck, and OpenBind affinity benchmarks it matches or exceeds recent specialized methods under comparable similarity splits. Full analyses, including experiments in which the similarity cutoffs are progressively relaxed, are provided in Supplementary Text, Section 2.

### Prospective virtual screening across ligandable and protein–protein-interaction targets

Retrospective benchmarks show that DOScore provides high enrichments under strict similarity control. However, an even more important test for drug discovery is prospective: given only a target structure and a virtual library, can the workflow produce genuinely novel binders? We ran prospective campaigns against four targets chosen to represent distinct protein classes and inhibition modalities. CD73 and IRAK4 present orthosteric or catalytic pockets (Fig. 3a, b), Factor XIa a catalytic site that natively engages an extended peptide substrate (Fig. 3c), and IL-17A a pocket whose occupancy should disrupt a distal protein–protein interface. In the latter two cases, a small molecule must compete against the far greater number of contacts made by a peptide or protein partner. In the case of IL-17A, we sought an allosteric PPI disruptor, further increasing the difficulty.

**Figure 3:**
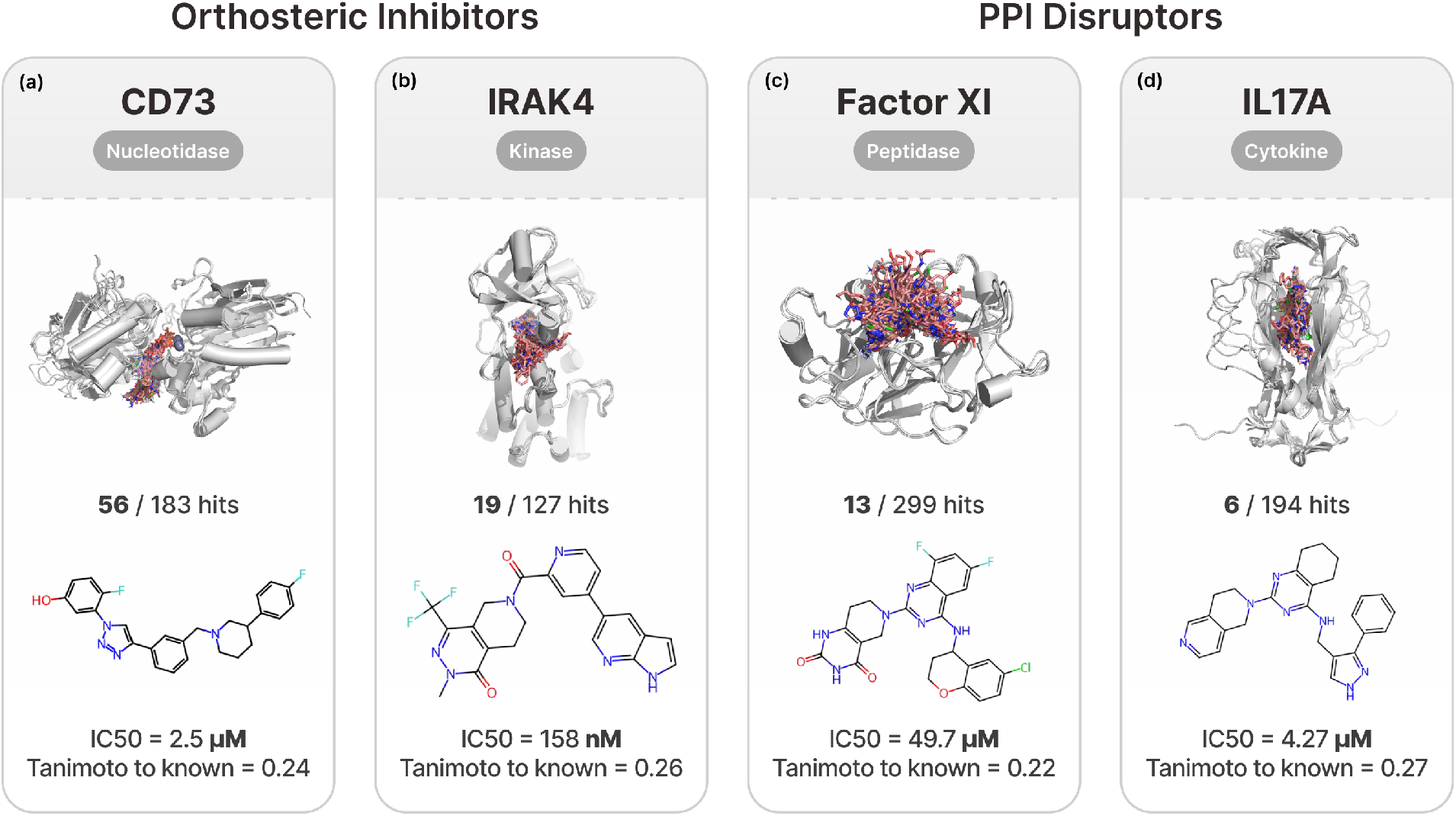
Prospective virtual screening identifies novel inhibitors across diverse target classes. Prospective virtual screening campaigns were conducted against four therapeutically relevant targets representing distinct target classes and binding-site architectures. CD73 (nucleotidase), IRAK4 (kinase), and Factor XI (FXIa protease) were targeted through orthosteric inhibition, whereas IL-17A, a protein–protein interaction (PPI) target, was targeted through an allosteric binding site. For each target, the experimentally determined hit rate, a representative validated hit, its biochemical potency (*IC*50), and its structural similarity to the closest previously reported inhibitor (ECFP4 Tanimoto coefficient) are shown. The prospective campaigns identified 56 active compounds from 183 tested for CD73, 19 from 127 for IRAK4, 13 from 299 for Factor XI, and 6 from 194 for IL-17A. Across all four targets, representative hits exhibited low structural similarity to known inhibitors (ECFP4 Tanimoto coefficients of 0.22–0.27), demonstrating the ability of the virtual screening workflow to prospectively identify chemically novel scaffolds across both conventional orthosteric targets and more challenging protein–protein interaction targets.

Each target was clinically-validated or genetically-supported, presented a characterized binding site, and lacked an approved small-molecule therapy at the time of screening. For each, we screened an ultra-large make-on-demand library and selected 100-300 compounds for experimental testing.

#### CD73 (NT5E)

CD73 (Fig. 3a) is the principal ectoenzyme generating immunosuppressive adenosine in the tumor microenvironment. Clinical development is led by quemliclustat (AB680; *K*_*i*_ ≈ 5 pM), with oral programs LY3475070 and ORIC-533 (*31*, *32*). Nearly all of these agents, including quemliclustat, descend from the nucleotide scaffolds first built around the substrate analog AMPCP, and the dinuclear-zinc active site strongly favors phosphate-bearing ligands. Given the potential cross-reactivity of that compound class, novel non-nucleotide scaffolds are of considerable interest, and were the goal of our virtual screening.

#### IRAK4

IRAK4 (Fig. 3b) is a clinically validated kinase in Toll-like receptor signaling, with the inhibitors zimlovisertib (*33*), zabedosertib (*34*), and emavusertib (*35*), and the degraders KT-474 (*36*) and KT-413 (*37*). Given a dense preclinical literature of ATP-competitive chemotypes (*38*), we asked whether our virtual screening could identify novel scaffolds.

#### Factor XIa (FXIa)

FXIa (Fig. 3c) is a leading next-generation anticoagulation target, motivated by genetics showing that FXI deficiency lowers thrombotic risk without the bleeding liability of factor Xa and thrombin inhibitors (*39*, *40*). Unlike a compact enzyme pocket, the FXIa active site is adapted to bind an extended peptide substrate, so a small molecule must recover, within a few hundred daltons, the affinity that the peptide draws from contacts across the cleft. This is *a priori* a harder problem than CD73 or IRAK4. The recent halt of the Phase III asundexian trial (*41*) for inferior efficacy suggests that even this validated target still needs chemically distinct, selective inhibitors.

#### IL-17A

IL-17A (Fig. 3d) is a validated inflammatory cytokine whose blockade by biologics is efficacious across autoimmune diseases (*42*, *43*). Reproducing this effect with an oral small molecule is far harder: binding of a small molecule in a distal pocket must propagate to the protein surface and disrupt extended contacts at the protein–protein interface. DC-806, from DICE Therapeutics, provided the first clinical proof that direct small-molecule inhibition of a cytokine is achievable (*44*). Our virtual screening therefore targeted novel allosteric PPI inhibitors of IL-17A.

Across all four, our virtual screening based on DODock and DOScore identified chemically novel hits (Tanimoto ≈ 0.22–0.27 to known ligands; see Fig. 3). Overall virtual screening workflow, experimental validation, as well as the IRAK4, Factor XI, and IL-17A campaigns, are described in SI’s Materials and Methods Section 9.2. We next examine the CD73 campaign in detail.

### Prospective discovery of novel CD73 inhibitors

CD73 is a conformationally-dynamic ectoenzyme whose active site is unusually difficult to model. Its two domains undergo a large rigid-body rotation between an open, catalytically inactive state and a closed, competent state, and the two states interconvert during turnover (*45*). The closed state coordinates two Zn^2+^ ions and, in some catalytic intermediates but not others, a bound phosphate is present. A subpocket near L415 is occupied by ordered water across all CD73 crystal structures, irrespective of ligand or conformation.

In our CD73 campaign, we selected the closed conformation as the primary target while also requiring some affinity for the open conformation. We modeled the bound phosphate and disallowed disruption of the ordered-water subpocket near L415. We docked against an ensemble of experimentally determined conformations to account for the pocket’s structural plasticity. Interactions conserved across all CD73 co-crystal structures were imposed as three pharmacophore constraints: an aromatic contact with F417 and F500, a hydrogen-bond acceptor to N390, and a hydrogen-bond donor to D506.

Against this receptor model we screened roughly 80 billion synthetically accessible compounds in Enamine REAL Space, yielding 183 compounds that were acquired and experimentally assayed (Fig. 4a). Of these, 56 inhibited CD73 with a biochemical IC_50_ below 500 *μ*M, 54 below 100 *μ*M and 9 below 10 *μ*M; 14 were active in cells, one at 570 nM potency (see Fig. S6). The hits were highly diverse (Fig. S6), and were structurally unrelated to previously reported CD73 inhibitors (Fig. 4b).

**Figure 4:**
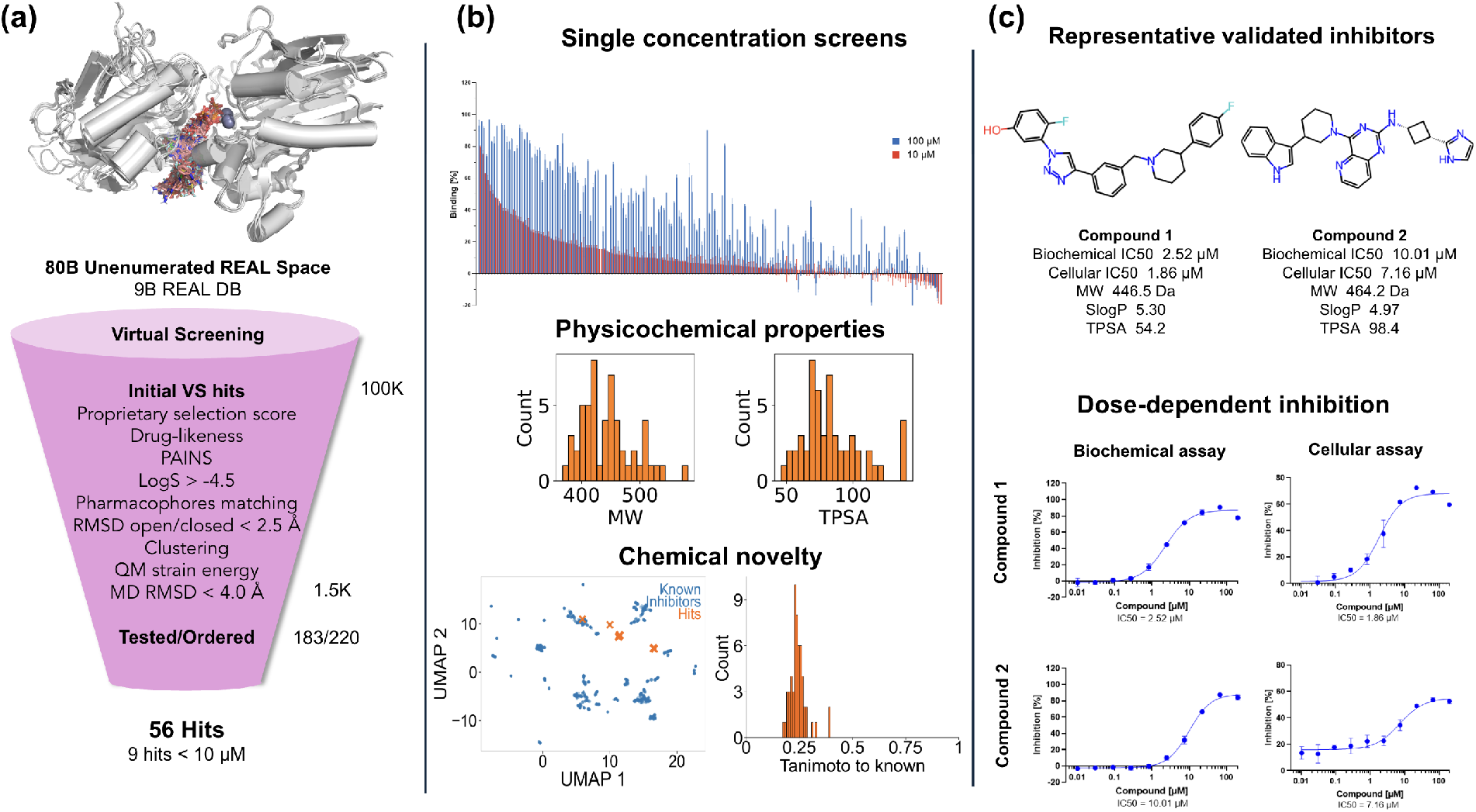
Structure-based virtual screening and experimental validation of CD73 inhibitors. **(a)** Prospective virtual screening workflow. Approximately 80 billion compounds from Enamine REAL Space were computationally screened and progressively prioritized to 220 compounds for procurement, of which 183 were experimentally evaluated. The number of compounds remaining after each filtering stage and the corresponding biochemical and cellular hit rates are shown. **(b)** Characterization of validated hits. Top, single-concentration biochemical inhibition measured at 100 and 10*μM*. Middle, molecular weight (MW) and topological polar surface area (TPSA) distributions of validated hits. Bottom left, UMAP projection of ECFP4 fingerprints comparing validated hits (orange) with previously reported CD73 inhibitors (blue). Bottom right, distribution of ECFP4 Tanimoto similarity between validated hits and the closest known CD73 inhibitor, illustrating the chemical novelty of the identified compounds. **(c)** Representative validated CD73 inhibitors. Chemical structures, biochemical and cellular *IC*50 values, physicochemical properties, and representative dose-response curves for two non-nucleotide inhibitors.

For comparison, an earlier CD73 virtual screening returned nucleoside-mimetic hits from a library preselected for similarity to known inhibitors, with no novel scaffolds being identified (*46*). CD73 was also among the least tractable targets in the largest prospective virtual screening study reported to date, in which the AtomNet platform was applied to 318 targets and returned confirmed hits for the majority of them (*47*). Against CD73 (NT5E), 335 compounds were tested and a single inhibitor was confirmed in dose–response, at 176 *μ*M (a hit rate of 0.3% and the weakest potency reported for any of their 22 internal targets). At a matched potency threshold of 100 *μ*M, that campaign returned no hits from its primary screen, whereas our screen returned 54 of 183, a hit rate of ∼30%.

This at least hundredfold increase in hit rate may partly reflect differences in library size (16 billion compounds in their screen against ∼80 billion REAL Space in ours), receptor preparation, screening strategy and assay format. On the other hand, an increase of this magnitude is what the retrospective results anticipate. DODock recovered poses accurately on systems with no close training analogues, where classical docking programs and co-folding models both degrade (Fig. 1). DOScore then retained its ranking power once protein and ligand similarity to the training data were removed (Fig. 2). The CD73 campaign demonstrates that virtual screening can now identify chemically novel, biochemically- and cellularly-active inhibitors from a pocket that has otherwise resisted it.

## Discussion

The persistent underperformance of virtual screening, we have argued, does not reflect a fundamental barrier to modeling molecular recognition, but rather specific weaknesses of two dominant approaches. Classical docking is generalizable yet strongly capped in accuracy by its largely additive, pairwise form and highly parsimonious parameterization. Newer machine-learning methods, by contrast, are more expressive but do not perform well out of distribution. DODock and DOScore were designed to achieve the high accuracy of machine-learning methods on in-distribution systems while still performing reasonably well on novel ligands and pockets. Indeed, DODock’s accuracy degrades gracefully with distance from the training set, falling only from 89% on the most similar complexes to roughly 50% on wholly novel pockets and ligands, whereas co-folding models fall from the mid-80s to below 25% across the same range. DOScore enriches actives strongly under simultaneous protein- and ligand-similarity filtering, a regime in which the scoring functions we compared against lose most of their enrichment.

Retrospective benchmark results are further supported by our prospective campaigns. We predicted a blind PCSK9 pose to within approximately 1.2 Å heavy-atom RMSD at an atypical binding site. Against CD73, a well-documented difficult target for which a prior large-scale machine-learning screen reported a 0.3% hit rate, virtual screening with DODock and DOScore gave a hit rate of ∼30%. For IL-17A, our screen identified allosteric inhibitors that disrupt a distal protein–protein interface, an objective among the most difficult in small-molecule discovery.

These findings suggest that the marginal role of virtual screening in hit discovery reflects a tractable accuracy problem rather than an inherent limit, though important caveats remain. The compounds reported here are hit-stage starting points (the most potent reaching low-triple-digit nanomolar activity against CD73 and IRAK4) rather than clinical candidates. Converting such hits into medicines still depends on medicinal-chemistry optimization downstream of the screen, although computational modeling can contribute substantially there as well. Performance is also target-dependent: the extended, peptide-adapted active site of Factor XIa gave a predictably lower hit rate than the more conventional enzyme pocket of IRAK4, a reminder that recognition against surfaces evolved to engage large partners is intrinsically harder. Binding-affinity prediction (scoring) remains the natural target for further gains.

Should docking and scoring shift to generalization instead of memorization, structure-based interrogation of tens of billions of synthetically accessible compounds may become a routine rather than exceptional route to novel chemical matter, closing the long-standing gap between the promise of virtual chemical space and its realized use in drug discovery.

## Acknowledgments

We thank Michael Antonov for championing this work, Elin Barnes for managing the validation experiments, and Natalie Ma for helpful discussions and feedback. We are grateful to Erik Arabyan for his help in reviewing the targets for the validation studies.

## Supplementary Materials

### Supplementary Text

#### 1 Docking

##### 1.1 Additional benchmarks

**Figure S1:**
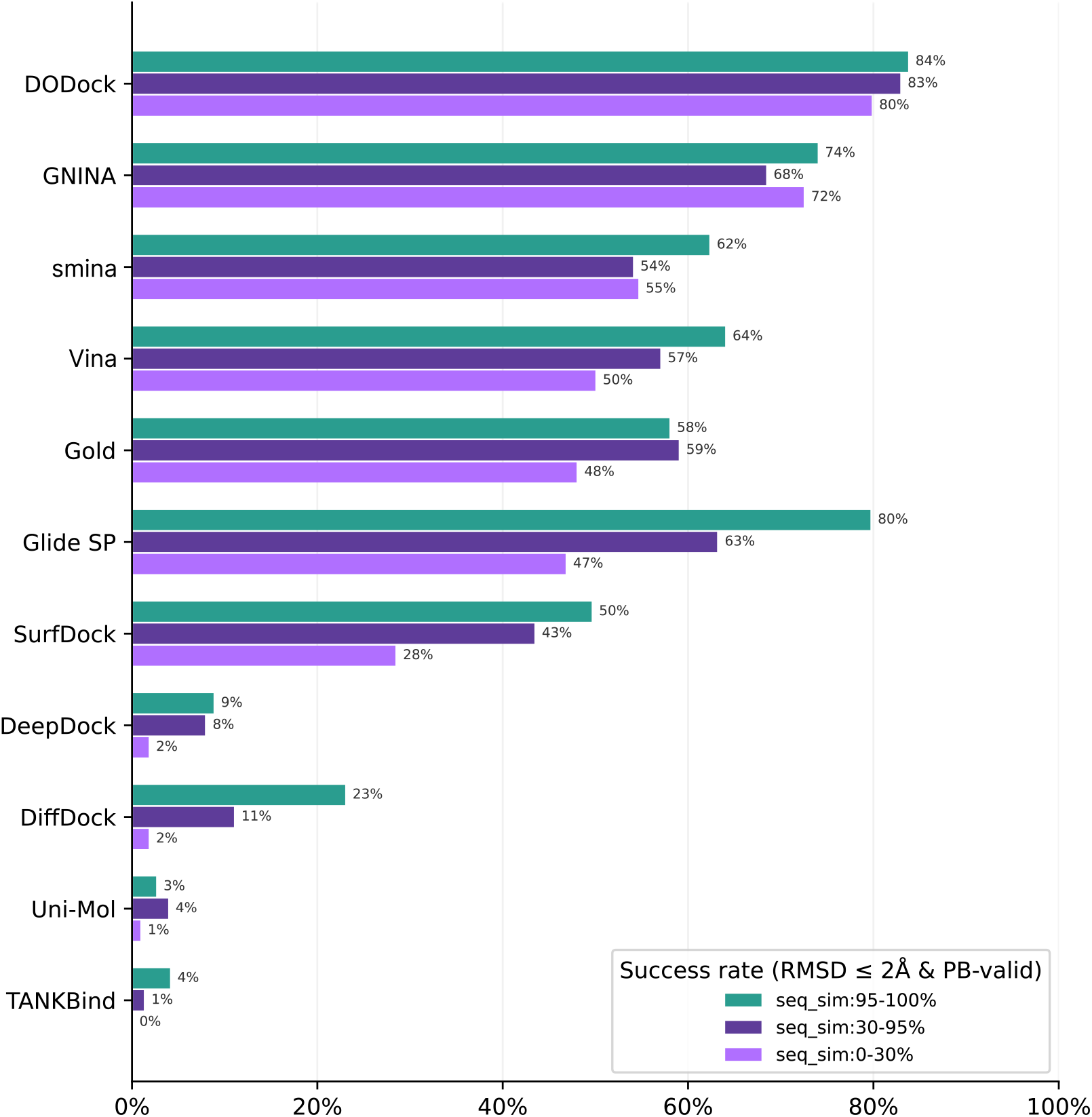
Docking performance on the PoseBusters benchmark across protein sequence-similarity bins. Success requires a symmetry-corrected heavy-atom RMSD of at most 2 Å and passage of all applicable PoseBusters validity checks.

We additionally evaluated DODock on the PoseBusters benchmark using a model trained on PDBbind v2020 complexes. This setup follows the temporal-split evaluation protocol used by other docking methods. We evaluated the complete benchmark as provided, without additional filtering.

Performance was stratified according to protein sequence similarity to the training set. We applied the strict PoseBusters-valid docking success criterion: a prediction was considered successful only if it had a symmetry-corrected heavy-atom RMSD of at most 2 Å relative to the experimentally resolved ligand pose and passed all applicable PoseBusters validity checks. The reported success rate therefore represents the fraction of predictions that are both geometrically accurate and physically plausible.

DODock achieved the highest success rate in every sequence-similarity bin, with success rates of 84%, 83%, and 80% in the 95–100%, 30–95%, and 0–30% bins, respectively. Its performance decreased by only four percentage points between the highest- and lowest-similarity bins. The strongest competing method achieved 80%, 68%, and 72% in the corresponding bins, giving DODock improvements of four, fifteen, and eight percentage points, respectively. These results indicate that DODock’s performance remains comparatively stable as sequence similarity to the training set decreases.

We next extended our Runs N’ Poses evaluation, which previously compared DODock with cofolding methods, by evaluating three established redocking tools on the same benchmark: AutoDock Vina v1.2.7 (*1*), smina v2017.11.9 (*2*), and GNINA v1.3.3 (*3*).

We started from the 2,149-complex evaluation subset defined in Section 5.3. We further restricted the analysis to complexes for which AutoDock Vina, smina, and GNINA all completed successfully, yielding a final paired evaluation subset of 2,142 complexes.

Initial ligand conformations were generated using the RDKit (*4*) ETKDGv3 conformer-generation procedure. Conversion from SDF to PDBQT format was performed using Meeko v0.7.1 (*5*). For each redocking method, we used a cubic search box with an edge length of 25 Å, centered at the centroid of the crystallographic ligand. All methods used an exhaustiveness of 8 and returned up to nine poses per run.

smina was run with the Vinardo scoring function (*6*). Because its default Vina scoring function produced results nearly identical to those of AutoDock Vina, we report the Vinardo configuration as a non-redundant baseline. All other options were left at their default values.

**Figure S2:**
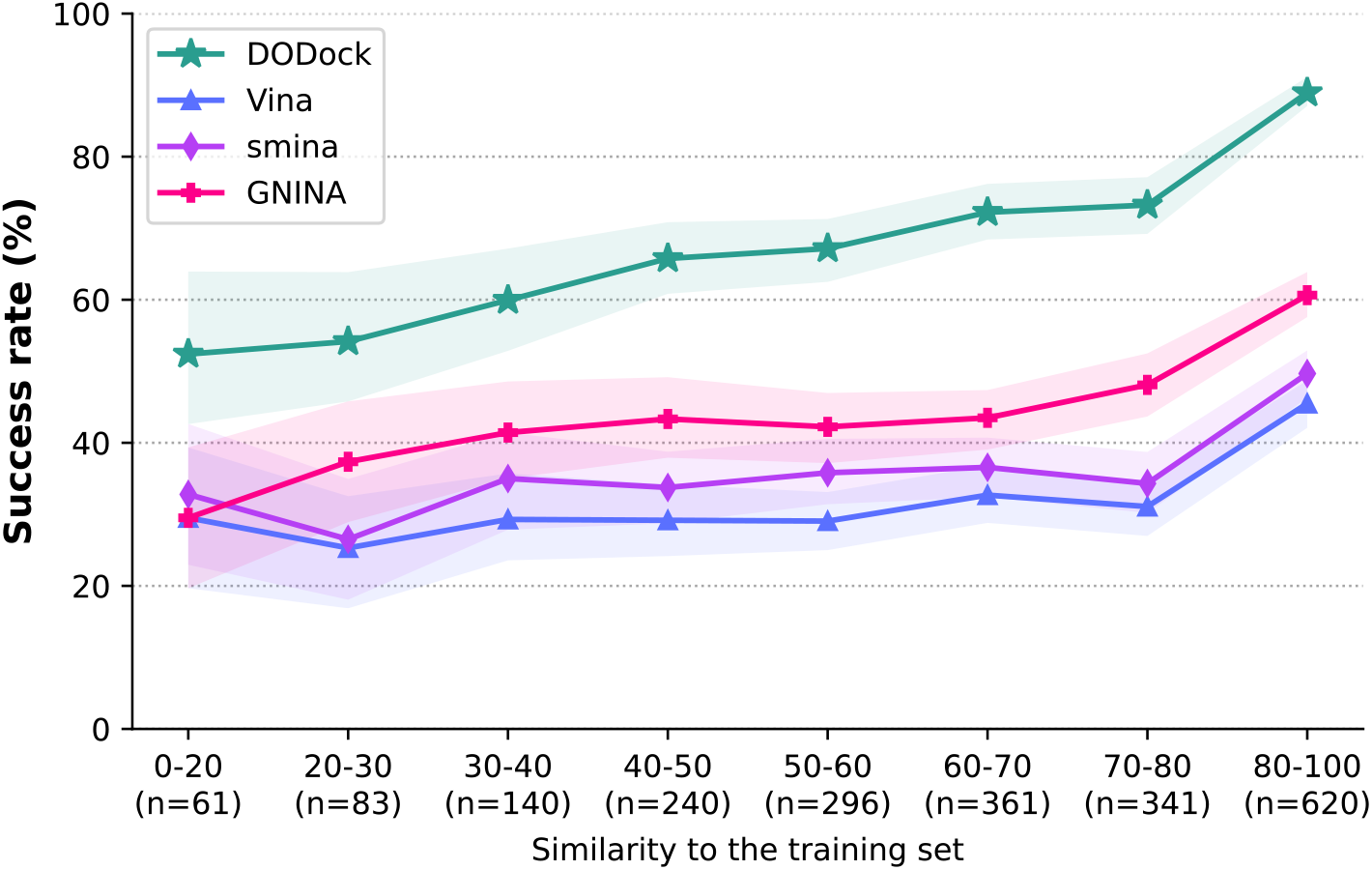
Comparison of DODock with conventional redocking methods on the Runs N’ Poses benchmark. AutoDock Vina, smina, and GNINA were evaluated using the same crystallographically defined search region. Success rates are reported as a function of protein similarity to the training set.

DODock achieved the highest success rate across every sequence-similarity bin. In the most dissimilar bin (0–20%), DODock achieved a success rate of approximately 52%, compared with approximately 33% for the strongest redocking baseline, smina–Vinardo. In the 80–100% bin, the corresponding rates were approximately 89% for DODock and 61% for GNINA. GNINA was the strongest conventional baseline in all bins above 20% similarity, while DODock maintained an advantage of approximately 17–29 percentage points across the benchmark. Thus, DODock’s advantage was maintained for both proteins closely related to and substantially different from those in the training set.

##### 1.2 Ablation studies

We performed three groups of experiments to characterize the docking pipeline. First, we ablated DODock’s three principal components: diffusion-based pose sampling, post-diffusion refinement, and learned pose ranking. Second, we evaluated sensitivity to prior binding-site information by comparing pocket-defined and blind docking. Third, we examined the runtime–accuracy trade-off under different diffusion-sampling budgets.

###### 1.2.1 Pipeline-component ablations

To quantify the contribution of each component of DODock, we ablated each of its three stages in turn, either removing it or replacing it with a simplified baseline while keeping the remaining stages unchanged.

All component ablations were evaluated on the PoseBusters benchmark set using models trained under the PDBbind v2020 time-split setting. We used the default sampling configuration of *N*_confs_ = 4 starting ligand conformers and *N*_traj_ = 4 independently sampled reverse-diffusion trajectories per conformer, yielding *N*_total_ = 16 candidate poses. The same input structures and evaluation procedure were used throughout. Results are reported as the Top-1 success rate, where a prediction was considered successful if the top-ranked pose had a symmetry-corrected ligand heavy-atom RMSD below 2 Å.

###### Ablating diffusion-based sampling

To evaluate the contribution of diffusion-based sampling, we replaced the learned diffusion model with a random pose generator that sampled the same number of ligand poses within the specified binding region. These poses were subsequently processed using the same post-diffusion refinement and pose-ranking stages as in the complete pipeline. This ablation tests whether refinement and ranking alone are sufficient to recover accurate top-ranked poses when learned pose-generation is replaced by random sampling (Table S1).

**Table S1:** Effect of diffusion-based pose sampling. Comparison of diffusion-based and random pose generation before post-diffusion refinement. Results are reported for the PoseBusters benchmark set using the PDBbind v2020 time-split setting.

| Pose source | Top-1 success (%) | Mean runtime (s) |
| --- | --- | --- |
| Diffusion-based sampling | **85.0** | 180 |
| Random sampling | 81.8 | 200 |

###### Ablating post-diffusion refinement

To evaluate the contribution of post-diffusion refinement, we removed this stage and passed the poses generated by the diffusion model directly to the pose-ranking module. This ablation measures how much the global and local optimization procedures improve the final ranked poses beyond those produced by diffusion-based sampling alone (Table S2).

**Table S2:** Effect of post-diffusion refinement. Comparison of the pipeline with and without post-diffusion refinement. Results are reported for the PoseBusters benchmark set using the PDBbind v2020 time-split setting.

| Post-diffusion refinement | Top-1 success (%) | Mean runtime (s) |
| --- | --- | --- |
| Included | **85.0** | 180 |
| Omitted | 69.8 | 80.4 |

###### Ablating learned pose ranking

To evaluate the contribution of learned pose ranking, we replaced the two-model ranking procedure with direct ranking by the intermolecular component of the DOFast score. Diffusion-based pose sampling and post-diffusion refinement were kept unchanged. This ablation measures the improvement provided by the learned ranking models over direct selection using the DOFast interaction score (Table S3).

**Table S3:** Effect of pose ranking. Comparison of learned pose ranking with direct selection using the DOFast interaction score. Results are reported for the PoseBusters benchmark set using the PDBbind v2020 time-split setting.

| Ranking method | Top-1 success (%) | Mean runtime (s) |
| --- | --- | --- |
| Learned ranking models | **85.0** | 180 |
| DOFast interaction score | 65.78 | 140 |

###### 1.2.2 Effect of binding-site information

DODock’s default docking protocol assumes that the target binding pocket is provided as input. To evaluate the dependence of docking performance on this prior information, we compared the pocket-defined protocol with the blind-docking procedure described in Section 7.7. This comparison was performed on both PoseBusters and OpenBind benchmark sets to assess the effect of binding-site information across benchmarks with different compositions (Tables S4 and S5).

**Table S4:**
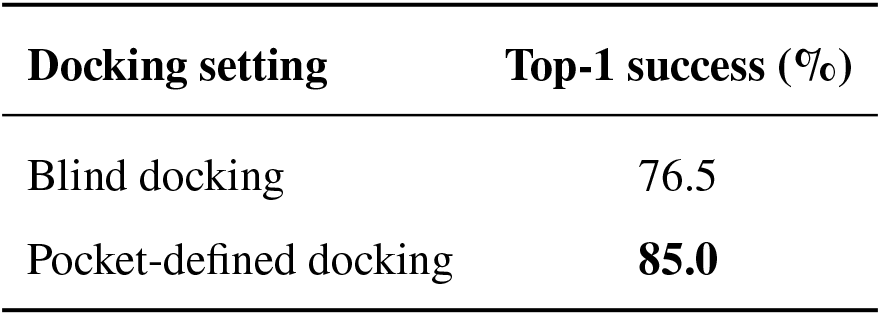
Effect of binding-site specification on PoseBusters. Comparison of blind and pocket-defined docking using the model trained under the PDBbind v2020 time-split setting.

**Table S5:**
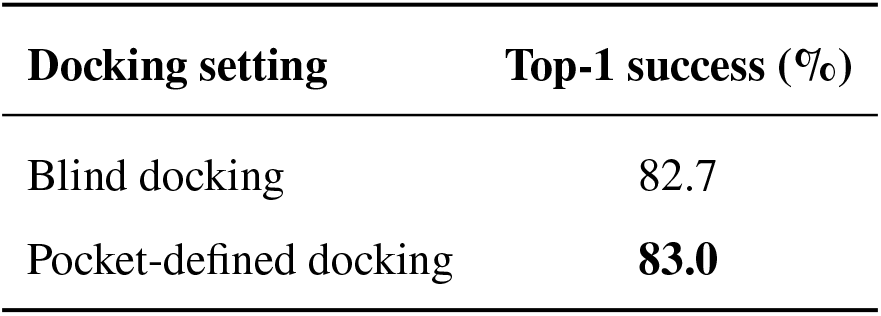
Effect of binding-site specification on OpenBind. Comparison of blind and pocket-defined docking using the corresponding benchmark-specific similarity-filtered model.

###### 1.2.3 Runtime–accuracy trade-off

To characterize the runtime–accuracy trade-off, we varied the number of starting ligand conformers *N*_confs_ and the number of independently sampled reverse-diffusion trajectories per conformer *N*_traj_, yielding *N*_total_ = *N*_confs_ × *N*_traj_ candidate poses per configuration. Each protein–ligand docking calculation was executed as a single-threaded process using one physical core of an AMD EPYC 9654 processor. For each configuration, the average runtime was computed as the mean per-complex wall-clock runtime across the benchmark runs. The results are reported in Table S6.

**Table S6:** Runtime–accuracy trade-off on PoseBusters. Results for the model trained under the PDBbind v2020 time-split setting as the number of starting conformers *N*_confs_ and independently sampled reversediffusion trajectories per conformer *N*_traj_ were varied. Each configuration generated *N*_total_= *N*_confs_ × *N*_traj_ candidate poses. All reported values are averages over two independent benchmark runs.

| $N_{\text{confs}}$ | $N_{\text{traj}}$ | Top-1 success (%) | Mean runtime (s) |
| --- | --- | --- | --- |
| 1 | 1 | 73.7 | 21.7 |
| 1 | 4 | 76.9 | 30.9 |
| 2 | 1 | 80.5 | 34.9 |
| 2 | 4 | 82.1 | 65.5 |
| 3 | 1 | 82.0 | 40.9 |
| 3 | 4 | 84.6 | 104.1 |
| 4 | 1 | 83.3 | 57.1 |
| 4 | 4 | **85.0** | 180 |
| 5 | 1 | 83.1 | 74.4 |

##### 1.3 DOFast scoring results

Search algorithms in DODock use the scoring function DOFast (described in Section 7.3). To isolate the effect of the scoring function, we held the Monte Carlo search procedure fixed and varied only the score used to guide optimization. Specifically, we performed separate searches using the Vina and DOFast scoring functions on CASF-2016 (redocking; Section 5.3.1) and PoseBusters (Section 5.3.1) benchmark sets. Both experiments used the Monte Carlo procedure described in Algorithm 5.

For each run, the final pose was selected from the generated candidates according to one of four criteria: RMSD, pose rank score, score, or energy score. Selection by RMSD is an oracle criterion because it requires knowledge of the crystallographic pose and therefore measures the sampling ceiling of the optimization procedure. The remaining criteria measure practical pose-ranking performance. A prediction was considered successful when the selected pose had a symmetry-corrected ligand heavy-atom RMSD below 2 Å from the crystallographic pose. The resulting optimization and pose-selection performance is summarized in Table S7.

**Table S7:**
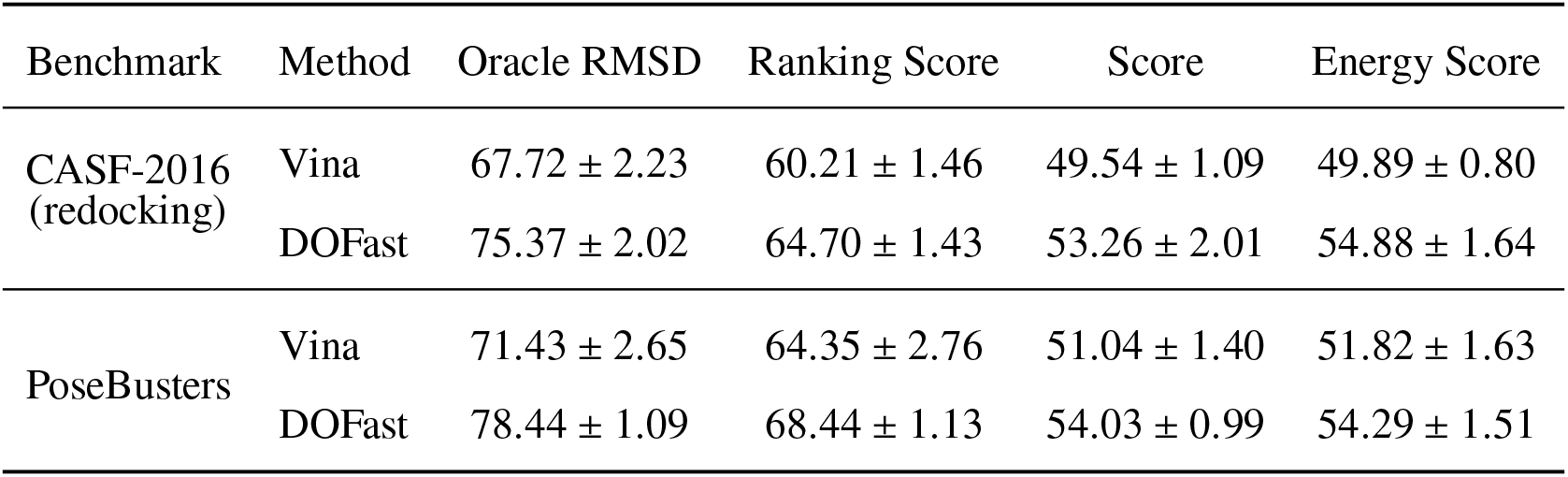
Monte Carlo optimization performance using the Vina and DOFast scoring functions. Values are success rates (%) at the 2 Å symmetry-corrected RMSD threshold of the structure, selected by each of the following metrics: Oracle RMSD (structure with minimal RMSD), Ranking Score (described in Section 7.5), method-specific Score (Vina or DOFast, respectively), method-specific Energy Score (derived from the corresponding intermolecular score according to Eq. S49), reported as the mean ± standard deviation over five independent runs.

##### 1.4 Binding-site and ligand-pose accuracy of cofolding methods

**Figure S3:**
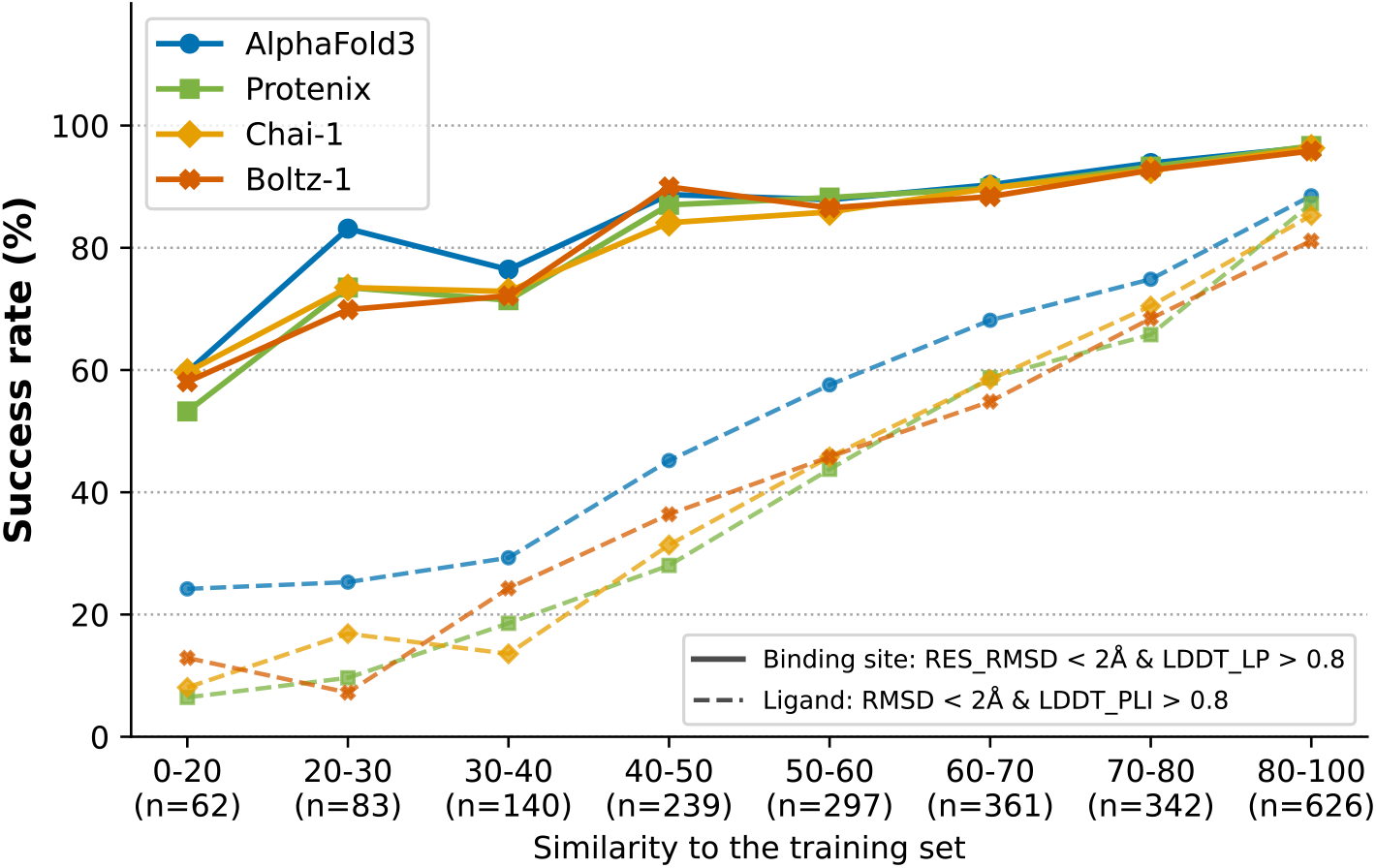
Binding-site versus ligand pose accuracy of cofolding methods as a function of training-set similarity. Success rate on 2,150 RunsN’Poses complexes, binned by combined ligand-shape/pocket-coverage similarity to the nearest training system (bin sizes in parentheses). Solid lines: binding-site success (full-residue RMSD < 2 Å and LDDT-LP > 0.8). Dashed lines: ligand success (RMSD < 2 Å and LDDT-PLI > 0.8). One confidence-ranked top-1 pose per complex (ipTM), identical complex set across all methods.

Additionally, to separate a cofolding model’s ability to locate and build the correct binding pocket from its ability to place the ligand correctly within it, we scored every prediction under two independent criteria: a binding-site criterion (full-residue pocket RMSD < 2 Å and LDDT-LP > 0.8) and a ligand criterion (ligand RMSD < 2 Å and LDDT-PLI > 0.8). Four cofolding methods - AlphaFold3, Protenix, Chai-1 and Boltz-1 - were evaluated on the RunsN’Poses benchmark, restricted to proper (drug-like) ligands passing a 3 Å clash filter and further intersected across all four methods so that every method is scored on an identical set of 2,150 protein–ligand complexes (Fig. S3). That the pocket degrades only mildly across most of the similarity range while the ligand pose degrades severely indicates that the bottleneck is the docking, not the folding.

#### 2 Scoring

##### 2.1 Additional benchmarks

###### 2.1.1 Impact of different train-test splits

We evaluated DOScore using various train-test split methods. The results (Fig. S4) show that using a protein-only split with a less strict 90% sequence identity cutoff (as described in Section 5) dramatically inflates DOScore’s test performance: average enrichment estimation increases from 16.74 (under our rigorous split) to 22.87 with the default model selection scheme.

**Figure S4:**
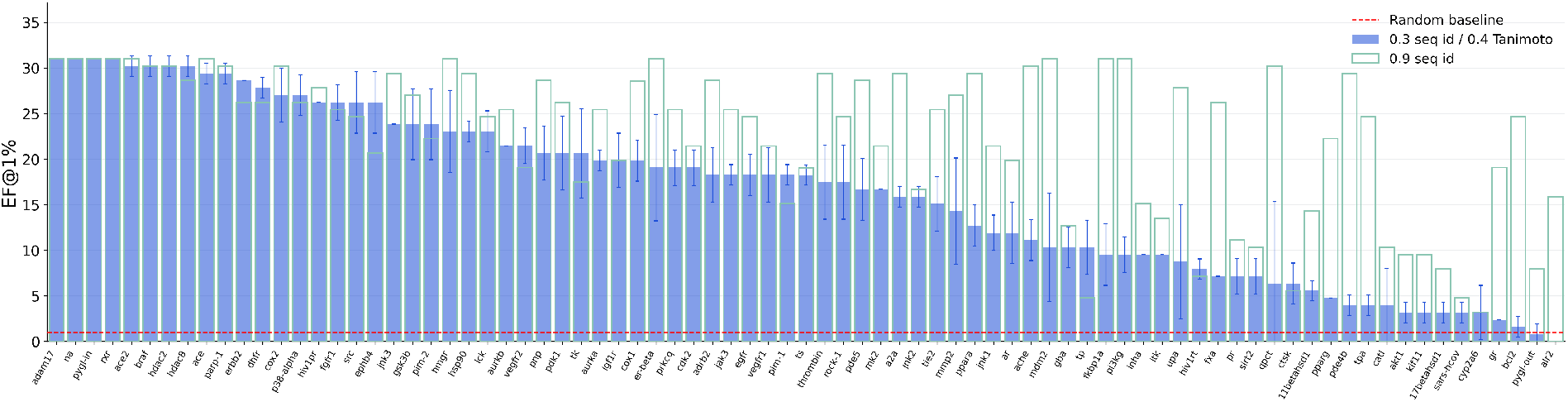
DEKOIS 2.0 performance with different train-test split methods. Our default train-test split method is denoted “0.3 seq id / 0.4 Tanimoto", while the protein-only split with 90% sequence identity cutoff is denoted “0.9 seq id". For each split method we repeated the training with three different random seeds.

Furthermore, the training dynamics (Fig. S5) indicate that evaluating training progress and selecting checkpoints under a weak-similarity split may favor models that increasingly memorize the training data rather than generalize. For example, under our default model-selection scheme (Section 8.3), which averages checkpoints from 2,000 to 10,000 training, the average DEKOIS 2.0 EF@1% for the protein-only 90% split is 22.87. Averaging later checkpoints, from 15,000 to 25,000 steps, further increases the enrichment to 24.93. In contrast, under the more rigorous 30% protein and 0.4 ligand split, the default model-selection scheme achieves an EF@1% of 16.74, whereas averaging the later checkpoints reduces the enrichment to 15.24.

These findings underscore the importance of rigorous train–test splits for avoiding overoptimistic performance estimates and ensuring that checkpoint selection prioritizes generalization rather than memorization.

**Figure S5:**
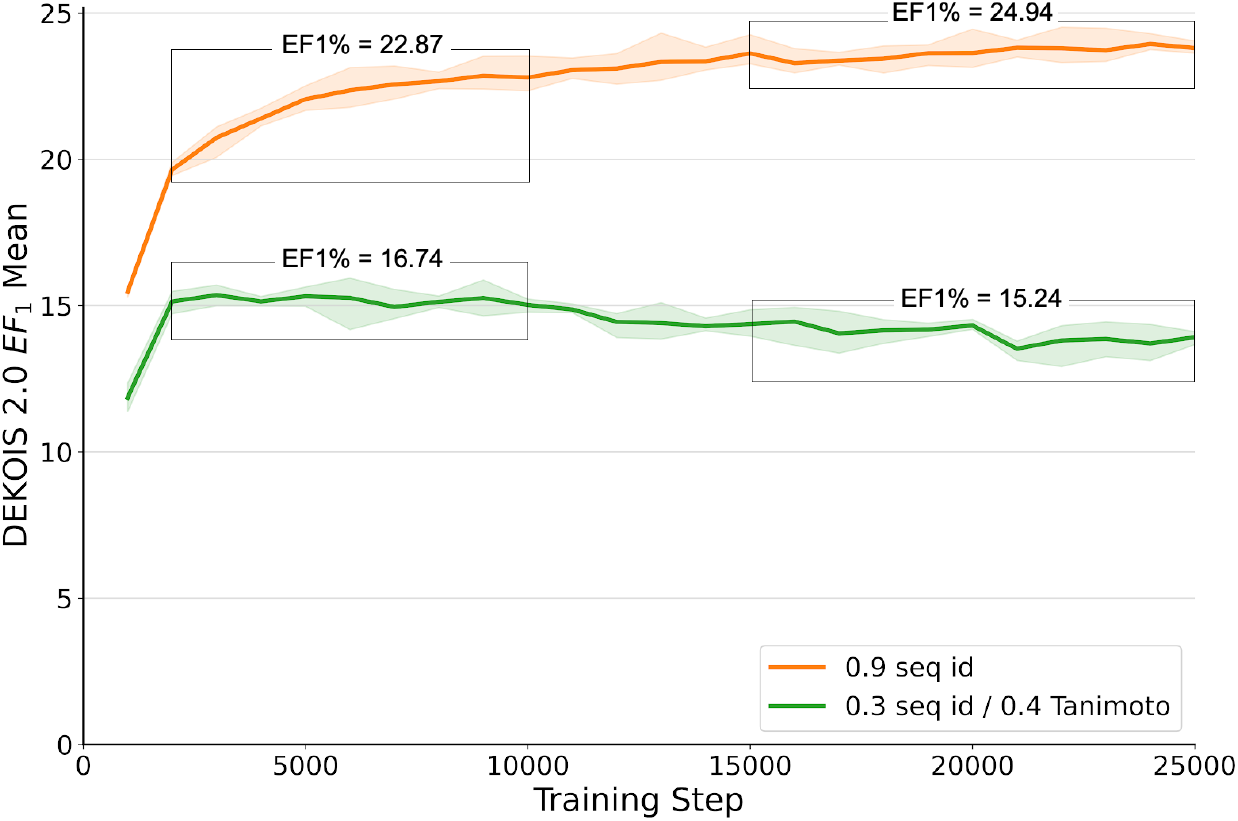
DEKOIS 2.0 performance based on the train-test split method and training steps. Our default train-test split method is denoted “0.3 seq id / 0.4 Tanimoto", while the protein-only split with 90% sequence identity cutoff is denoted “0.9 seq id". For each train-test split method we repeated the training with three different random seeds. The solid line reports the average DEKOIS 2.0 mean enrichment across the three runs, and the shaded area indicates the maximum and minimum mean enrichment scores from the three runs.

###### 2.1.2 Comparison with other methods

DOScore outperforms other docking-based methods on DEKOIS 2.0 benchmark (Table S8). To compare DOScore’s performance with recent state-of-the-art hit identification methods, we benchmarked it on DUD-E (*7*) and DEKOIS 2.0 sets using the protein 30% train-test similarity split from LigUnity’s paper (*8*). We compare the results with LigUnity and DrugCLIP (*9*) using the same split method, and show that our scoring model outperforms both methods on the benchmarks (Table S9).

**Table S8:** Hit identification performance of DOScore compared to other docking-based methods on DEKOIS 2.0 benchmark. DOScore is evaluated under our primary split protocol (30% Prot. / 0.4 Lig. (Sec. 5.1)). Enrichment factors (EF@1%) for other methods are obtained from prior work (*13*, *14*).

| Method | EF@1% |
| --- | --- |
| KarmaDock(FF) (14) | 16.29 |
| Glide SP (15) | 12.10 |
| Surflex-Dock (16) | 7.30 |
| LeDock (17) | 5.86 |
| Gold (18) | 5.36 |
| Vina (19) | 4.51 |
| TANKBind (20) | 2.90 |
| DOScore | 16.74 |

**Table S9:**
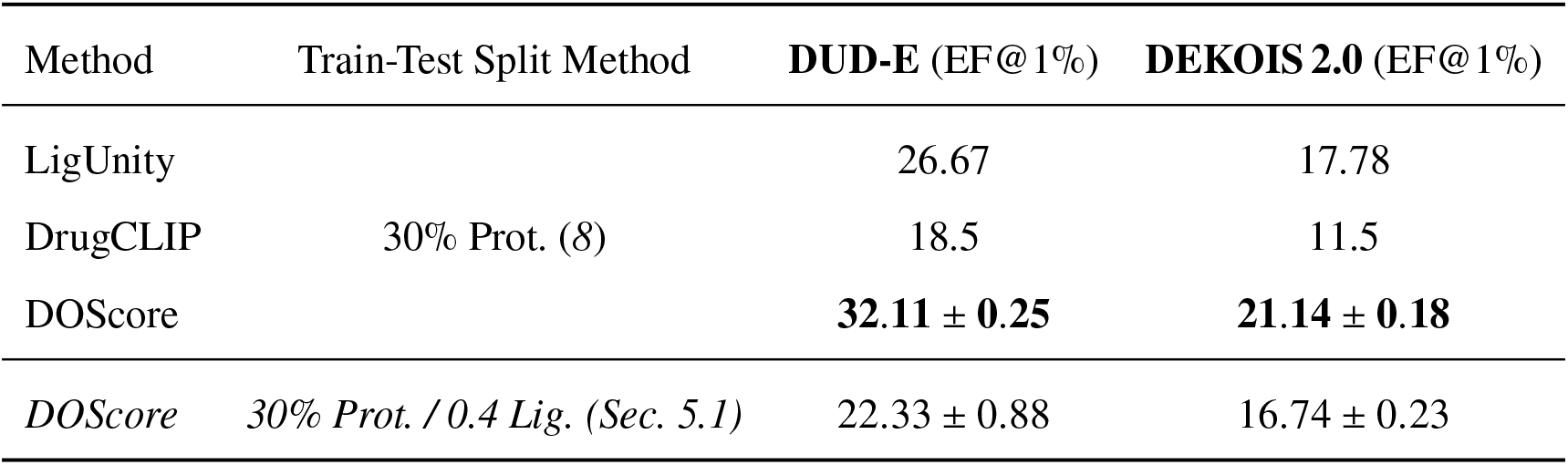
Hit identification performance of LigUnity (*8*), DrugCLIP (*9*), and DOScore on DUD-E and DEKOIS 2.0. Models are evaluated under the LigUnity protein 30% similarity split protocol for direct comparison, alongside DOScore performance under our primary split protocol. Enrichment factors (EF@1%) for LigUnity and DrugCLIP are reported from prior work (*8*), while DOScore results represent the mean ± std over three independent runs; best value per column in bold.

We additionally evaluate the fine-grained scoring variant of DOScore on several other scoring tests. On Merck (*10*) and JACS (*11*) benchmarks of the Schrodinger set (*12*), we again compare our results with LigUnity, using the same protein and ligand train-test split method (Table S10). Here, we report the correlation statistics between absolute binding affinites.

**Table S10:**
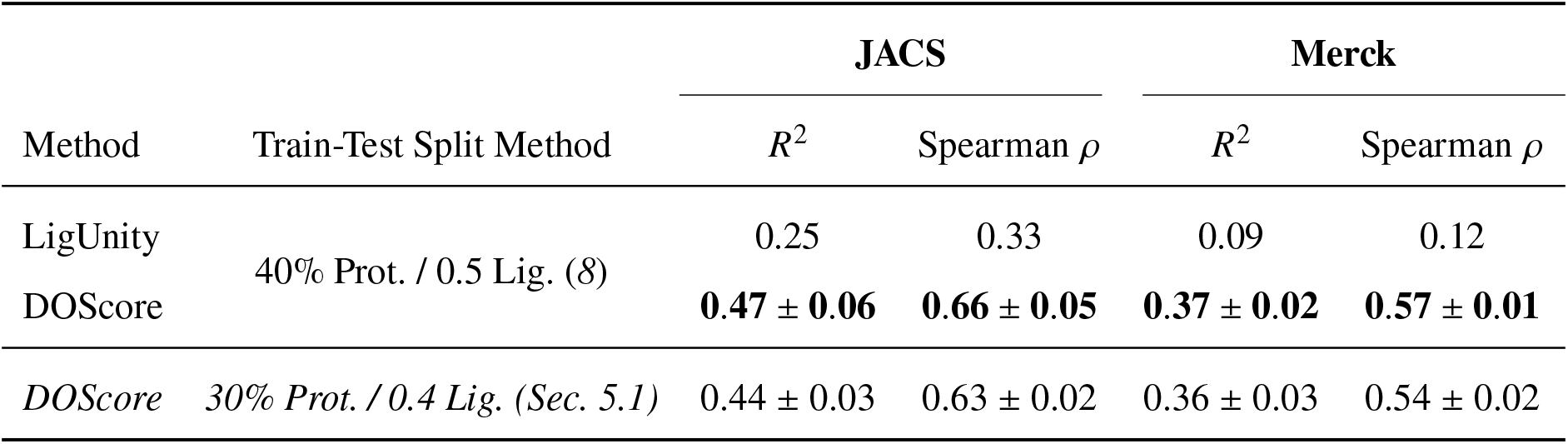
Affinity ranking comparison with LigUnity on JACS and Merck benchmarks. Models are evaluated under the LigUnity split protocol (40% protein and 0.5 ligand similarity cutoff) for direct comparison, alongside DOScore performance under our primary split protocol. *R*^2^ and Spearman *ρ* are calculated between predicted and experimental affinity values. LigUnity metrics are reported from the original paper, while DOScore results represent the mean ± std over three independent runs; best value per column in bold.

We also calculate and report the model’s performance on the recently released OpenBind affinity test set (Table S11). We applied the strict train-test split rules to the training set, removing all train complexes with proteins that had 30% or higher sequence identity to the test proteins and all complexes with ligands that had higher than 0.4 Tanimoto similarity to the benchmark’s ligands.

**Table S11:**
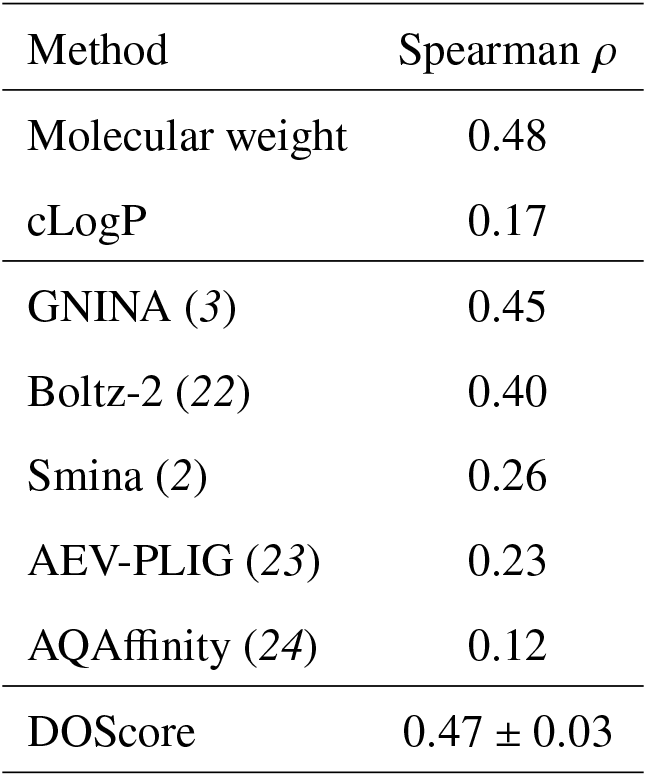
Affinity ranking comparison on the OpenBind benchmark. DOScore is evaluated under our primary split protocol (30% Prot. / 0.4 Lig. (Sec. 5.1)) and compared against various methods. Spearman *ρ* is calculated between predicted and experimental affinity values. Baseline metrics are obtained from the OpenBind blog post (*21*), while DOScore results represent the mean ± std over three independent runs.

##### 2.2 Ablation studies

To assess the contribution of DOScore design decisions (described in Section 8.2), we performed series of ablation studies using the DEKOIS2 data set for testing. We report average EF1% and EF5%, together with their standard deviations across three independent runs.

Evaluation results for the ablations are presented in Table S12.

**Table S12:**
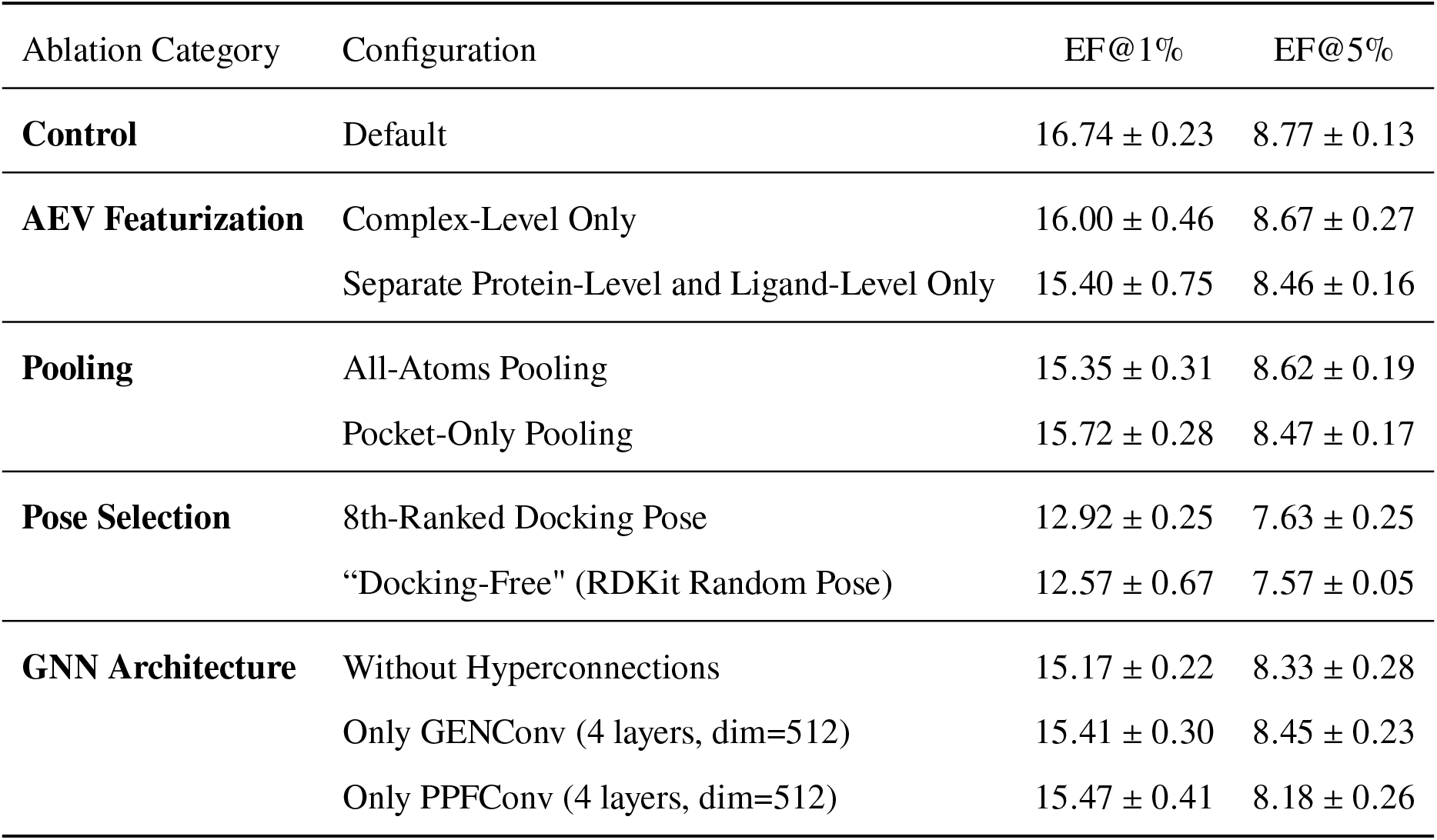
Results of ablation studies on the DEKOIS2 dataset. Values are reported as mean ± standard deviation over three independent runs.

###### AEV Features

To evaluate the importance of AEV representations, we investigate two architectural variants. In the first variant, we removed interface-specific features, only keeping the joint representation of the whole complex. In the second variant, the protein and ligand were encoded separately, eliminating cross-molecular interaction from the feature construction.

Both modifications resulted in a decrease in enrichment factors, demonstrating the importance of incorporating interaction-aware information at the feature level.

###### Readout module

Two alternative strategies for node pooling were evaluated. In the first strategy, pooling was performed over all nodes of the protein-ligand complex. The second strategy restricts pooling to the protein nodes only.

###### GNN module architecture

Three alternative architectures were considered for the GNN module design. In the first variant hyperconnections were removed from the GNN module to assess their impact on performance. In the second and third variants, the original dual-GNN architecture was replaced with a single GNN module based on either PPF or GEN. Both models consisted of four layers and had a hidden size of 512, matching the size of the original architecture.

###### Docking poses

To determine the impact of the docking pose on the scoring performance, we evaluated two alternative pose-selection strategies. In the first variant, during training and evaluation the topranked docking pose was replaced with the eighth-ranked pose generated by the docking procedure. In the second variant, we trained and evaluated a “docking-free" modification of the model entirely omitting the docking pose information.

For the “docking-free" variant of the model, for each protein-ligand complex we generated a structure where the ligand was represented by a randomly generated RDKit conformer and translated 100 Å away from the pocket. Because the ligand is placed far outside the pocket, all protein–ligand distances exceed the relevant thresholds: no protein–ligand edges are formed during graph construction, and cross protein–ligand features in the atomic featurizer collapse to zero. The transformer layer includes a learnable pairwise distance bias derived from interatomic distances, which could otherwise introduce artificial protein–ligand distance signals in this setting. To prevent this, we preserved the original distance-based biases for protein–protein and ligand–ligand atom pairs, while replacing all protein–ligand distance biases with a single learnable scalar shared across every protein–ligand pair. This design ensures that the “docking-free" model encoded protein and ligand representations without any information about their relative spatial arrangement.

#### 3 Experimental validation

##### 3.1 Clusters of experimentally validated hits

**Figure S6:**
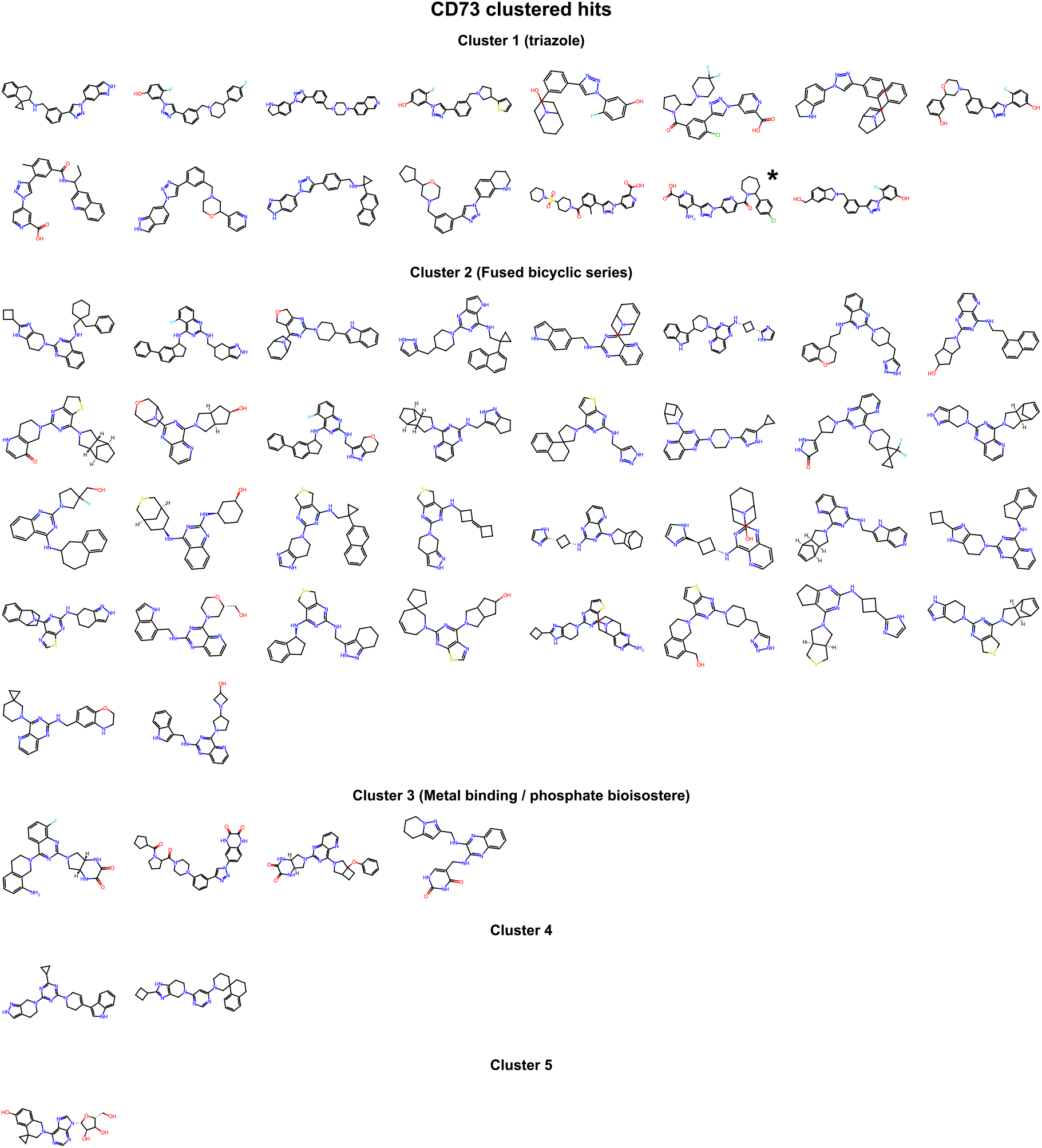
Clusters of CD73 experimentally validated hits. The molecule marked with a star exhibited 570 nM cellular activity.

**Figure S7:**
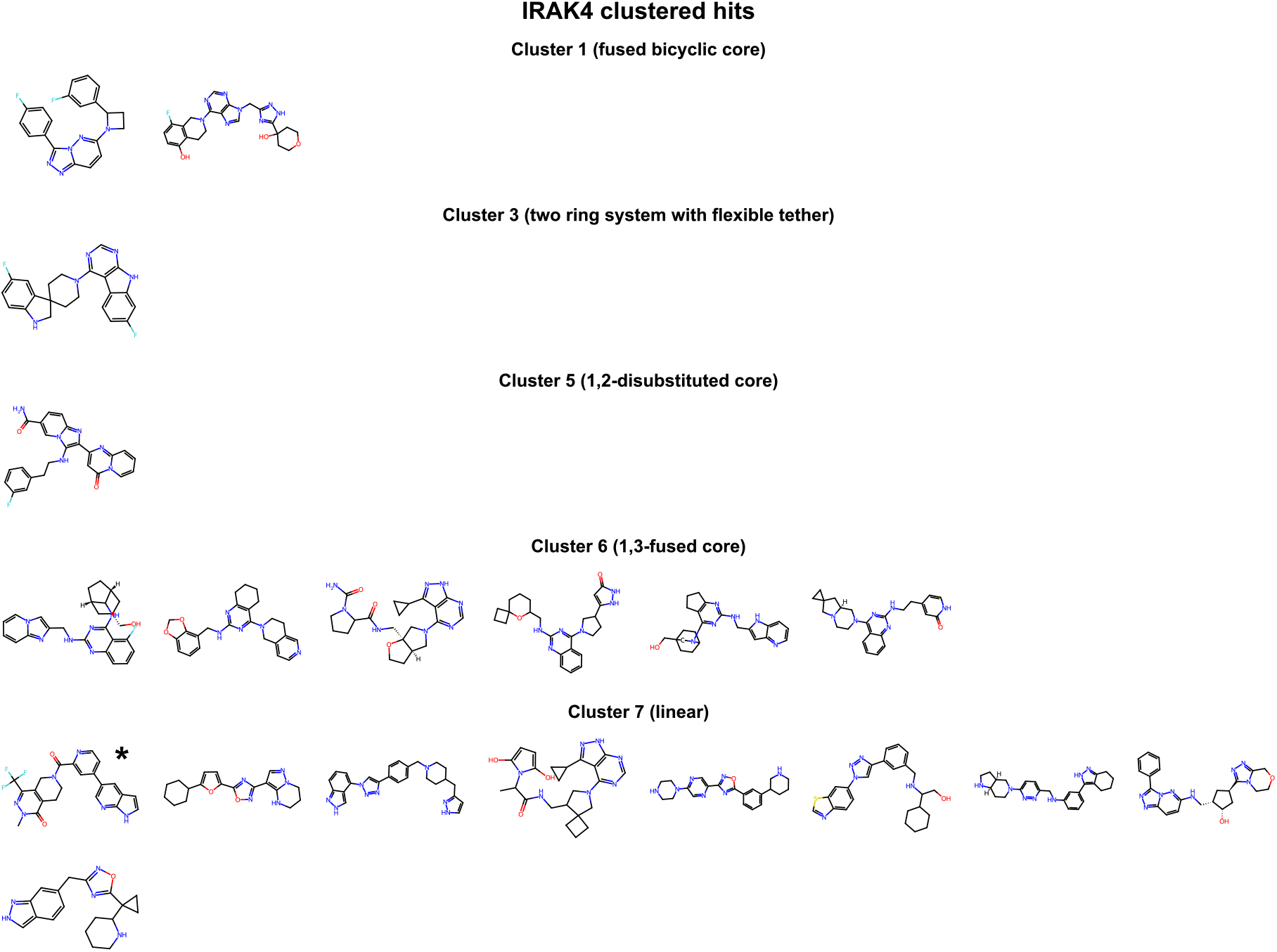
Clusters of IRAK4 experimentally validated hits. The molecule marked with a star exhibited 158 nM activity.

**Figure S8:**
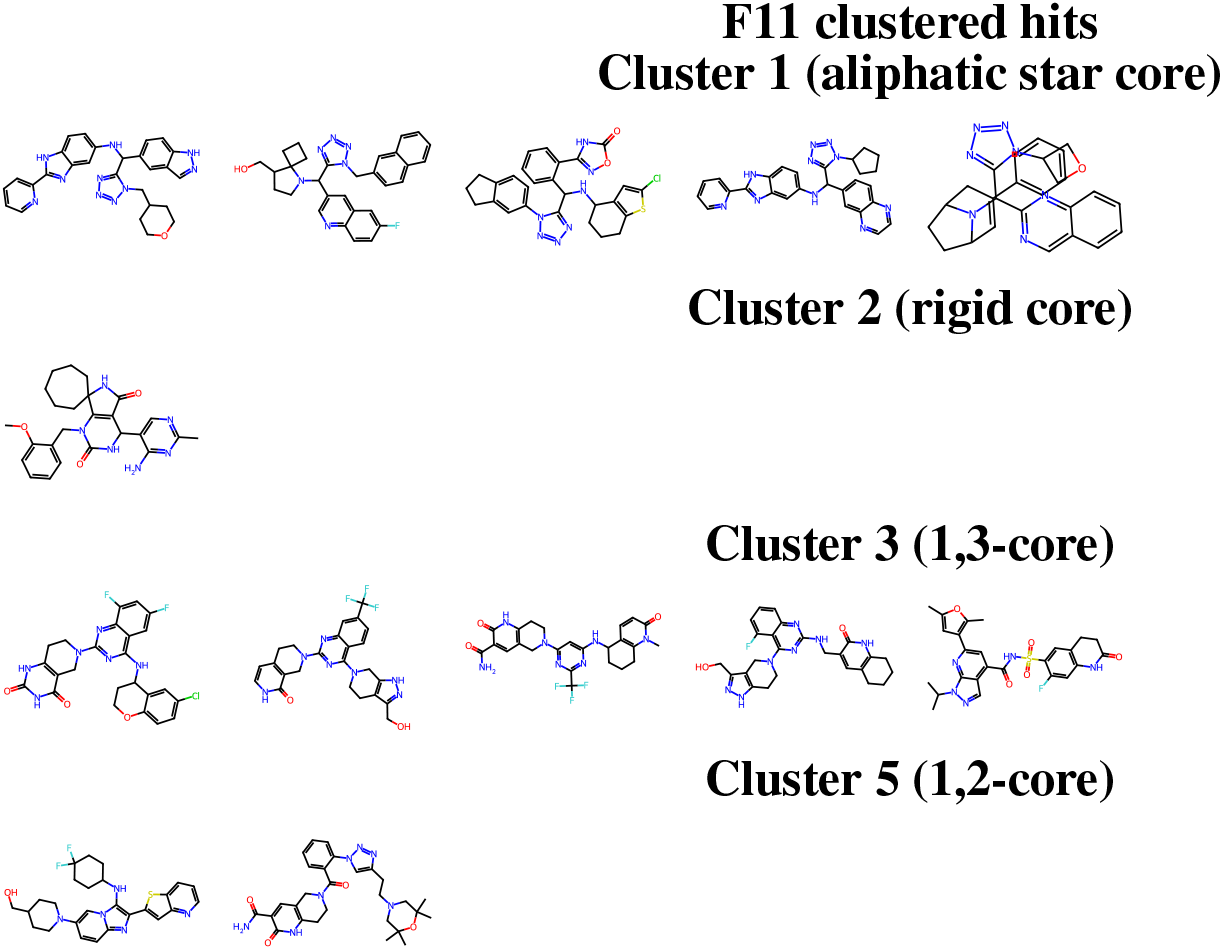
Clusters of Factor XI experimentally validated hits.

**Figure S9:**
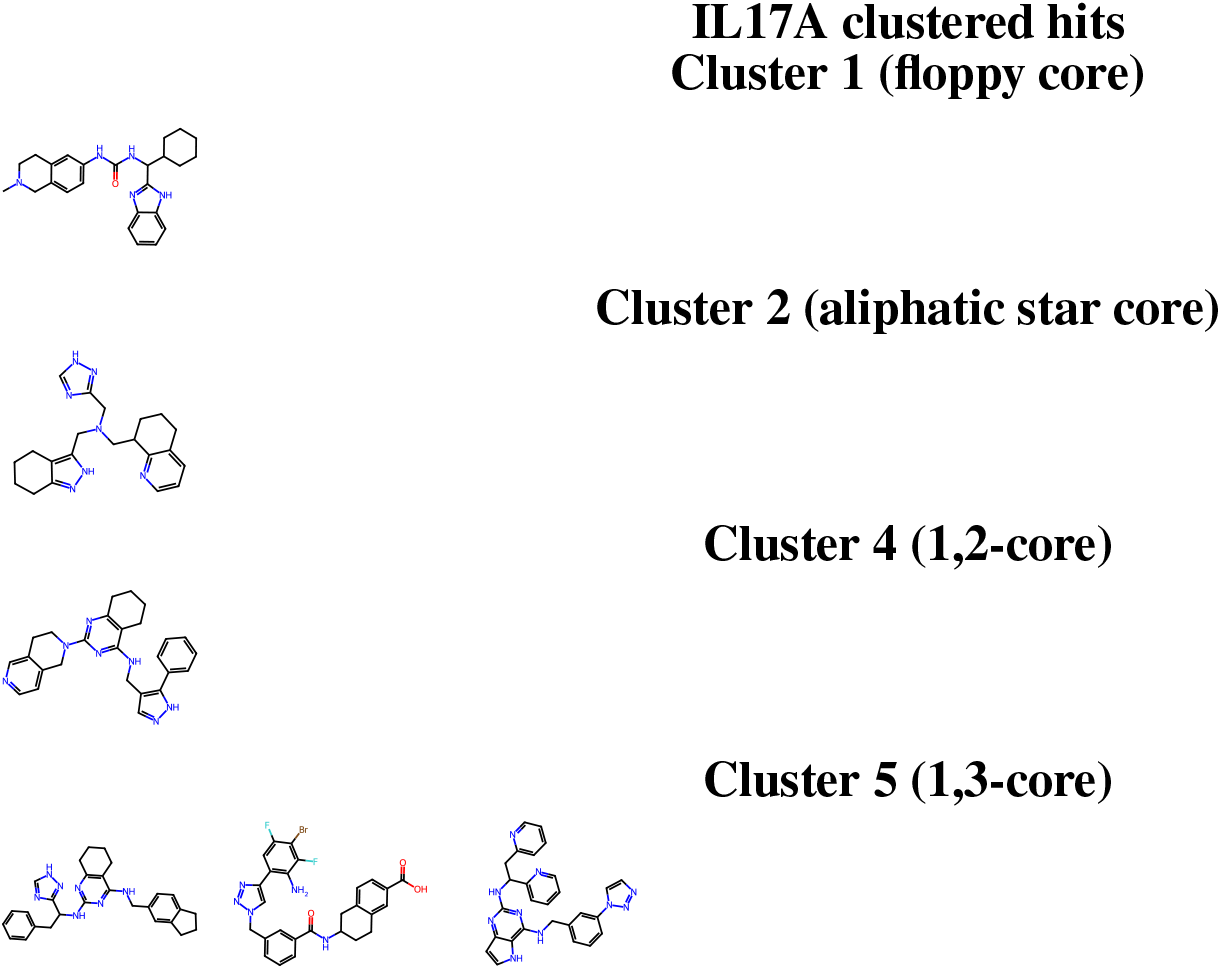
Clusters of IL17A experimentally validated hits.

### Materials and Methods

#### 4 Data sources and curation

We constructed separate but consistently processed datasets for training the docking and scoring methods, DODock and DOScore, respectively. The docking dataset was assembled from experimentally resolved protein–ligand complexes, where the bound ligand pose provides the structural reference for training and evaluation. The scoring dataset was assembled from assay-derived protein–ligand records, for which receptor structures were retrieved and processed independently.

##### 4.1 Docking training data sources

The docking dataset was constructed from both publicly available protein–ligand complex datasets, including PDBbind v2020 (*25*) and BioLiP2 (*26*) (July 2025), and an internally curated protein– ligand complex dataset derived from a 2025 snapshot of the PISCES pdbaa sequence database (*27*). For the PISCES-derived component, we first filtered pdbaa entries to retain X-ray structures with chain lengths between 40 and 10,000 residues, resolution ≤ 3.5 Å, and *R*-factor ≤ 0.25. The corresponding structures were then retrieved from the RCSB Protein Data Bank (*28*). Retrieved structures were processed using a multi-stage quality-control pipeline designed to remove entries with unsupported ligands, incomplete binding sites, ambiguous ligand definitions, or physically implausible protein–ligand geometries. The filtering stages are summarized in Table S13. After processing and filtering, the docking dataset contained 215,873 protein–ligand complexes.

**Table S13:** Filtering stages used during construction of the docking dataset.

| Stage | Filter | Complex Removal Criterion |
| --- | --- | --- |
| 1 | Undefined ligand identifiers | Ligand whose PDB chemical component identifier is UNK or UNX. |
| 2 | Unsupported ligand elements | Ligand containing any element other than H, C, N, O, P, S, F, Cl, Br or I. |
| 3 | Ligand heavy-atom count | Ligand with fewer than 8 or more than 100 heavy atoms. |
| 4 | Missing ligand heavy atoms | Ligand missing one or more heavy atoms relative to the corresponding ligand definition in the PDB Chemical Component Dictionary. |
| 5 | Steric clashes or invalid geometry | Nonphysical heavy-atom overlaps or invalid geometry involving the ligand or nearby protein residues. |
| 6 | Nearby non-cofactor ligands | Ligand with heavy-atom distance of 5 Å or less to other non-cofactor ligand. |
| 7 | Ligand buried surface area | Ligand buried solvent-accessible surface area ratio below 0.3. |
| 8 | Nearby non-standard residues | Ligand with heavy-atom distance of 8 Å or less to a non-standard protein residue. |
| 9 | Missing atoms in nearby residues | Ligand with heavy-atom distance of 5 Å or less to a protein residue with one or more missing heavy atoms. |
| 10 | Nearby missing-residue gaps | Ligand with a nearby gap, defined by 8 Å or less heavy-atom distance from the ligand to the resolved residue at the edge of the gap. |

###### Additional data preprocessing

For docking, it is important to ensure that experimentally resolved ligand poses are representable within the conformational space used by the docking pipeline. We therefore applied a reference-guided ligand optimization procedure within the docking transformation space defined in Section 7.1. Ligands were initialized from pipeline-compatible 3D conformations and optimized over rigid-body translation, rotation, and the allowed torsional angles to match the corresponding crystallographic ligand poses as closely as possible. In our pipeline, ligand 3D conformations are generated using RDKit ETKDGv3 (*29*), followed by MMFF94 (*30*) force-field minimization. This preprocessing procedure identifies complexes for which the crystallographic ligand pose can be reproduced within the ligand representation and transformation space used during docking. We applied the procedure to the full docking dataset and stored the optimized aligned ligand pose alongside the original crystallographic pose only when their RMSD was below 2 Å. Otherwise, the complex was retained with only the crystallographic ligand pose as the reference.

##### 4.2 Scoring training data sources

The scoring dataset was constructed from the PocketAffDB training data (*8*, *31*). PocketAffDB is a pocket-annotated assay dataset introduced in the LigUnity paper and provides assay-level proteinligand supervision covering 26,748 assays with 810,319 experimental measurements. It groups activity and affinity measurements, originally derived from public bioactivity resources such as BindingDB (*32*) and ChEMBL (*33*), into assays and associates them with binding-site annotations. The experimental bioactivity measurements included values such as *IC*_50_, *K*_*i*_, and *K*_*d*_.

Each assay contains ligands annotated with experimental bioactivity measurements, together with one or more pocket entries defined by a PDB identifier and protein chain identifier. These pocket entries reference structures in PocketAffDB and include co-crystallized ligands that define the relevant binding sites. Thus, the PocketAffDB records provide assay-level protein–ligand supervision together with ligand and binding-site metadata, but do not directly provide standardized receptor structures and ligand poses suitable for direct use as model inputs.

For the PocketAffDB-derived scoring data, we did not rely directly on the processed protein structures or protein chains distributed with PocketAffDB. Instead, we retrieved the corresponding PDB structures from RCSB Protein Data Bank (*28*) and processed the structures ourselves. This was done to avoid potential artifacts from the PocketAffDB processing pipeline and to retain complete protein structures, since PocketAffDB provides only selected protein chains for some entries and some provided chain selections may omit chains that are relevant to our model inputs. For each PDB identifier associated with an assay, we retrieved and processed the full protein structure from RCSB, while checking that the retrieved structures were consistent with the chains specified by the source records, the corresponding PocketAffDB pocket annotations, and the provided ligand identities. Assays for which none of the associated pocket structures could be processed successfully were excluded.

Following structure retrieval, validation, and filtering, we retained processed protein structures corresponding to 18,022 unique PDB IDs. The resulting dataset comprised 25,276 assays and 774,840 experimental measurements.

#### 5 Train-test splitting and benchmarking methodology

To assess model generalization, we constructed benchmark-specific training splits by applying explicit protein- and ligand-level similarity filters. These filters were applied prior to training to exclude candidate training complexes or interaction records that were closely related to examples in the corresponding benchmark set.

##### 5.1 Similarity filtering criteria

Protein similarity was assessed using Foldseek (*34*). For each candidate protein and each benchmark protein, we performed all-to-all chain comparisons by aligning every chain from the candidate protein against every chain from the benchmark protein. A candidate protein was excluded on the basis of protein similarity if at least one of its chains aligned to a benchmark chain with query coverage ≥ 0.5 and fractional sequence identity ≥ 30%. For the docking dataset, we additionally excluded candidate proteins for which at least one candidate chain aligned to a benchmark chain with query coverage ≥ 0.5 and lDDT ≥ 0.6. In both cases, the query was defined as the chain from the candidate training protein. This criterion was intended to remove close sequence homologs as well as structurally related proteins, including cases where substantial structural similarity remains despite limited sequence identity.

Ligand similarity was evaluated using the Tanimoto coefficient between 2048-bit radius-2 Morgan fingerprints (ECFP4 (*35*)), generated using RDKit (*4*). A candidate training ligand was excluded if its similarity to any ligand in the corresponding benchmark set exceeded 0.4.

Candidate examples satisfying either the protein-level or ligand-level similarity criterion were removed from the corresponding training split.

The resulting splits reduce overlap between the training and benchmark data in both protein target space and ligand chemical space. Consequently, benchmark performance more directly reflects generalization to new protein–ligand systems, rather than performance driven by highly similar proteins, ligands, or complexes observed during training. Unless otherwise stated, all docking and scoring experiments use the processed datasets and similarity-controlled splits described in this section.

##### 5.2 Model selection

To ensure a consistent comparison across training runs, for both docking and scoring the model checkpoints were selected using a fixed training budget rather than benchmark-specific validation performance. For all experiments the diffusion model checkpoint was selected after a fixed number of training steps, while the ranking model checkpoint was selected after a fixed number of complete passes over the corresponding training dataset. Additional details on the training procedures for DODock’s diffusion and ranking models are provided in Sections 7.2, 7.5, respectively. For DOScore, the model selection is detailed in Section 8.3.

##### 5.3 Benchmarks and evaluation metrics

###### 5.3.1 Docking

We evaluated docking performance on several protein–ligand docking benchmarks: the 285-complex CASF-2016 set under our targeted-redocking protocol (hereafter, CASF-2016 (redocking)) (*36*), the 308-complex PoseBusters Benchmark set (*37*), Runs N’ Poses (*38*), and OpenBind (*21*). Unless otherwise stated below, benchmark structures and metadata were used as provided by the corresponding authors without manual modification. Any restrictions or additional filtering applied to an evaluation subset are described in the corresponding benchmark section.

The primary docking accuracy criterion was heavy-atom ligand RMSD between the predicted and experimentally resolved ligand pose. RMSD was computed with symmetry correction, so that chemically equivalent atom permutations were treated as equivalent. For benchmark sets where physical and chemical validity annotations were available or could be computed, we additionally report PoseBusters validity (*37*).

For OpenBind and Runs N’ Poses, we also report accuracy under the additional criterion LDDT-PLI ≥ 0.8, following the corresponding benchmark protocols. This criterion measures recovery of the local protein–ligand interaction environment. In analyses using this criterion, a prediction was counted as successful only if it satisfied the RMSD requirement and the LDDT-PLI threshold; for OpenBind, the combined criterion additionally requires PoseBusters validity.

###### CASF-2016 (redocking)

We use the 285 protein–ligand complexes of the CASF-2016 benchmark set (*36*) for an end-to-end targeted-redocking evaluation. This evaluation is distinct from the official CASF-2016 docking-power protocol, which assesses scoring functions by ranking predefined decoy poses. For this evaluation, the docking models were trained on the corresponding similarity-filtered split, obtained by applying the protein- and ligand-level filters described in Section 5 with respect to all benchmark complexes. We report the percentage of complexes for which the top-ranked predicted pose has a symmetry-corrected ligand heavy-atom RMSD of at most 2 Å or 1 Å from the crystallographic pose (Fig. 1a).

###### Runs N’ Poses

Runs N’ Poses is a benchmark designed to evaluate generalization of protein– ligand co-folding and docking methods across different levels of similarity to the training set (*38*). For this evaluation, the docking model was trained on the time-split training complexes provided by the Runs N’ Poses benchmark, using the 2021 cutoff defined in the corresponding study. We used the similarity bins provided by the benchmark and report the performance separately for each bin. Following the benchmark protocol, a prediction was considered successful only if it satisfied both criteria: symmetry-corrected heavy-atom RMSD < 2 Å and LDDT-PLI > 0.8. This stratified evaluation allows performance to be assessed as a function of increasing similarity with training data.

For cross-method comparison (Fig. 1b), we used a common-subset evaluation. Specifically, we restricted the Runs N’ Poses analysis to benchmark complexes for which all four competing methods produced results and for which DODock also completed successfully. This yielded an initial common subset of 2,286 complexes. We further removed complexes containing severe steric clashes between ligands annotated as proper benchmark ligands. A clash was defined as a distance of less than 3 Å between any pair of heavy atoms belonging to different proper ligands. After this additional filtering step, the final evaluation subset contained 2,149 complexes.

###### PoseBusters

We use the 308-complex PoseBusters Benchmark set (*37*) to evaluate both docking accuracy and the physical plausibility of predicted protein–ligand complexes. For this benchmark, we consider two training settings. In the first setting, the docking model was trained on the corresponding similarity-filtered split, obtained by applying the protein- and ligand-level filters described in Section 5 with respect to all PoseBusters complexes. In the second setting, the model was trained on a time-split constructed from PDBbind v2020 complexes, reflecting the training setup commonly used by other methods evaluated on PoseBusters.

Results for the two settings are reported separately. For comparison with published methods, we use the PDBbind v2020 time-split model and report the strict combined success criterion, requiring both symmetry-corrected heavy-atom RMSD < 2 Å and PoseBusters validity (Figure S1). For the similarity-filtered model, we report success rates stratified by protein family using RMSD thresholds of 2 Å and 1 Å (Fig. 1e).

###### OpenBind

OpenBind evaluates docking and co-folding methods on recently released protein– ligand structures with associated structure–affinity data (*21*). For this benchmark, the docking model was trained on the corresponding benchmark-specific similarity-filtered split described in Section 5. We used the benchmark inputs and reference poses as provided. Following the OpenBind evaluation protocol, we excluded reference complexes classified as suspected crystal-contact artifacts, fragment-binder screening cases, or non-PoseBusters-valid. The resulting evaluation set comprised 802 follow-on compound complexes.

We report PB-valid RMSD success, requiring both symmetry-corrected heavy-atom RMSD ≤ 2 Å and PoseBusters validity. We additionally report the stricter combined criterion requiring RMSD ≤ 2 Å, PoseBusters validity, and LDDT-PLI ≥ 0.8 (Fig. 1d). This combined metric evaluates whether a method recovers an accurate ligand pose while also producing a physically valid complex with a well-recovered local interaction pattern.

###### 5.3.2 Scoring

We evaluated DOScore on hit identification benchmarks DUD-E (*7*), DEKOIS 2.0(*39*), and scoring benchmarks OpenBind (*21*), Merck (*10*), and JACS (*11*). For each benchmark, we removed from the training data all complexes containing proteins or ligands similar to those in the test set, using the train–test similarity filtering criteria described in Section 5.1 unless specified otherwise. The model was trained from scratch on the resulting split to prevent train–test leakage, and the benchmark set was not used for validation. We repeated the training three times for each benchmark and report the average performance.

Performance was evaluated using the docked structures of the benchmark set’s protein-ligand complexes. For each target, we used the protein files provided by the respective benchmark. For ligands, we used the SMILES strings supplied by the benchmark where available; when a benchmark instead provided ligand structures in SDF format, we extracted the corresponding SMILES from those structures using RDKit. These ligands were then docked into the provided protein structures for evaluation.

For DUD-E and DEKOIS 2.0, we evaluated hit identification performance using the enrichment factor (EF), which measures how strongly true actives are concentrated among the top-ranked compounds relative to random selection. For a given cutoff *x*, EF is defined in Eq. S1.

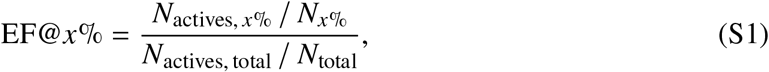

where *N*_*x*%_ is the number of compounds in the top *x*% of the model prediction-ranked list, *N*_actives, *x*%_ is the number of actives among them, and *N*_actives, total_ and *N*_total_ are the total numbers of actives and compounds, respectively. An EF value of 1 corresponds to random performance, while higher values indicate greater early enrichment of actives. We primarily report EF@1% (and additionally report EF@5% for the model ablation experiments).

For OpenBind, Merck, and JACS benchmarks, we calculate Spearman correlation between the predicted binding scores and the experimental affinities.

The benchmarking results are reported in the Supplementary Text (Section 2.1.2).

#### 6 Protein-ligand interaction modeling components

We employ neural network modeling for protein-ligand complexes across three distinct tasks: twice within the DODock docking pipeline (first as a diffusion-based denoiser and second as a ligand pose ranking model) and as the standalone DOScore model for estimating binding affinity. These three models are built upon a shared architectural framework that represents the protein-ligand complex as an atom-level graph processed by graph neural networks (GNNs) and attention mechanisms, with modifications tailored to each specific task.

##### 6.1 Graph formulation

The ligand and protein pocket are represented as a molecular graph G = (V, E), where nodes *v*_*i*_ ∈ V correspond to atoms and edges correspond to atom pairs within a specified distance threshold. The specific pocket atom selection and edge construction thresholds vary by task (Section 7.5.2 for the pose ranking model, Section 7.2.2 for the diffusion denoiser, and Section 8.2 for DOScore).

##### 6.2 AEV featurization

The protein-ligand graph’s nodes are featurized using the roto-translation invariant Atomic Environment Vector (AEV) representation introduced in the ANI neural network potentials (*40*), retaining the standard AEV form with the parameter set reported in Table S14. Here, the AEV featurizer is utilized twice: once for the full interaction features of the entire complex (AEV_full_), and once on the ligand and protein pocket separately (AEV_separate_). The subtraction of the full and separate embeddings isolates the ligand-protein interface interactions:

**Table S14:**
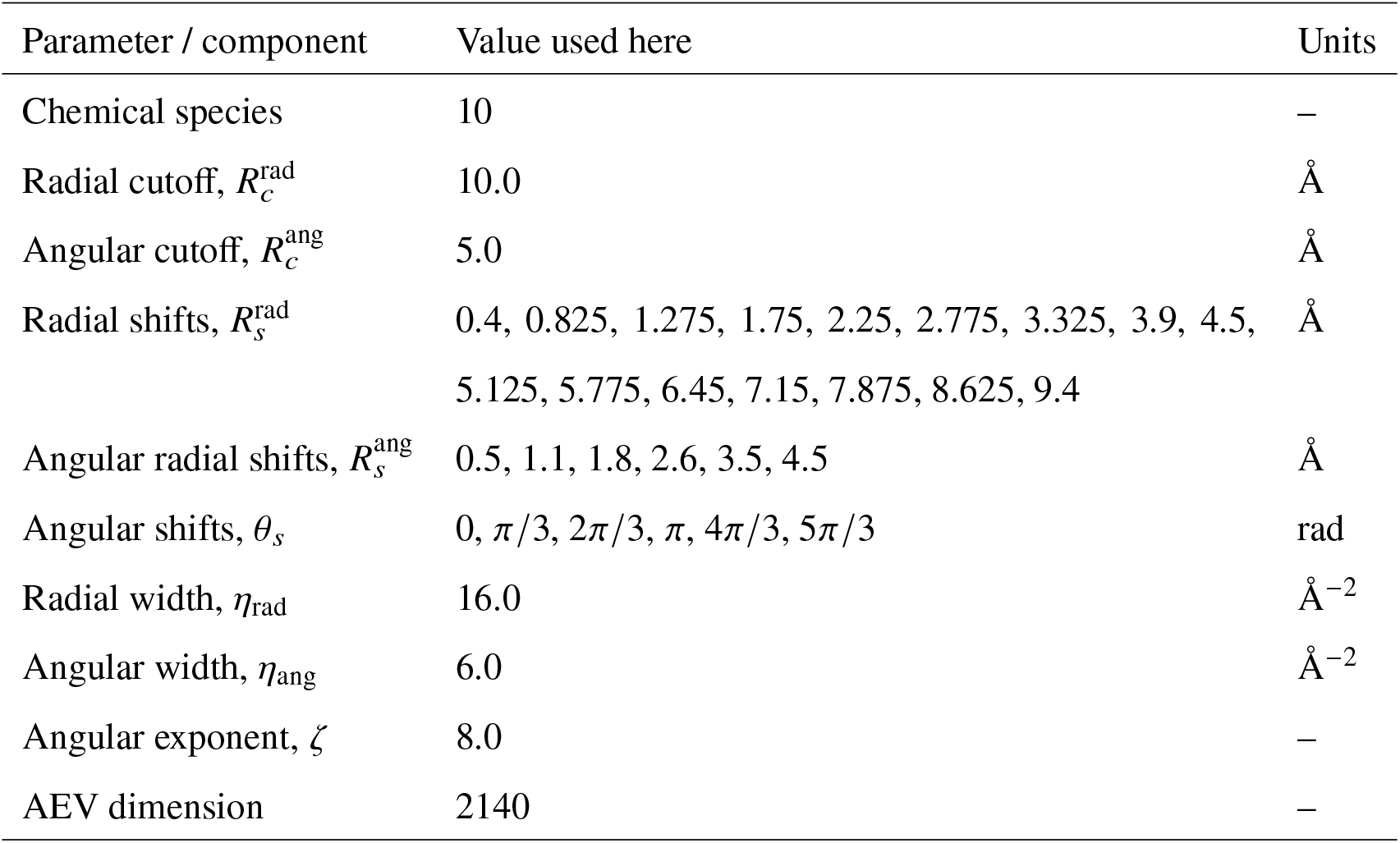
AEV featurizer parameters used in this work.

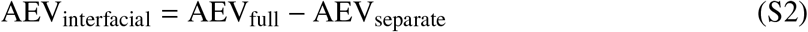

We concatenate AEV_full_ and AEV_interfacial_ to obtain the final AEV represention of each node. Then, an atom-type based MLP maps these concatenated embeddings into a smaller dimension space (*d*_model_) followed by layer normalization to define the initial node features ℎ^(0)^.

##### 6.3 Graph Neural Network operators

The neural networks in DODock’s denoiser and DOScore use PPFConv (*41*) and GENConv (*42*) GNN models, as described below.

###### Point-Pair Feature Convolution (PPFConv)

The PPFConv branch incorporates explicit geometric information through pairwise distance and orientation descriptors. The edge features use a modified PPF setup where the central node’s surface norm is replaced with the average vector of the position differences with neighboring nodes (Eq. S3).

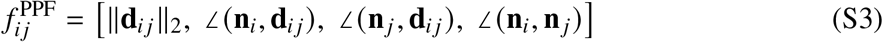

where **d**_*i* *j*_ = **p** _*j*_ − **p**_*i*_ is the relative position vector. This branch uses local and global MLP operators to process the geometric features.

###### Generalized Graph Convolution (GENConv)

The GENConv branch focuses on adaptive neighborhood aggregation to provide propagation of structural and chemical information across the graph. It uses a multi-layer MLP operator to aggregate features while mitigating feature over-smoothing.

When used together, the PPFConv branch and GENConv branch process the same graph in parallel, and their resulting node representations are combined through concatenation.

##### 6.4 Transformer with geometric attention bias

Global information exchange and readout are handled by multi-head attention and transformer modules (*43*). The transformer encoder architecture used in DODock’s denoiser and DOScore processes the GNN output node embeddings using a self-attention mechanism that incorporates pairwise structural biases directly into the attention score computation.

Spatial relationships are captured by utilizing the Euclidean distance *d*_*i* *j*_ between the 3D coordinates of the atoms. This distance is expanded using Radial Basis Functions (RBF) and processed through an MLP to generate a head-specific geometric bias *b*_*i* *j*_ (Eq. S4).

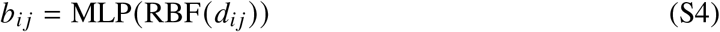

This bias is added directly inside the attention score calculation (Eq. S5).

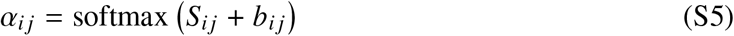

The output representation of each atom is calculated as the weighted sum of the value vectors.

#### 7 Docking model

DODock takes as input a ligand SMILES string, a protein structure, and a specification of the target binding pocket, and searches the docking transformation space defined in Section 7.1, generating a set of candidate binding poses from which it returns the top-ranked pose. As illustrated in Fig. 1c, the method comprises three principal stages: (i) diffusion-based pose generation, described in Section 7.2; (ii) post-diffusion refinement using the global and local optimization methods in Section 7.4, guided by the DOFast scoring function introduced in Section 7.3; and (iii) pose ranking using the learned models presented in Section 7.5. Section 7.6 combines the components into the complete docking pipeline and specifies the standard inference settings. Finally, Section 7.7 describes the extension of the method to blind docking.

##### 7.1 Search space

The argument space of the search is modeled as a transformation space of rigid roto-translation transformations combined with torsional angle rotations, conditioned on a ligand start conformation. The pivot of the global rotation is the ligand’s designated origin atom **L**[*o*], which remains stationary under rotation alone, keeping translation, rotation, and torsion decoupled. While the rotation transformation is defined as an element of *SO*(3) and applied to the ligand with the origin atom at the origin of the coordinate system, different algorithms use various representations of the rotation, including quaternions and Euler angles, and utilize conversions between them. Torsion-bond rotations are applied sequentially in a fixed order: each bond reads its pivot axis from the *already-updated* coordinates, implementing standard tree-of-torsions semantics in which downstream bonds inherit upstream rotations. Each torsion update rigidly rotates the molecular sub-tree downstream of the corresponding bond, thereby preserving the bond lengths, bond angles, and rigid fragments of the input ligand conformation. Once the pivot axis of a torsion is determined, the only transformation parameter is its angle of rotation. Hence, for a molecule with *d* torsional angles, the torsional transformation subspace is *SO*(2)^*d*^, and the overall transformation space is

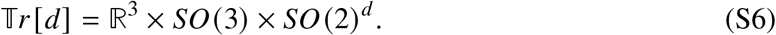

Algorithm 1 describes the procedure for applying a transformation to an input ligand conformation.

###### Algorithm 1

ApplyTransformation

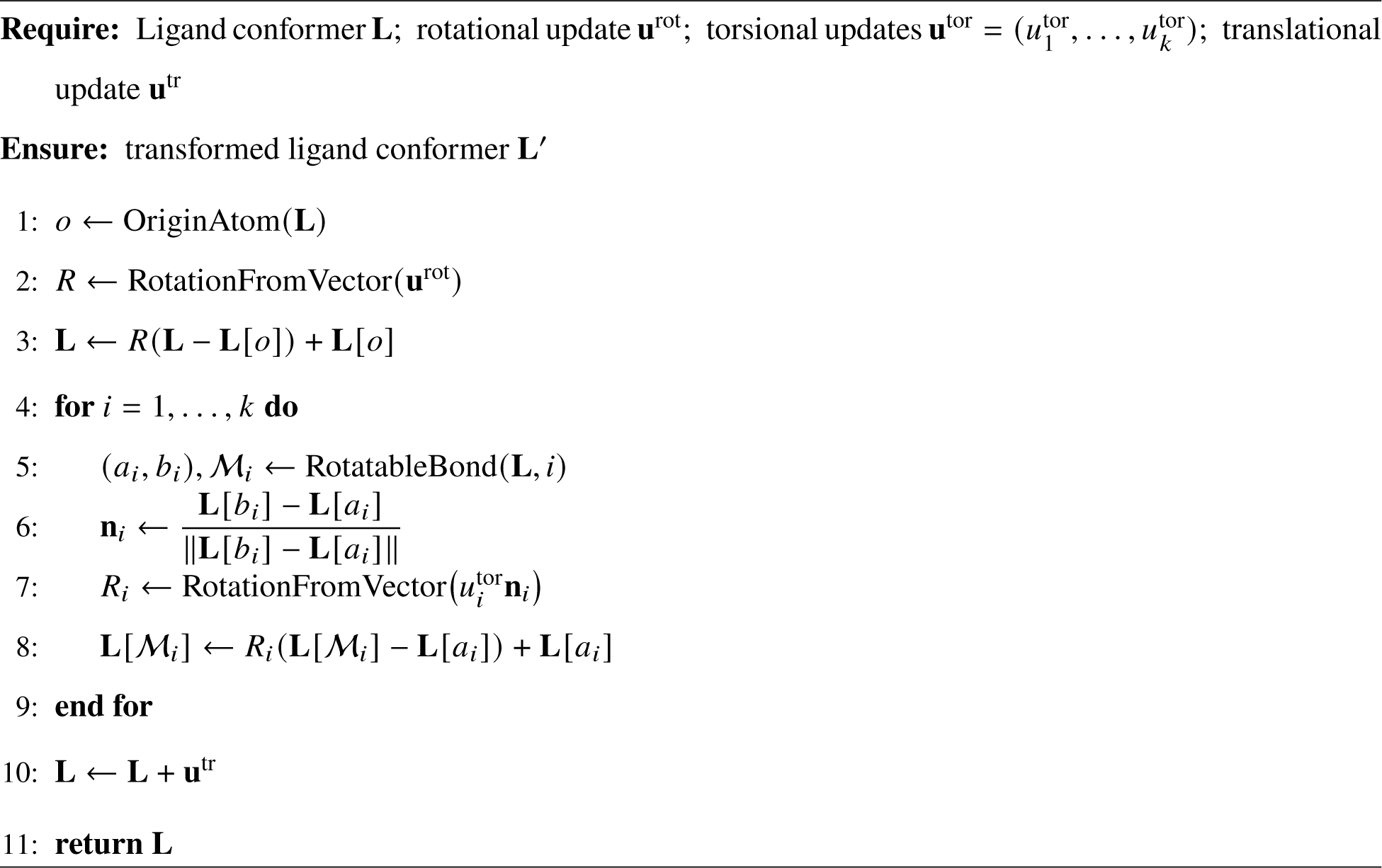

##### 7.2 Diffusion docking model

The diffusion part of the docking was motivated by DiffDock (*44*). The docking pose is parameterized by a translation vector *μ* ∈ ℝ^3^, a rotation matrix *R* ∈ *SO*(3), and a vector of torsion angles *τ* ∈ T^*d*^ *SO*(2)^*d*^ and its initial configuration *X*_0_, where *d* denotes the number of rotatable bonds. The corresponding initial poses (without added noise) are parameterized by *μ*_0_, *R*_0_, *τ*_0_ (which, in our setting, are all taken to be zero), and by initial configuration *X*_0_. Throughout, the receptor is modeled as a rigid body, as detailed in Section 7.1. The diffusion time **t** was represented as a three-dimensional vector (*t*_tr_, *t*_rot_, *t*_tor_) (where *t*_tr_, *t*_rot_, and *t*_tor_ denote the diffusion times for the translational, rotational, and torsional degrees of freedom (DOFs), respectively), and the corresponding standard deviations were denoted by (*σ*_tr_, *σ*_rot_, *σ*_tor_). The relationship between these standard deviations and the diffusion times is detailed in Section 7.2.3. An overview of the denoising cycle and docking model is shown in Figure S10.

###### 7.2.1 Pocket definition

The binding pocket is defined as all protein residues containing at least one heavy atom within 8.0 Å of any heavy atom of the aligned ligand (Section 4.1). For blind docking, the pocket is defined using a semi-optimal ligand pose rather than the aligned ligand pose. This pose is obtained as part of the blind-docking workflow described in Section 7.7.

###### 7.2.2 Pose space and graph construction

The ligand and protein pocket are represented as a molecular graph G = (V, E), where nodes *v*_*i*_∈ V correspond to atoms. Edges (*i*, *j*) ∈ E are formed between atom pairs within a type-dependent distance cutoff:

- **Ligand–ligand:** 4.0 Å
- **Protein–protein:** 8.0 Å
- **Ligand–protein:** 8 + 3 · min(*σ*_tr_, 4.0) Å

At high noise levels (*t*_tr_ ≈ 1), the ligand may be displaced far from the pocket, so a larger cutoff ensures sufficient protein–ligand connectivity which gives the model an opportunity to understand the direction to the protein. The graph is recomputed at every denoising step during inference based on the current ligand coordinates.

**Figure S10:**
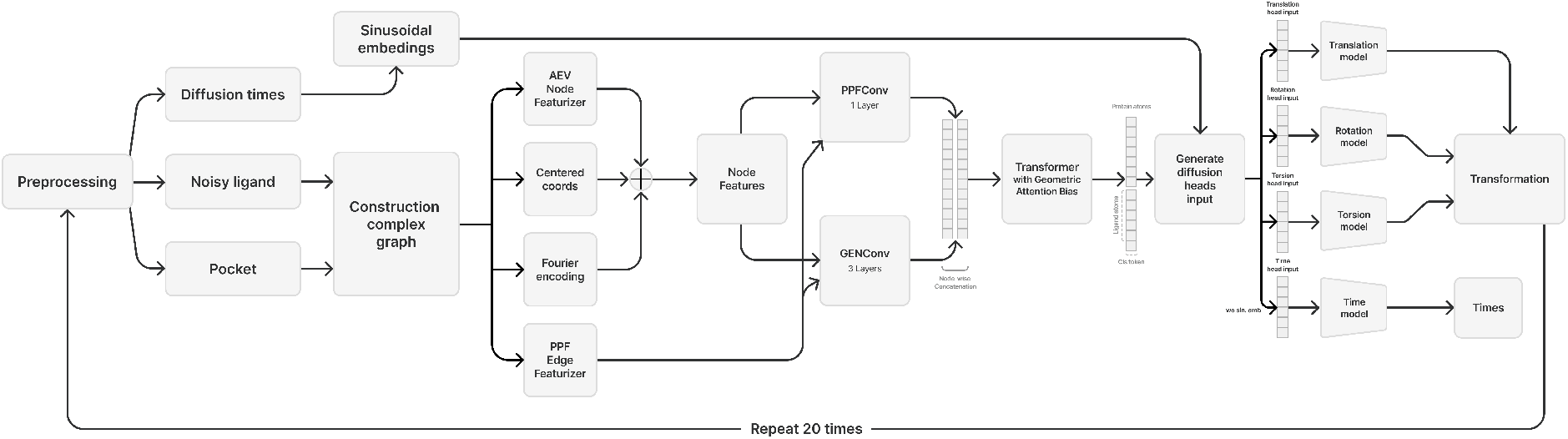
Denoising cycle of the reverse diffusion sampler. The noisy ligand pose is iteratively updated at each denoising step *t*_*i*_ using translation, rotation, and torsion scores predicted by the model to yield the final docked structure. Predicted diffusion times were used only for training.

###### 7.2.3 Forward diffusion process

We define the forward process for each degree of freedom (DOF) independently as a variance-exploding SDE (*45*):

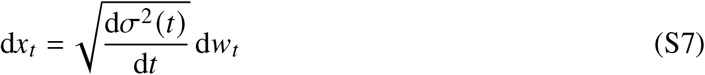

where *x*_*t*_ denotes the state of the corresponding DOF at diffusion time *t*, *w*_*t*_ is a Wiener process, and *σ*(*t*) is the noise-scale schedule controlling the amount of perturbation added at time *t*. This defines a family of corrupted pose distributions *q*(*x*_*t*_ | *x*_0_), where each DOF is perturbed independently.

###### Translation

The translational diffusion kernel is a Gaussian:

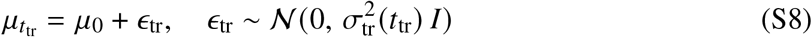

where *μ*_0_ denotes initial translation, 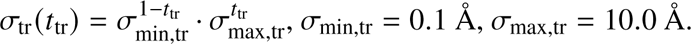

###### Rotation

The diffusion kernel on *SO*(3) is the IGSO(3) distribution (*46*, *47*), sampled in axis-angle parameterization with angle density:

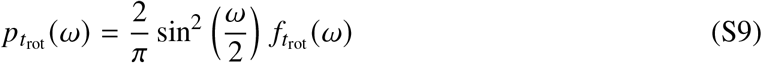

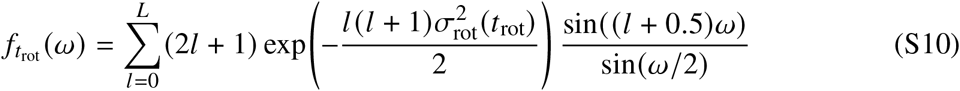

where 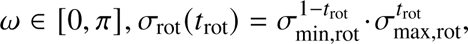 *L* = 2000, *σ*_min,rot_ = 0.1 rad, *σ*_max,rot_ = 1.65 rad.

A rotation is sampled by first drawing an angle *ω* from *p*_*t*_rot__ (*ω*) via inverse-CDF sampling on a precomputed grid, then pairing it with a uniformly random axis *n̂* ∈ *S*^2^ to form the axis-angle perturbation *ωn̂*, which is composed with the current rotation.

###### Torsion

Each rotatable bond angle is perturbed independently using a wrapped Gaussian on T (*48*):

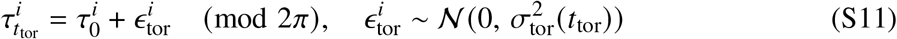

where *τ*_0_ denotes initial torsion, 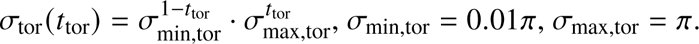

###### 7.2.4 Training dataset and optimization setup

Training examples were obtained from the docking dataset described in Section 4.1. Each model was trained for 300,000 optimization steps, and a checkpoint was saved for each of the last 7 epochs. The final model was obtained by averaging the parameters of the saved checkpoints.

We used the Adam optimizer (*49*) with a learning rate of 10^−4^, and a batch size of 64. The diffusion times for translation, rotation, and torsion were sampled independently from a uniform distribution on [0, 1]. We denote the resulting time vector by

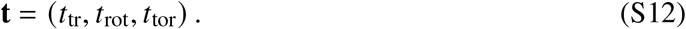

The model was trained using denoising score matching (*50*), which extends the score-matching framework of Hyvärinen (*51*) to noise-perturbed data. We also adopt the weighted score-matching formulation of Song et al (*45*).

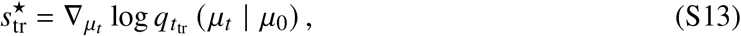

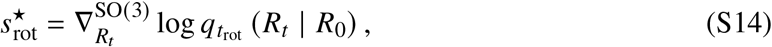

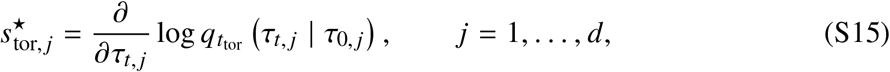

where *d* is the number of rotatable bonds and *q*_*t*_denotes the corresponding forward perturbation kernel, *μ*_0_, *R*_0_, *τ*_0_ are the initial transformations (in our case they are zero) and *μ*_*t*_, *R*_*t*_, *τ*_*t*_ are noisy transformations. The ground-truth rotational score target 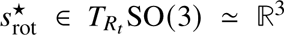 is computed analytically under the IGSO(3) distribution using axis-angle parameterization (*47*):

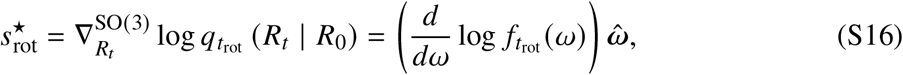

where *ω* ∈ [0, *π*] is the rotation angle between *R*_0_ and *R*_*t*_, *ω̂* ∈ ℝ^3^ is the unit rotation axis. The three denoising score-matching losses are

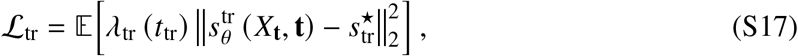

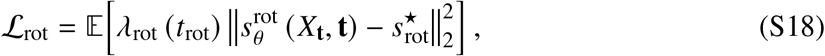

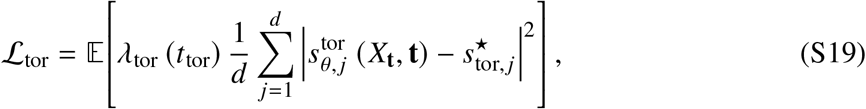

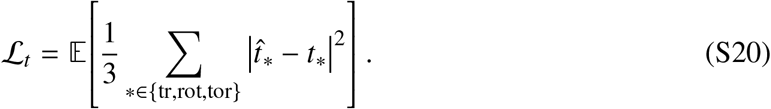

For ligands without rotatable bonds, i.e., *d* = 0, the torsional loss is defined to be zero.

In these expressions, the expectation is taken over aligned ligand conformation *X*_0_ sampled from the training dataset, independently sampled diffusion times **t**, and perturbations sampled from the corresponding forward diffusion kernels:

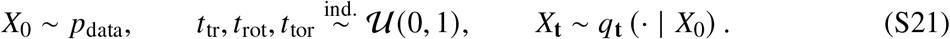

Following (*45*), the noise-dependent weights are chosen as the inverse expected squared norm of the corresponding conditional score:

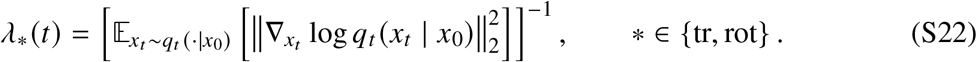

In this definition, the mathematical expectation is taken with respect to the forward perturbation at a fixed noise level *t*. Owing to the homogeneity of the perturbation kernels, the resulting normalization depends on the noise level but not on the particular clean translation or rotation.

For torsion, normalization is defined for a single torsional component:

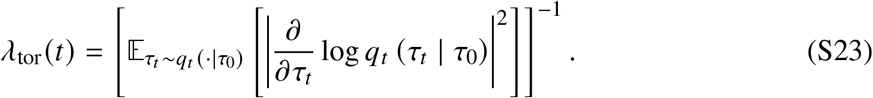

The same noise-dependent weight is applied separately to every rotatable-bond component, after which the component losses are averaged over the *d* rotatable bonds. This weighting balances the contributions from different noise levels, whose unnormalized target-score magnitudes may differ substantially.

The analytical targets are computed using the Gaussian conditional score for translation, the isotropic Gaussian distribution on SO(3), denoted IGSO(3), for rotation (*47*), and the wrapped Gaussian conditional score for torsion (*48*).

The total training objective combines the three score-matching losses with the auxiliary diffusion-time prediction loss:

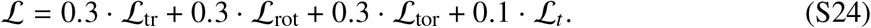

###### 7.2.5 Model architecture

The noisy ligand **L**_*ti*_ and receptor are assembled into a complex graph. Parallel GNN branches of PPFConv and GENConv (Section 6.3), consisting of 1 and 3 layers respectively, process this graph; their outputs are concatenated and passed through a transformer with geometry aware attention bias (Section 6.4). Separate score heads predict translation 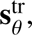 rotation 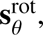 and torsion 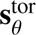 scores, which are combined into a pose update applied via Algorithm 1 to produce **L**_*ti*+1_.

###### Input features and embedding

Node features combine AEV (as described in Section 6.2), centered atomic coordinates, and sinusoidal Fourier encodings of the per DOF diffusion times (*t*_tr_, *t*_rot_, *t*_tor_). The 2140-dimensional AEV features are projected to *d*_model_ = 256 via a linear layer. Edges are featurized using point pair features (PPF) (*41*) which encode pairwise distances and relative orientations.

Because centered atomic coordinates are used directly as node features, the network is not SE(3)-equivariant by construction; the pocket-centered frame and the rigid-invariant edge features instead provide an approximate equivariance that is learned from data.

###### Score heads and readout

After the GNN and transformer, every atom carries a 512-dimensional contextual embedding. Three quantities are extracted per complex and shared across the global heads: the first transformer token ℎ_cls_ ∈ ℝ^512^ (a per-complex summary), the embedding of the ligand’s fixed origin atom ℎ_orig_ ∈ ℝ^512^ (the molecular anchor), and a task-specific diffusion-time embedding ℎ_*t*_^∗^ ∈ ℝ^512^ obtained by projecting the sinusoidal time encoding through a separate linear layer for each DOF ∗ ∈ {tr, rot, tor}.

Each score head produces a raw output that is rescaled by the corresponding noise-dependent factor, following the convention of (*44*).

###### Translation head

The translation score is predicted from a concatenation of the global summary, origin-atom embedding, and translational time embedding:

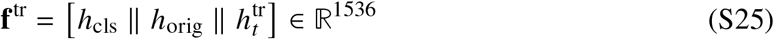

An MLP produces a raw three-dimensional vector. The final translational score is obtained by applying the noise-dependent output scaling described above:

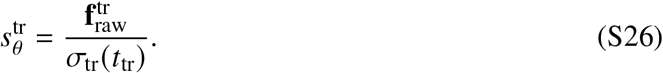

###### Rotation head

The rotation score uses an identical input structure with a rotation-specific time projection:

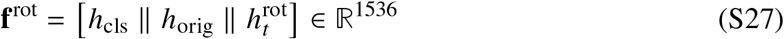

The same MLP architecture produces a raw three-dimensional tangent-space vector. The final rotational score is obtained by applying the noise-dependent rotational score-norm scaling:

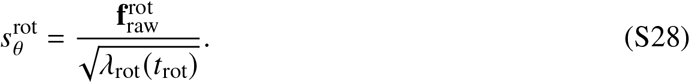

###### Torsion head

Unlike the global heads, the torsion head operates per rotatable bond *i* and uses a richer 5-part input of dimension 5 × 512 = 2560:

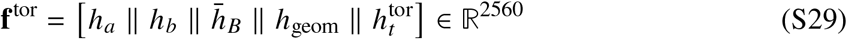

where *h*_*a*_, *h*_*b*_ ∈ ℝ^512^ are the transformer embeddings of the two atoms forming the rotatable bond, *h̄* _*B*_ ∈ ℝ^512^ is the mean embedding of all atoms on the rotating side, and *h*_geom_ = Linear(6, 512) applied to [**v**_⊥_ ∥ **b**] encodes the current torsion geometry: **b** is the unit bond vector and **v**_⊥_ is the unit vector from the bond midpoint to the rotating side centroid, projected perpendicular to the bond axis. This geometric pointer rotates physically with the torsion angle, allowing the model to distinguish its current position in angle space — information that is largely absent from the transformer features alone. An MLP produces a raw per-bond scalar.

A protein-reference residual is then added to break the remaining sign ambiguity. **p**_⊥_ is the unit vector from the bond midpoint to the pocket center of mass, projected perpendicular to the bond axis. The signed angle between the rotating fragment and the protein reference is encoded as:

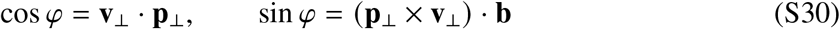

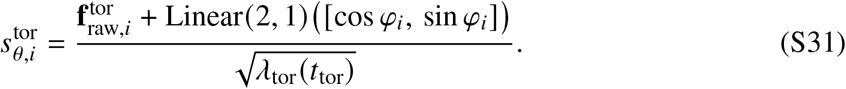

The residual linear layer is zero-initialized, so training begins from the pure 2560-dimensional head and gradually learns to exploit the directional correction.

###### Time-prediction heads

Three separate heads (tr, rot, tor) predict the diffusion time of each DOF purely from pose features, without access to any time embedding:

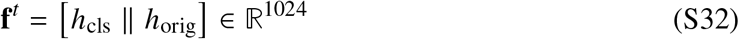

Each uses an MLP followed by a sigmoid to produce a scalar *t̂* ∈ [0, 1]. These heads act as a regularizer and monitoring signal; their loss is weighted at 0.1 in the total objective.

###### 7.2.6 Inference: reverse diffusion sampler

At inference time, RDKit with ETKDGv3 (*29*, *52*) is used to generate an ensemble of *N*_confs_ distinct ligand conformers. Each conformer is translated to the center of the target pocket and used to initialize *N*_traj_ independently sampled reverse-diffusion trajectories. The reverse process is used as an iterative refiner; trajectories from the same conformer differ only through the noise injected by the reverse-SDE update. The resulting diffusion batch therefore contains *N*_total_ = *N*_confs_ · *N*_traj_ trajectories.

The reverse-diffusion loop runs for 20 steps, and diffusion time decreases linearly from an initial start time 1 to 0. At each step, the complex graph is reconstructed from the current ligand coordinates, and the score network simultaneously predicts the translational, rotational, and torsional score components. The tempered reverse-SDE update, adapted from the temperature-guided sampler of Ingraham et al. (*53*), converts these predictions into incremental translation, rotation, and torsion transformations. We apply the same temperature parameterization independently to each degree of freedom, with the rotational and torsional updates taken in their respective tangent spaces; the corresponding parameters *ψ*^∗^, *T*^∗^_sampling_, and *σ*^∗^_data_. These increments are applied to the current ligand conformation using the transformation procedure defined in Section 7.1 and Algorithm 1. The complete reverse-diffusion sampling procedure is given in Algorithm 2.

###### Algorithm 2

Diffusionsampler – Reverse-diffusion sampling with temperature guidance

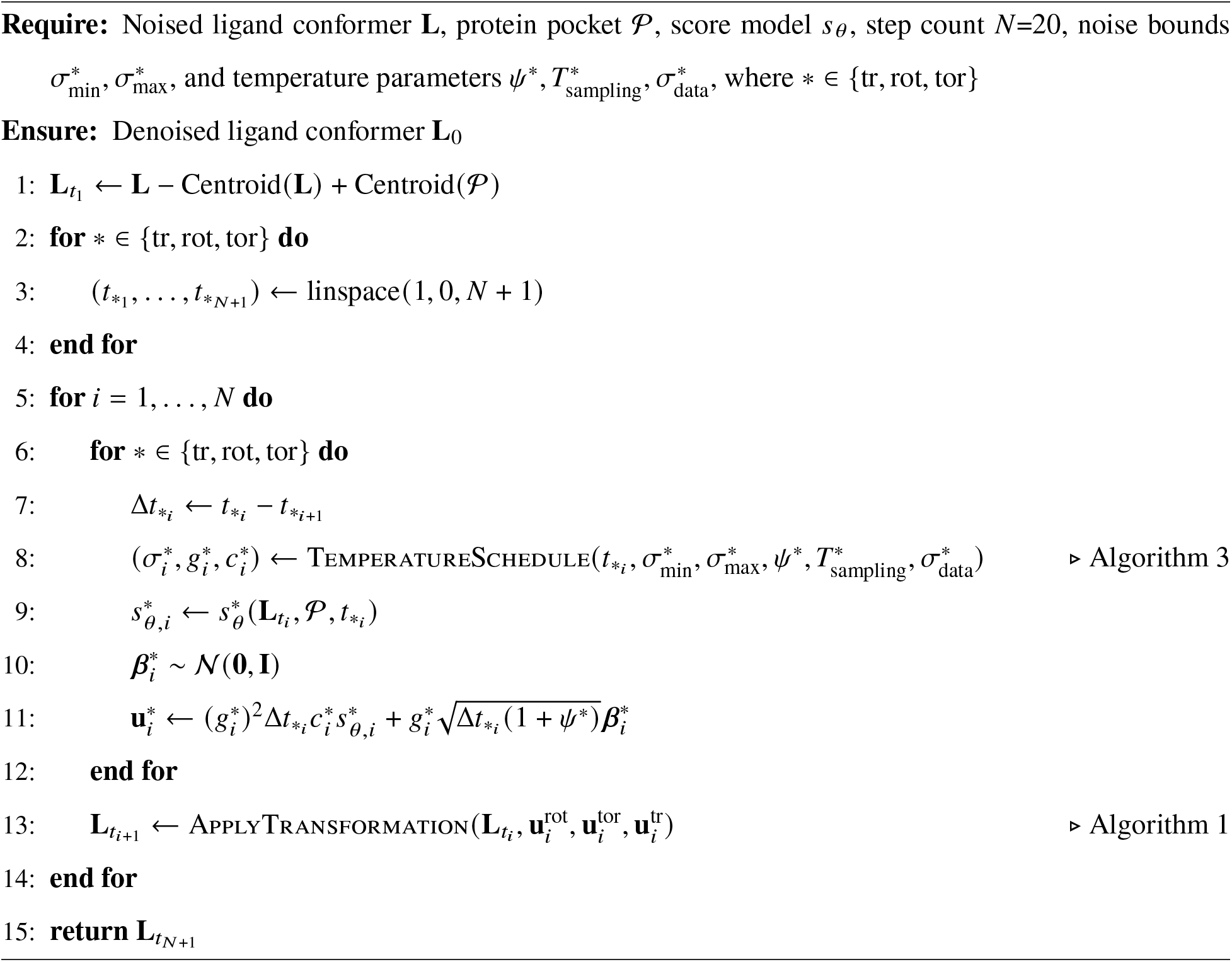

###### Algorithm 3

Temperature-guided noise schedule, adapted from (*53*).

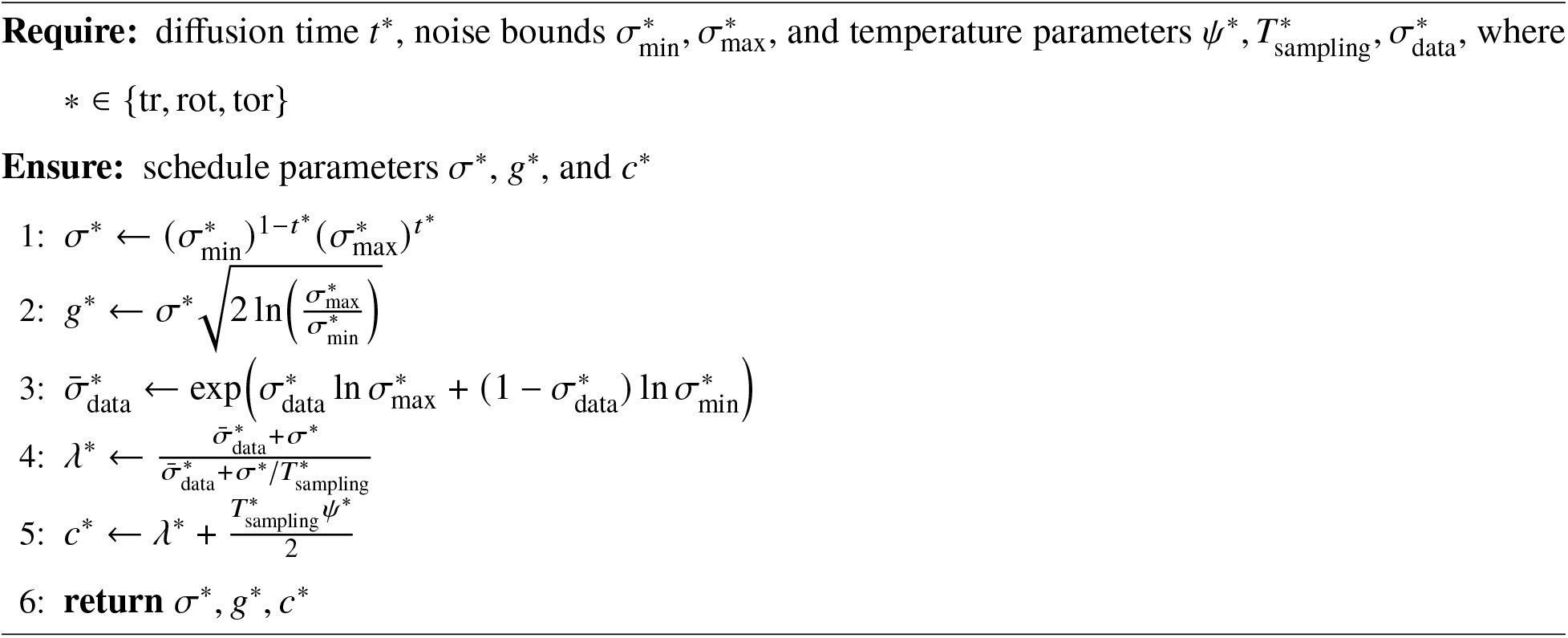

##### 7.3 DOFast score

###### 7.3.1 Score function model

The DOFast scoring function used by the search algorithms generalizes the Vina scoring function (*1*). The Vina scoring function sums the intramolecular term *S*^Vina^ and the intermolecular term *S*^Vina^, which describe the internal ligand interactions and the ligand–protein interactions, respectively. Both terms are expressed as sums of the same five atom-pair potentials with globally shared coefficients. For *t* ∈ {intra, inter}, the corresponding score is defined as

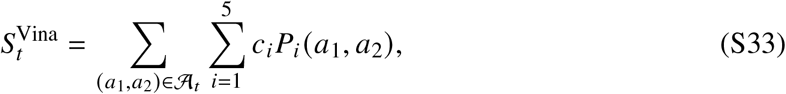

where *t* denotes the score-component type, A_*t*_ is the set of atom pairs contributing to that component, *P*_*i*_denotes the *i*-th atom-pair potential (Eqs. S37, S38, S39, S40, and S41), and *C*_*i*_ is its scalar coefficient:

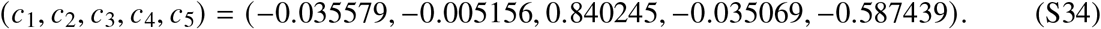

DOFast generalizes this formulation by allowing the coefficient of each potential to depend on the atom-type combination. The DOFast score is defined as

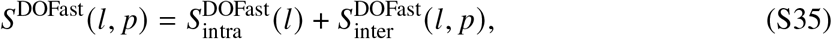

where *l* denotes the ligand and *p* denotes the protein. For *t* ∈ {intra, inter}, each score component is defined as

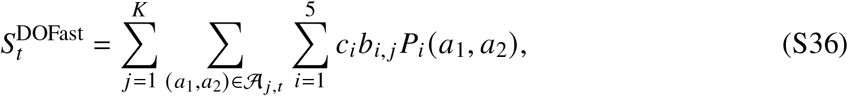

where *K* is the number of atom-type combinations, A_*j*,*t*_ is the set of atom pairs belonging to combination *j* and contributing to score component *t*, and *b*_*i*, *j*_ is the atom-type-dependent scaling coefficient for potential *i*, as described in more detail below. The dependence of *S*^DOFast^ on *l* and *p* is omitted in Eq. S36 for notational simplicity.

For the five selected atom types listed in Table S15 (*54*), each unordered atom-type pair, including same-type pairs, is assigned a distinct set of five scaling coefficients *b*_*i*, *j*_. All remaining atom-type pairs are assigned to a single additional group and share the same set of five coefficients. Thus, the atom pairs contributing to score *s* are partitioned into *K* = 16 groups, and, for atom pairs in group A_*j*,*t*_, the coefficient of the *i*th potential is *C*_*i*_× *b*_*i*, *j*_. This parameterization results in 80 scaling coefficients in total.

**Table S15:** Selected XS atom types defining the pair-specific coefficient groups in DOFast.

| XS Type | Description | Hydrophobic | H-Donor | H-Acceptor |
| --- | --- | --- | --- | --- |
| XS_TYPE_C_H | Carbon; not bonded to any heteroatom | ✓ | × | × |
| XS_TYPE_C_P | Carbon; bonded to at least one heteroatom | × | × | × |
| XS_TYPE_N_D | Nitrogen hydrogen-bond donor only | × | ✓ | × |
| XS_TYPE_O_A | Oxygen hydrogen-bond acceptor only | × | × | ✓ |
| XS_TYPE_O_DA | Oxygen acting as both donor and acceptor | × | ✓ | ✓ |

As the score is designed for ranking purposes, the relative value of the score between different structures of the same molecule is calculated, so the atom pairs included in the rigid parts of the molecule structure, having the same distance, independent of the structure transformation, are excluded from the score computation. The atom pairs, having a graph distance less than or equal to 3 in a molecule graph with chemical bond edges, are also excluded from the score computation. While the exclusion of bonded atom pairs prevents the physical-potential values from exploding because of the small distance between them, the exclusion of atom pairs with 2 and 3 graph distance is a computation time optimization.

Let *r* (*a*) denote the van der Waals radius of atom *a*, and let *x* = ∥*a*_1_ − *a*_2_∥ _2_ − *r* (*a*_1_) − *r* (*a*_2_) denote the surface distance between atoms *a*_1_ and *a*_2_. The five pairwise potential functions are defined as follows:

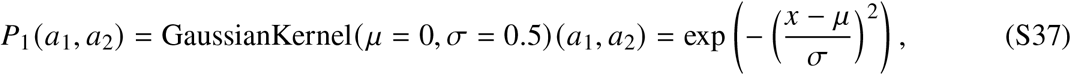

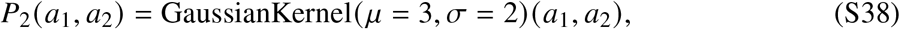

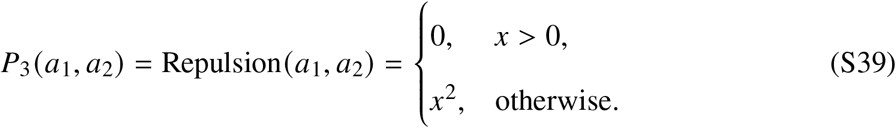

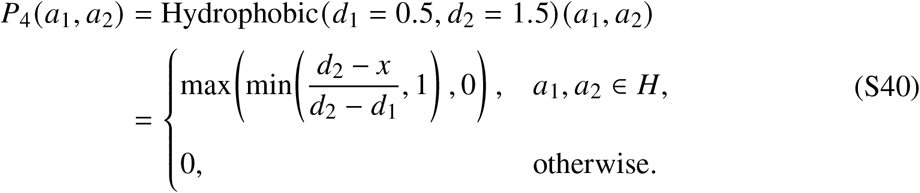

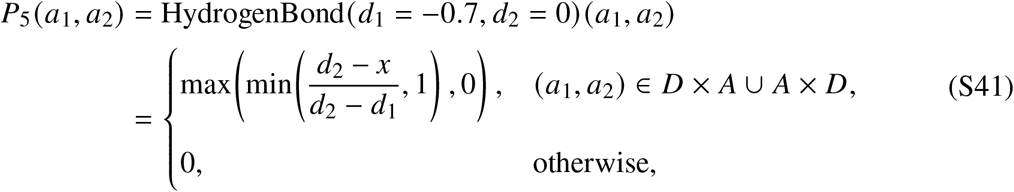

where *H* is the set of Hydrophobic atom types, *A* and *D* are the sets of acceptors and donors respectively.

###### Algorithm 4

DOFast score model training

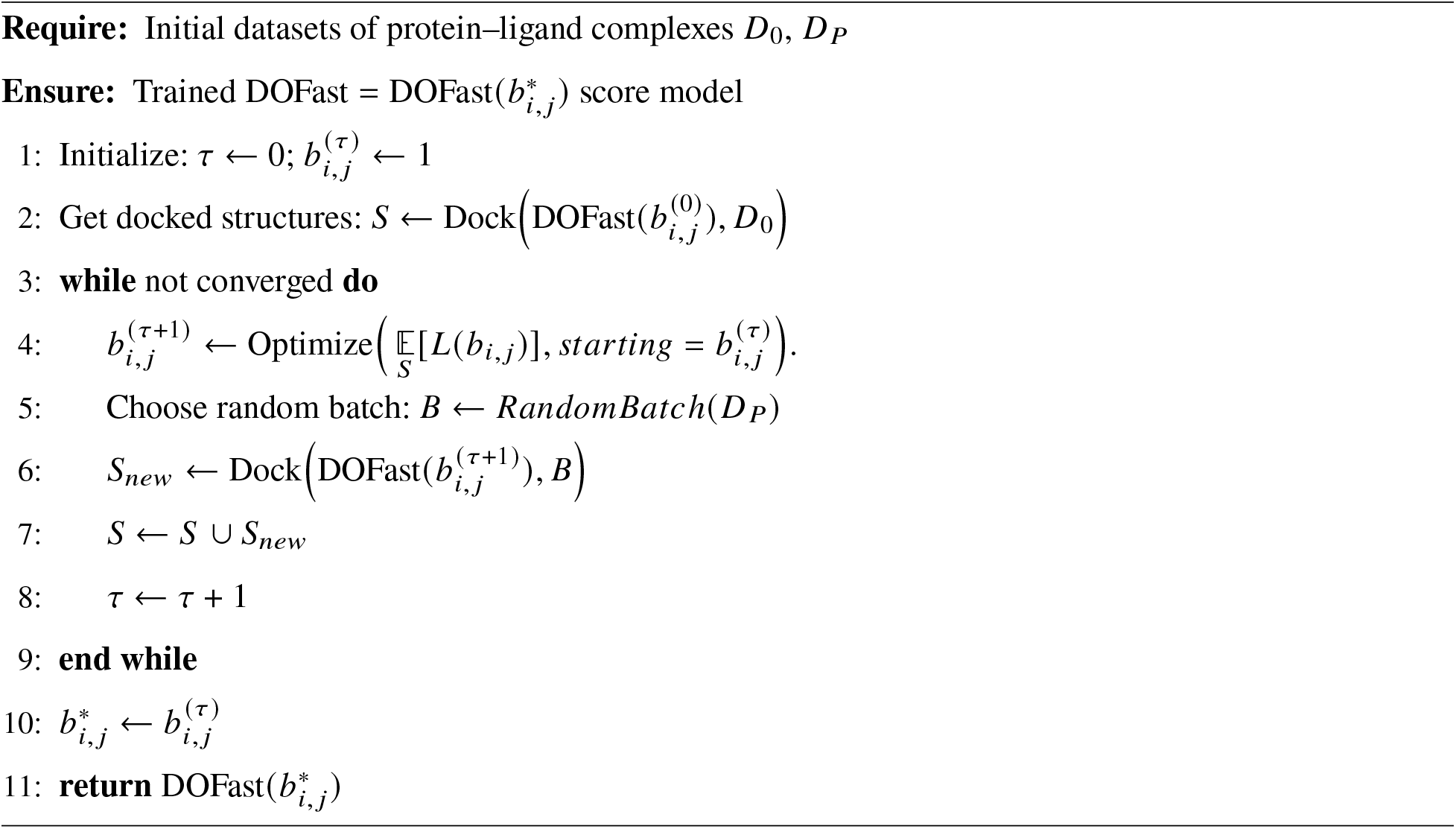

Let *D*_0_ denote the complete docking dataset, as described in Section 4.1. For model training, we select the subset *D*_*P*_ ⊂ *D*_0_ containing complexes whose corresponding experimental structures are included in PDBbind v2020 (*25*). We then optimize the DOFast scoring model using the iterative procedure described in Algorithm 4 and the objective function *L* = *L*(*b*_*i*, *j*_) defined in the following section.

When DOFast was retrained on benchmark-specific training splits, the resulting coefficient values remained approximately unchanged, showing no notable dependence on the benchmark-similar examples removed during filtering and consistent with the model’s limited parameterization (80 parameters).

###### 7.3.3 Optimization objectives

The idea of modeling a funnel-shaped energy score function was utilized in protein structure prediction tasks (*55*), (*56*), (*57*). Analogous methods were used to optimize the protein-ligand binding energy. The loss function for the optimization is the following:

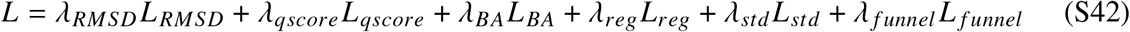

where

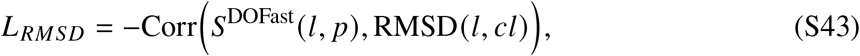

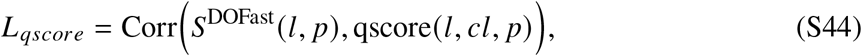

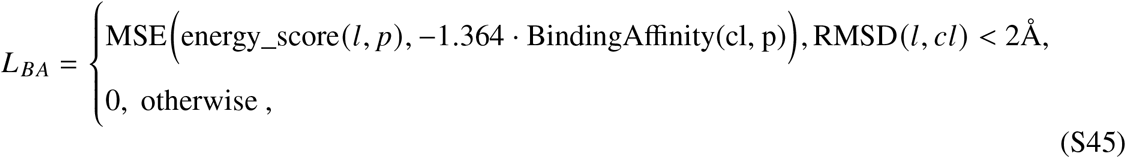

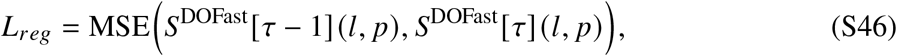

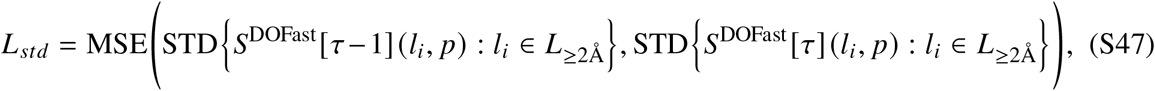

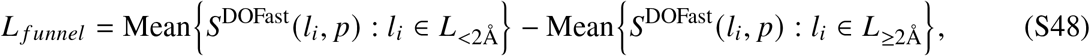

where RMSD is the symmetry-corrected root mean squared deviation, *l* and *Cl* are the ligand with query and crystal structures respectively, *p* is the protein, energy_score S49 is the binding affinity approximation, qscore is the contact map score defined in S50, *S*^DOFast^ [*τ*] is the score with 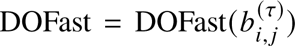 model at optimization step *τ*, *L*_<2Å_ and *L*_≥2Å_ are the ligand structures having less than 2 Å and greater than or equal to 2 Å RMSD from the crystal structure respectively,

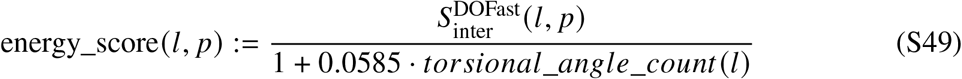

##### 7.4 Search algorithms

In this section, we introduce the search algorithms used with the DOFast scoring function to identify high-scoring docking solutions in the transformation space defined in Section 7.1.

###### 7.4.1 Monte Carlo search

We employ a Monte Carlo (*58*) search augmented with local refinement to optimize ligand poses. The complete search pipeline is presented in Algorithm 5, while the optimization of an individual solution is detailed in Algorithm 6. Local refinement is performed using either BFGS or the Nelder–Mead simplex method.

###### Algorithm 5

MonteCarloSearch

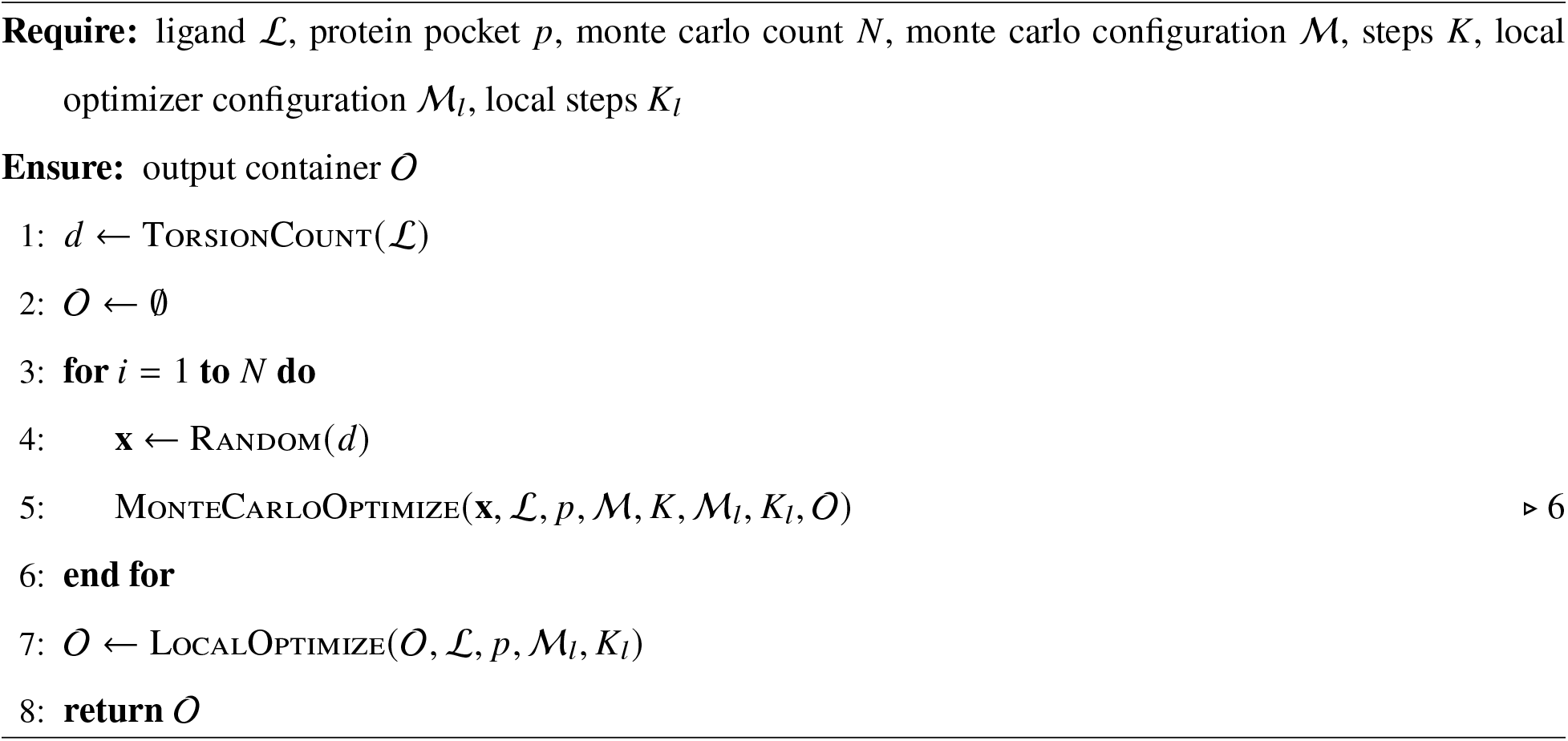

###### Algorithm 6

MonteCarloOptimize

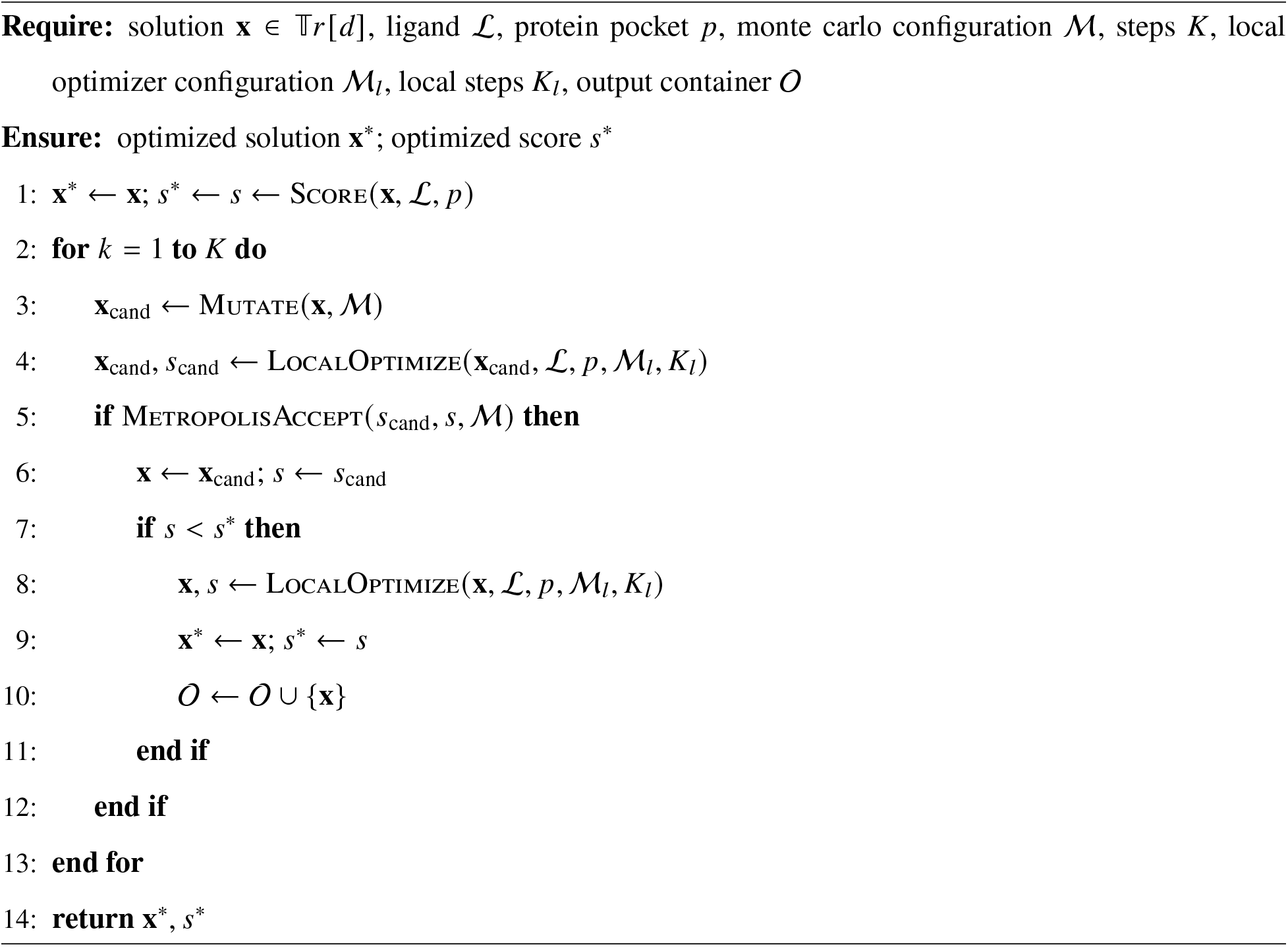

###### 7.4.2 Enhanced Scatter Search

The classical eSS search (*59*) is another well-known optimization method. Here we introduce a novel modification that includes two conceptual changes:

1. The combination candidates replace the solutions iteratively and not at the same time. This change provides better candidates for subsequent solutions as the previous solutions are updated.
2. The stagnation process is removed due to the fact that we apply the search on top of the diffusion sampling and consider the input solutions to be "near-optimal", hence the modified name ESSNoExplore.

The complete ESSNoExplore pipeline is presented in Algorithm 7, with each individual optimization step performed according to Algorithm 8.

###### Algorithm 7

ESSNoExplore

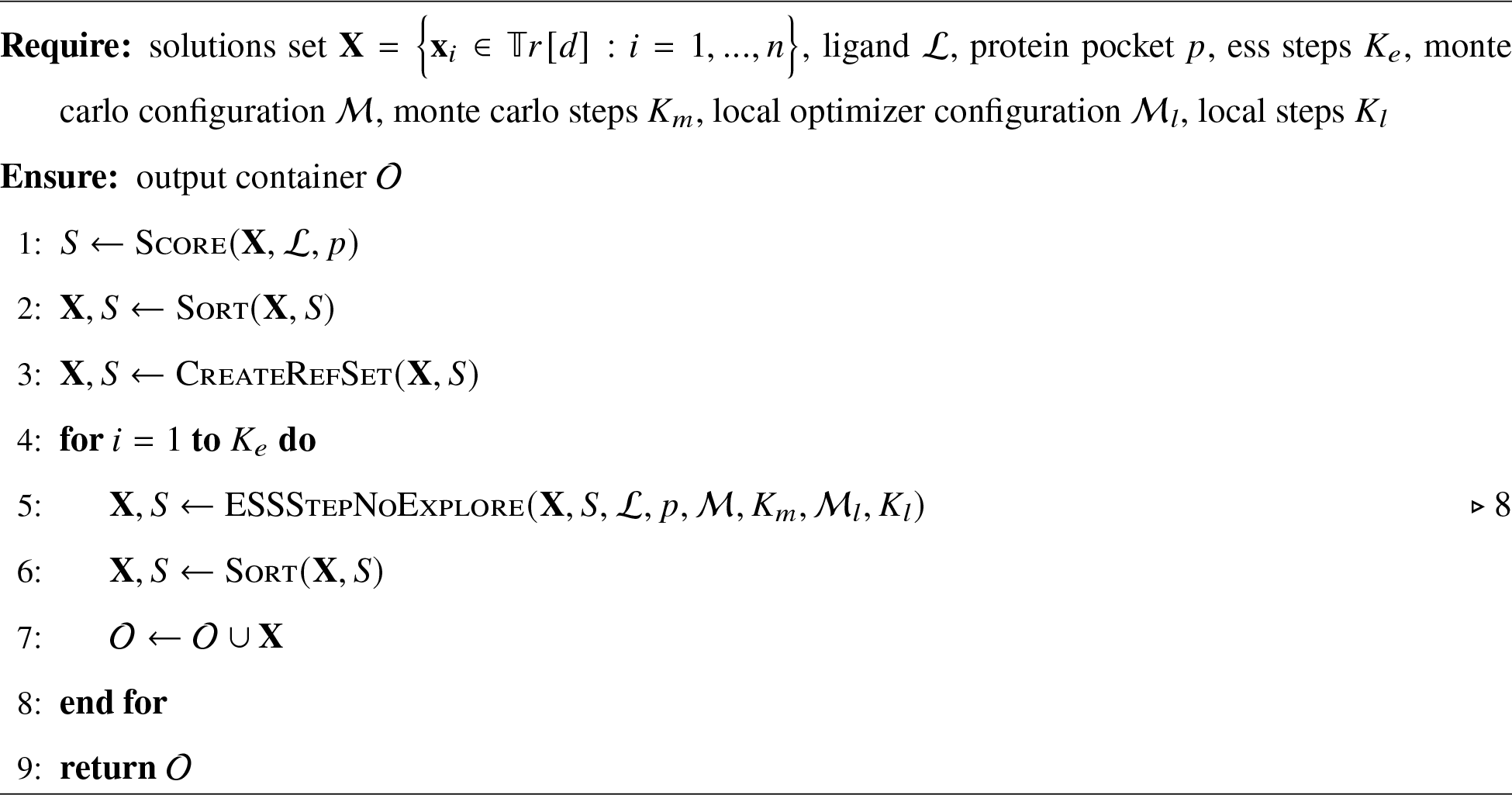

###### Algorithm 8

ESSStepNoExplore

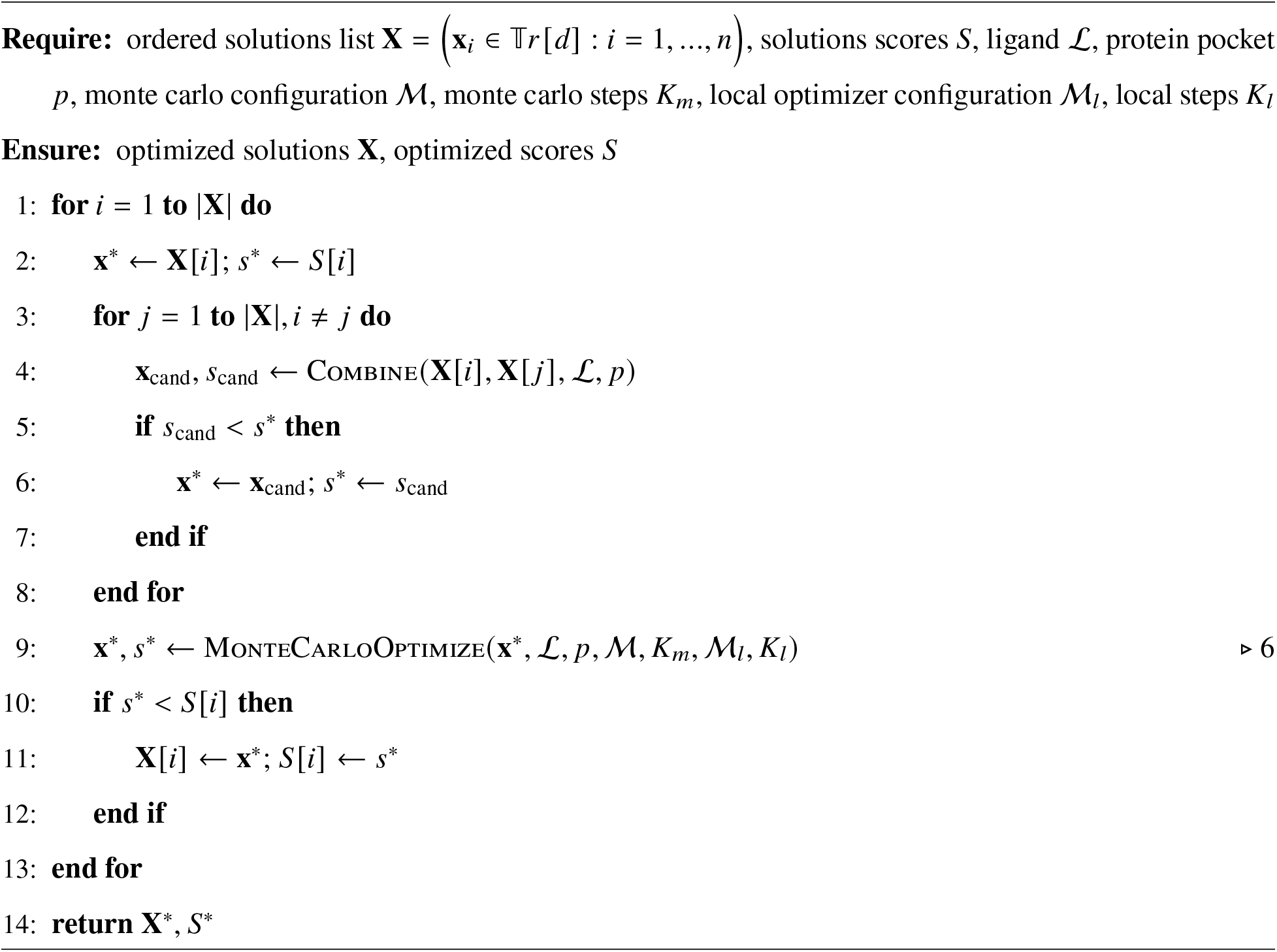

##### 7.5 Ranking score

In this section, we introduce the ranking models used to select the top-ranked ligand pose from the candidates generated during the search phase. For training, we define target scores with values in the interval [0, 1] that measure the agreement between a candidate pose and the crystallographic ligand pose, including their respective protein–ligand contacts. We then train neural networks to predict these scores from the candidate ligand pose and protein structure, without access to the crystallographic pose at inference time. The final selection procedure, which uses two ranking models, is described in Algorithm 9.

###### Algorithm 9

SelectTopNByRanking

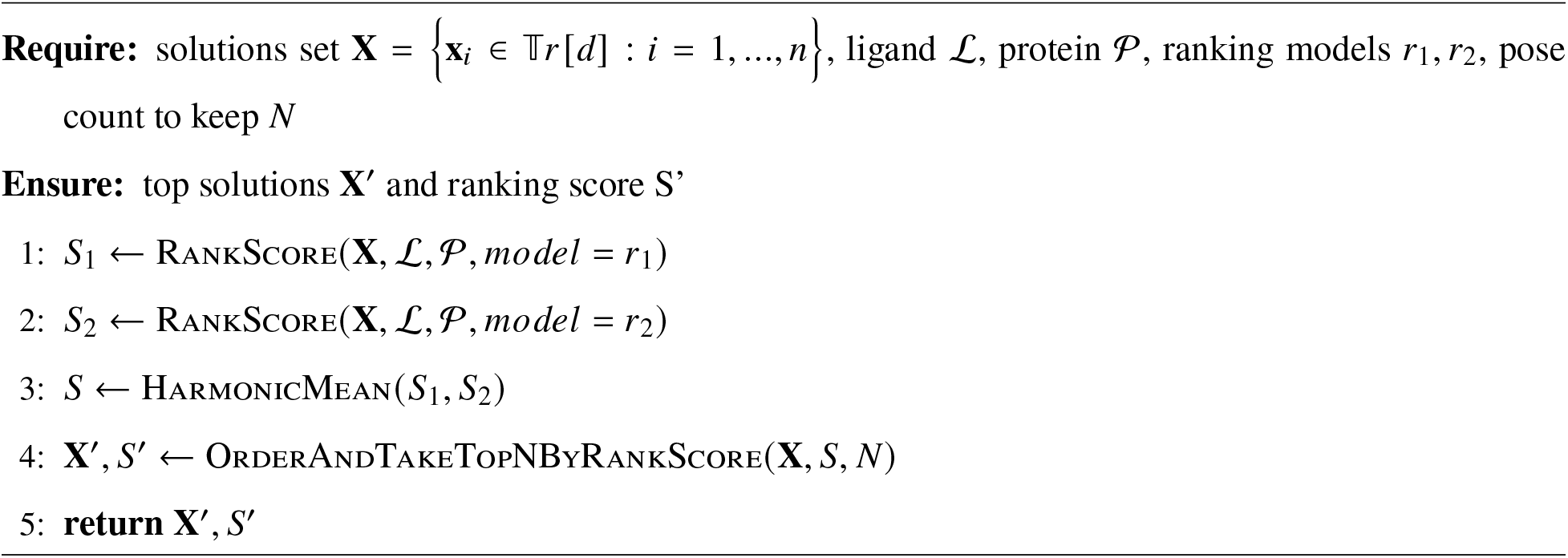

###### 7.5.1 Target scores

The target scores are defined as contact-map-based scores. We define both ligand-level and atomlevel scores.

###### Qscore

Following the contact-preservation logic of the native-contact *Q* measure (*60*), we define a protein–ligand variant for a crystallographic ligand (target ligand), a ligand in a different conformation (query ligand), and the protein. The score measures how many protein contacts of the target ligand are preserved by the query ligand, with a value of 1 indicating the best match to the target. It is defined as:

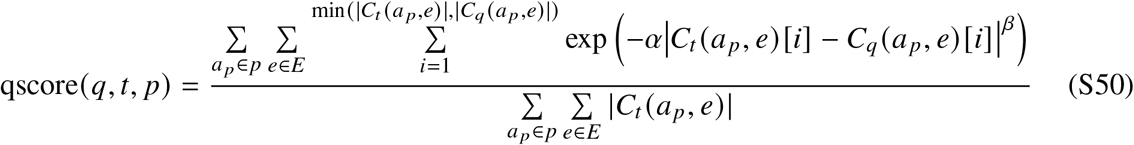

where *q*, *t*, *p* are the query ligand, the target (crystal) ligand and the protein respectively, *a*_*p*_-s are the protein atoms, *E* is the set of the element types of the molecule atoms, *C*_*q*_ (*a*_*p*_, *e*) and *C*_*t*_ (*a*_*p*_, *e*) are the contact distance sorted arrays of the protein atom *a*_*p*_ from the atoms of type *e* of the query and target ligands respectively. The contacts are defined to have distance up to 4.5 Å for crystal atoms and 5.5 Å for query ligand atoms. The *α* = 0.3 and the *β* = 4.

###### Atomic LDDT-PLI (local Distance Difference Test - Protein-Ligand Interaction)

The LDDT-PLI score (*61*) is the modified version of classical LDDT score (*62*) to be a ligand-level interaction score. Here we define its atom-level analog. To do so, we first find the best match between crystal and query ligand atoms in the molecule graphs, using the DockRMSD tool (*63*). The idea of the score is similar to the one described in the previous section, that is to describe which part of the protein atom contacts of the crystal atom are preserved in case of the query ligand atom. The contact value function is chosen to be step-function instead of the gaussian variant used in QScore. Here is the formula:

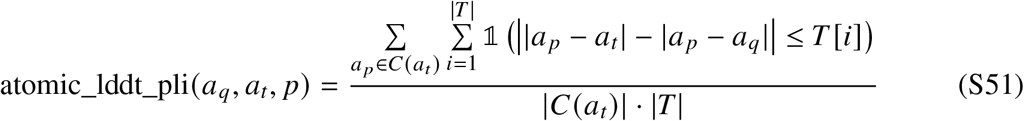

where *a*_*q*_, *a*_*t*_and *p* are the query ligand atom, target ligand atom and the protein, the *C*(*a*_*t*_) are the protein contact atoms of the crystal ligand atom *a*_*t*_. The *T* is the threshold set. The contacts are defined to have distance up to 15 Å and the *T* = {1, 2, 4, 8}.

These atomic scores are utilized both as raw target scores and as aggregats using the harmonic mean function to represent a ligand-level score:

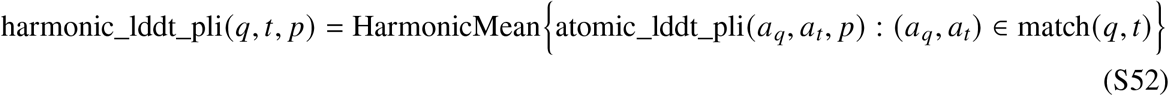

###### 7.5.2 Neural network architecture

The neural network architecture consists of three main parts: the roto-translation invariant AEV featurizer (Section 6.2), Graph Convolutional Network (GCN (*64*), (*65*)) block and the final attention module (*43*), (*66*). The model (Figure S11) is a multi-head neural network aiming to predict each of the target scores.

**Figure S11:**
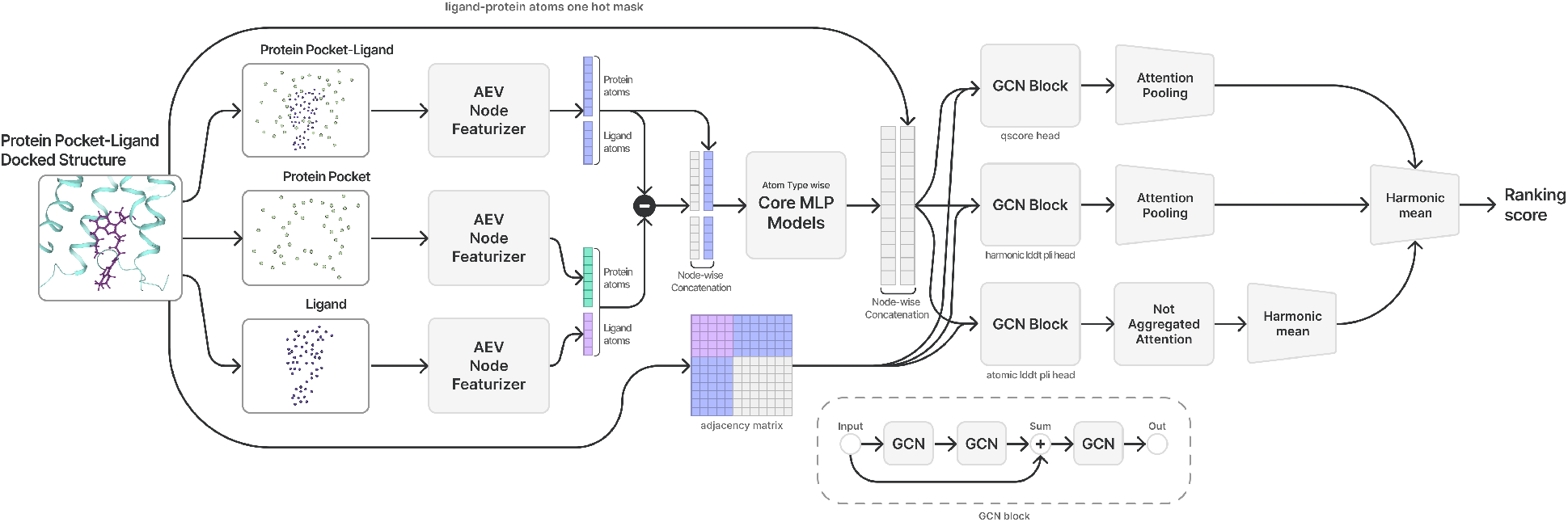
The multi-head ranking model. The figure shows the ranking score inference for an input protein-ligand docked structure.

###### Input processing

The input of the model consists of the following components:

- atom types and coordinates of the ligand
- atom types and coordinates of the protein pocket

The protein pocket is defined as the union of the 32 closest protein atoms within 10Å of each ligand atom.

###### GCN

For each target score, the graph for GCN block is constructed using AEV as node features and the edges are defined with distance threshold 10 Å between ligand and protein atoms and 4 Å between ligand atoms.

###### Attention module

For each target score, the attention module is an attention pooling layer in case of ligand-level (scalar) target scores and an unaggregated attention for atomic (vector) target scores. A linear projection layer follows in order to get a scalar value in case of ligand-level target heads and a vector for atom-level target head. The input for the module is the output features of the corresponding GCN block.

###### Head aggregations

The target scores include 2 ligand-level and 1 atom-level scores. The atom-level score vector is aggregated by the harmonic mean function in order to represent a scalar ligand-level score. The choice of harmonic mean aggregator is based on the bias towards the smallest value in the array, meaning that if one of the atoms is misplaced, then the overall ligand has a wrong position as well. The 3 heads are again aggregated by the harmonic mean aggregator resulting in a single scalar confidence score.

###### 7.5.3 Dataset

The source dataset comprises the protein–ligand complexes described in Section 4.1. For each benchmark-specific model, the complexes were filtered according to the corresponding similarity-based split, as described in Section 5. Each complex was docked using the procedures described in Sections 7.2 and 7.4, and the target ranking scores were computed for the resulting ligand poses. To increase pose diversity, we generated candidates using both the complete docking pipeline and its individual optimization techniques as standalone pose-generation procedures. For complexes for which these procedures did not generate near-native poses, we supplemented the dataset, when possible, using the optimized aligned ligand poses retained by the reference-guided preprocessing procedure described in Section 4.1. Additional near-native candidates were generated by applying small random perturbations to these aligned poses. The final ranking dataset therefore contains candidates obtained using multiple pose-generation and optimization procedures.

###### Adversarial examples finetuning

The following procedure was performed in order to mitigate the false negative type of errors, i.e. examples with a bad score but small RMSD.

1. Start from the aligned ligand structure described in Section 4.1,
2. Perform reverse optimization of the current score using Monte Carlo and local refinement algorithms described in Section 7.4,
3. Collect structures with bad score, but small RMSD.

###### 7.5.4 Training procedure

The training objective is a weighted MSE loss on each of the prediction heads’ attention module output and the true target:

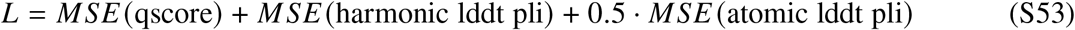

The model was trained with Adam optimizer (*49*) with a learning rate of 10^−4^, no learning-rate scheduler, and a batch size of 128. The key point of the training procedure is the two sampling techniques that provide different training dynamics.

1. The first one orders the data samples by the order of the docking sources described in the previous section. Each of the docking sources (except for the once obtained with data augmentation techniques or constrained docking) represents a complete data set on its own, which explains the faster convergence behavior of this type of training.
2. The second one orders the data samples by the order of the complexes. This approach ensures the model to see the structures of the same complex at once which results in better understanding of the physical energy landscape of the model. This method provides more stable, yet longer training dynamics.

We divide each epoch into 20 smaller phases. The weights of the first model are obtained by averaging the weights from phases 38–42, corresponding to the end of the second epoch and the two phases on either side of it. The weights of the second model are obtained by averaging the weights from phases 78–82. As a result, we get two different models that are combined in order to perform ranking.

##### 7.6 Docking pipeline

This section combines the components described above into the complete end-to-end docking procedure and defines the default inference protocol used in our experiments. Unless otherwise specified, all docking experiments use this protocol.

Candidate poses are first generated using the diffusion-based sampling procedure described in Section 7.2. Under the default inference settings, we generate *N*_confs_ = 4 starting ligand conformers and sample *N*_traj_ = 4 independent reverse-diffusion trajectories from each conformer, yielding 16 candidate poses. The resulting poses are refined using the procedure in Algorithm 10, which applies the global and local optimization methods described in Section 7.4 under the DOFast score function. The refined candidates are then evaluated using the two ranking models, and the final pose is selected according to Algorithm 9. The complete end-to-end procedure is summarized in Algorithm 11.

The contributions of the individual pipeline stages are evaluated in Section 1.2.1, while the runtime–accuracy trade-off associated with varying *N*_confs_ and *N*_traj_ is examined in Section 1.2.3.

###### Algorithm 10

PostDiffusionRefinement

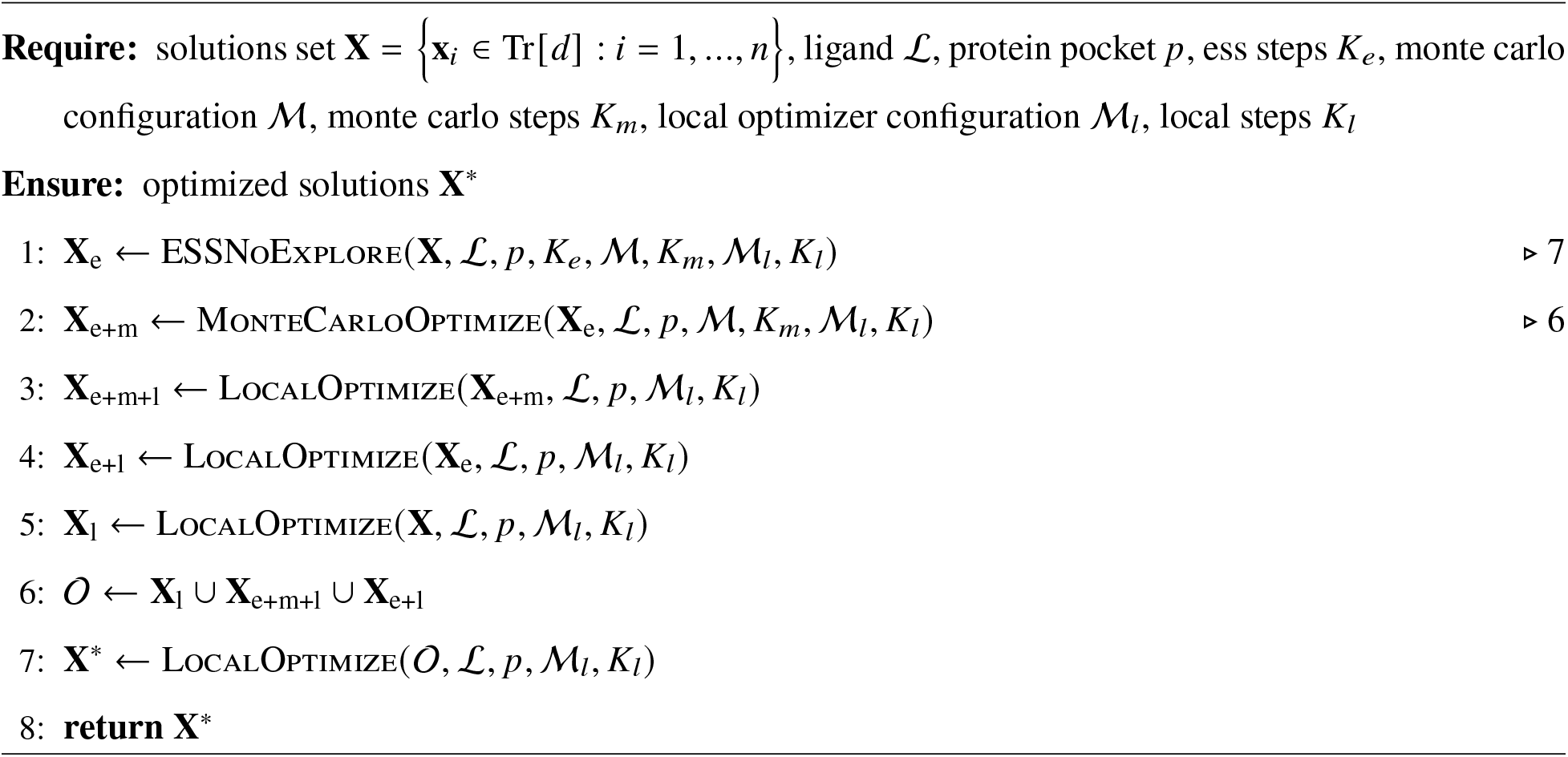

###### Algorithm 11

DockingPipeline

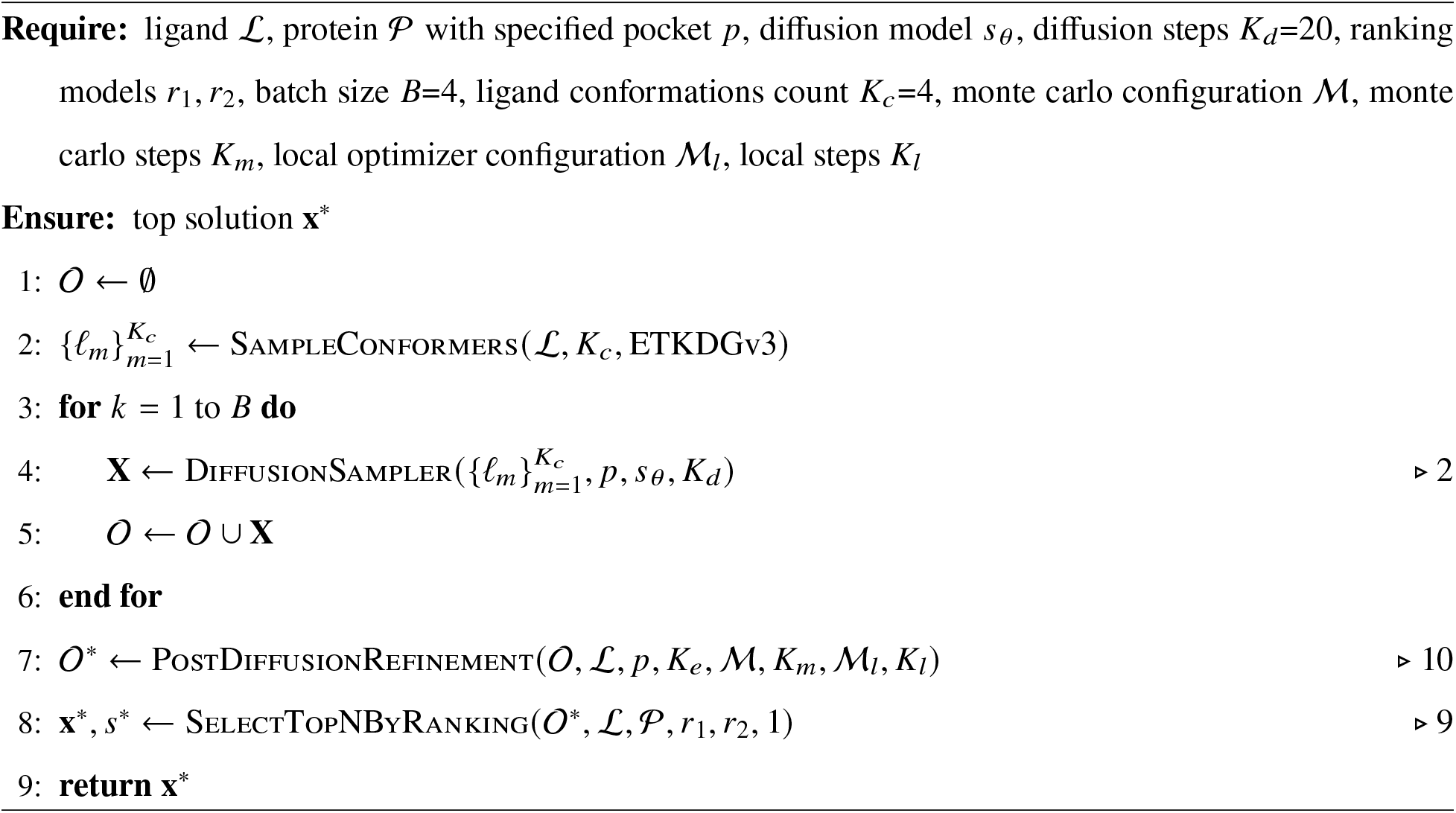

##### 7.7 Blind docking

For blind docking, we first identify candidate cavities directly from the input protein structure using a grid-based geometric procedure. A Cartesian grid with 1 Å spacing is constructed to cover the full protein, and grid points overlapping protein atoms are removed. For each remaining grid point, we estimate local enclosure by sampling 50 random line orientations and casting rays in both directions along each line. This yields two scores: *N*_1_, the number of sampled lines for which the protein is intersected in any direction, and *N*_2_, the number of sampled lines for which the protein is intersected in both directions. Intuitively, *N*_1_ captures points near protein boundaries and grooves, while *N*_2_ captures points embedded within more enclosed cavities.

We retain grid points whose *N*_1_ and *N*_2_ scores are both at or above the corresponding 75th-percentile thresholds. To remove sparse or noisy regions, points with fewer than 20 retained neighbors within 2 Å are iteratively discarded until convergence. The remaining points are connected using a 2 Å adjacency cutoff and grouped into connected components. The *K*_*p*_ largest components are retained as candidate cavities and used as the initial search regions for the blind-docking pipeline. The complete cavity-detection procedure is given in Algorithm 12.

###### Algorithm 12

PocketFinder

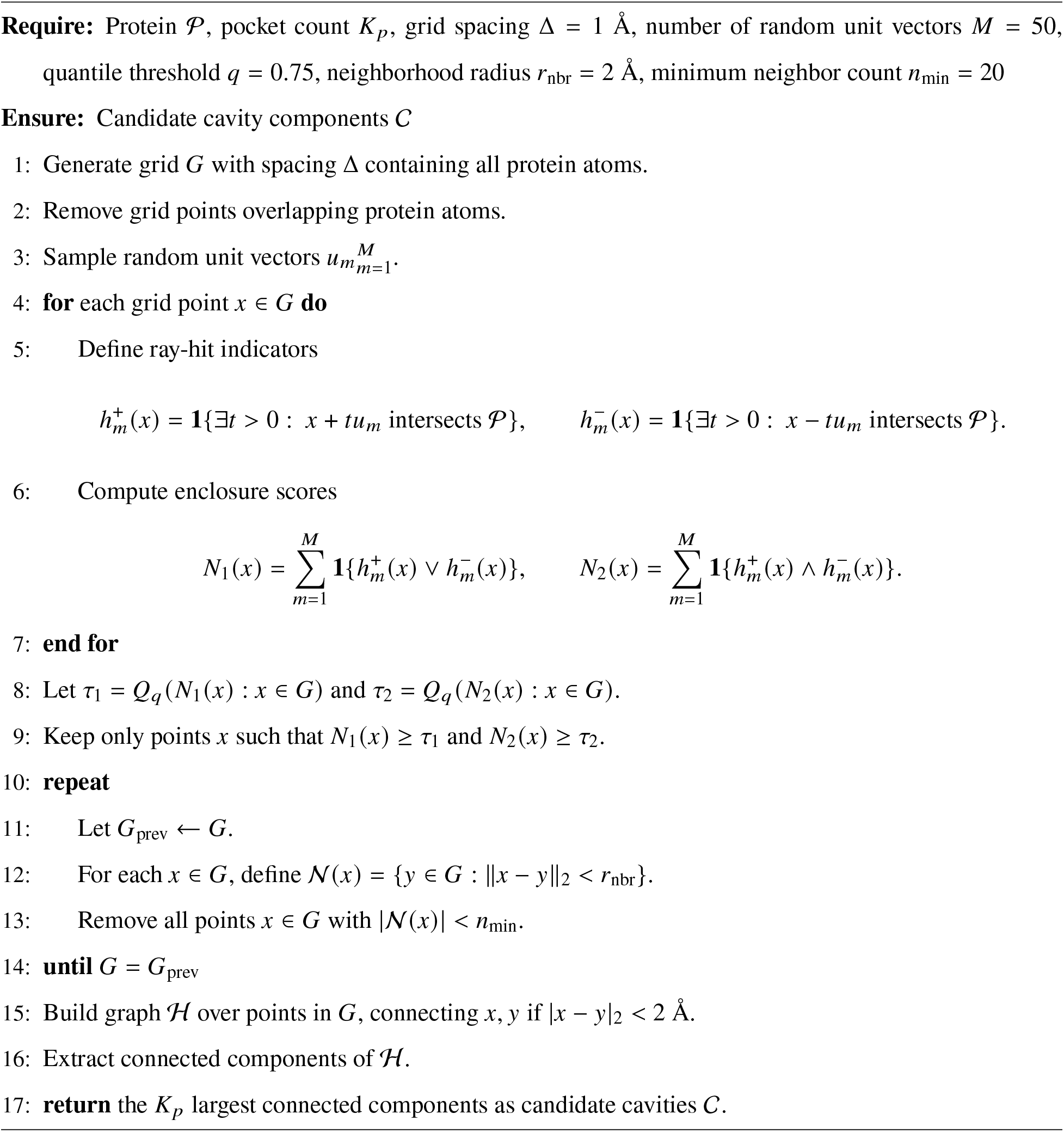

The detected cavity components serve as the initial search regions for the blind-docking pipeline. Within each cavity, a preliminary Monte Carlo search is performed to generate ligand poses, which are then used to define the corresponding set of protein pocket atoms supplied to the diffusion sampler. After diffusion sampling, the generated poses are clustered, and the resulting clusters are used to redefine the candidate pockets. Post-diffusion refinement is then applied independently within each redefined pocket. Finally, the refined candidates from all pockets are pooled and evaluated using the ranking models, and the overall top-ranked pose is returned. The complete procedure is described in Algorithm 13. The performance of the blind-docking pipeline is evaluated in Section 1.2.2, where it is compared with the pocket-defined docking protocol.

###### Algorithm 13

BlindDockingPipeline

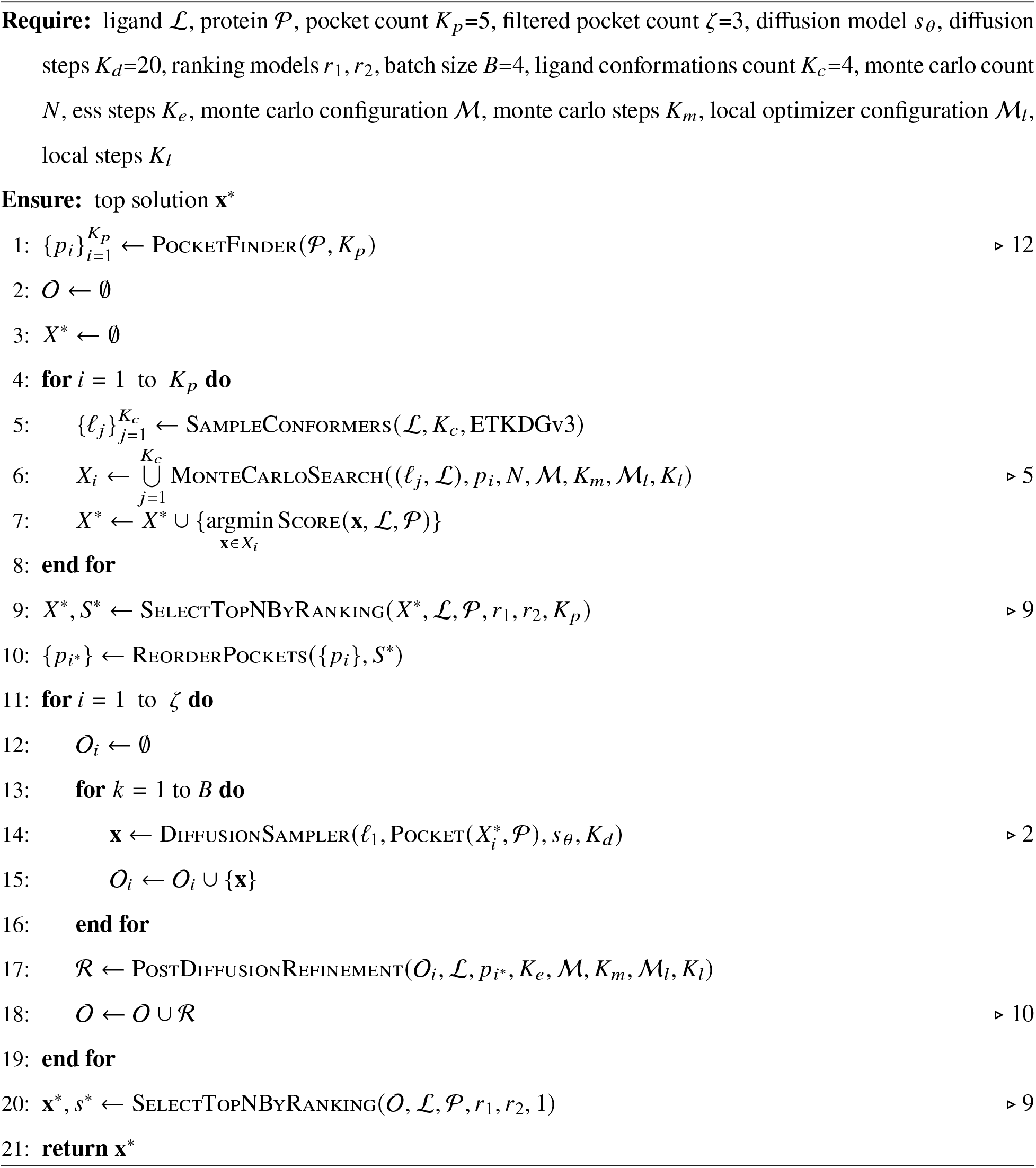

#### 8 Scoring model

##### 8.1 Scoring data generation

For training the scoring model, DOScore, we generated structure-based data from the PocketAffDB-derived assay records and the corresponding reprocessed RCSB proteins (Section 4.2). For each assay, we used the co-crystallized ligands associated with the PocketAffDB pocket entries to define the relevant binding sites, and docked all bioactivity-labeled assay ligands into the associated binding sites. Docking was performed separately for each ligand–pocket pair, and only the top-ranked pose was retained. This produced a set of docked complex structures in which assay ligands were placed into assay-specific binding sites. As a result, we obtained a dataset of docked protein-ligand complex structures with experimentally measured bioactivity labels.

For hit-identification training, we additionally generated decoy molecules for each bioactivity-labeled assay ligand using the procedure described in the following section. The decoys were docked into the same assay-associated pockets using the same protocol, again retaining only the top-ranked pose for each decoy–pocket pair. Thus, we obtained a dataset for hit identification training, which for each assay contained docked structures for its bioactivity-labeled ligands and corresponding decoy molecules, all placed into the binding pockets linked to that assay.

The application of the generated datasets is described in Section 8.3.

###### Decoy generation method

For the enrichment of our training dataset we developed a computational pipeline for generating high-quality molecular decoys (Alg. 14). Given a set of bioactivity-labeled assay compounds (ligands) for a protein target, the pipeline selects decoy molecules that are physically similar to the assay ligands in terms of drug-like physicochemical properties but are structurally distinct, thereby reducing the likelihood that decoys themselves are binders. The pipeline draws candidates from large public chemical databases (ChEMBL and BindingDB) and applies a multi-stage filtering procedure that enforces both property similarity constraints and structural dissimilarity thresholds.

The construction of property-matched, topologically dissimilar decoy sets follows the rationale established by DUD-E (*7*) and DEKOIS 2.0 (*39*) with specific modifications described below.

For each assay, decoys were generated as follows:

###### Algorithm 14

GenerateDecoys

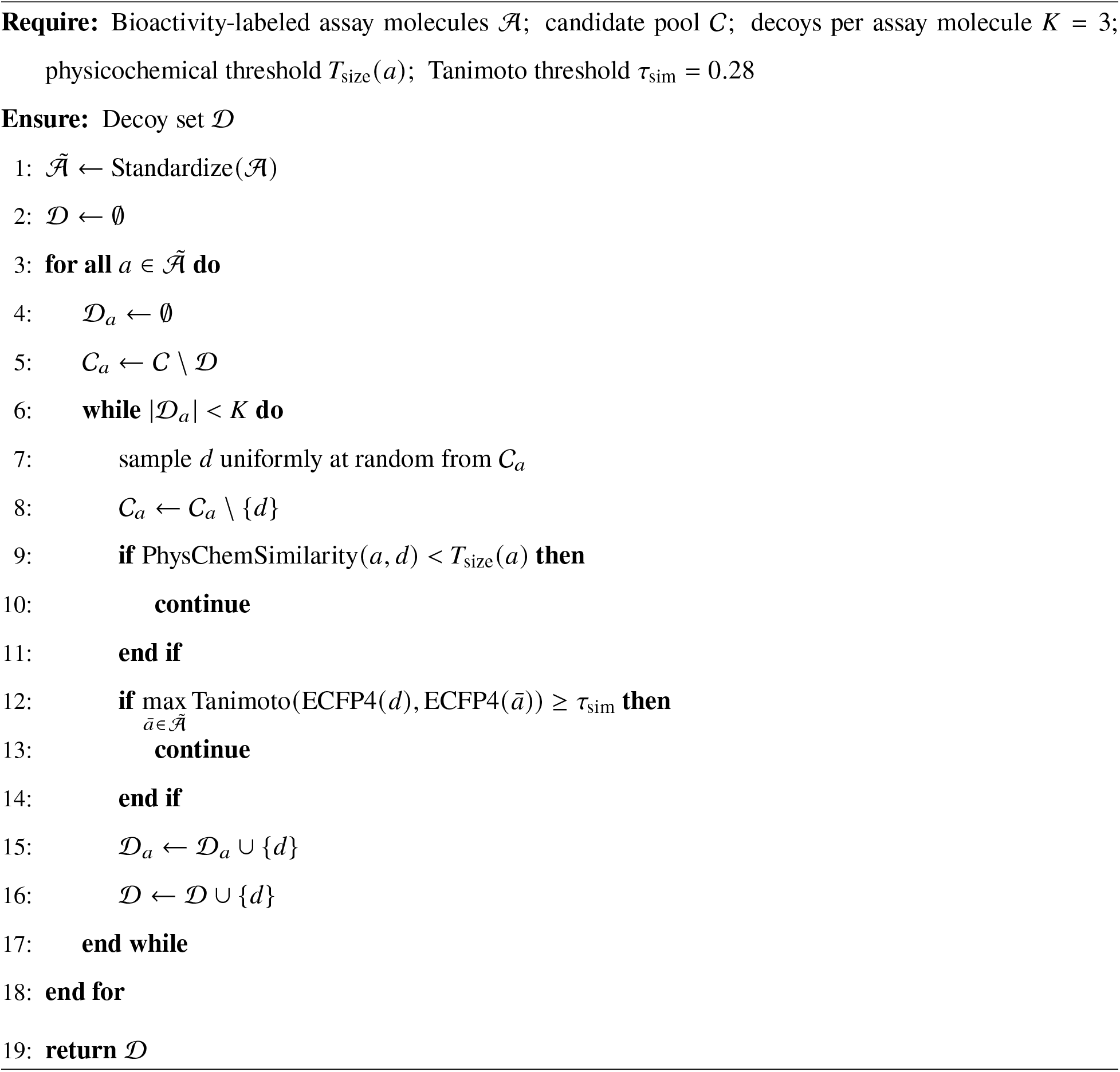

###### 1. Assay molecule standardization

Bioactivity-labeled molecules from experimental assays were standardized by removing salts, solvents, and known counter-ions, and retaining the largest fragment by heavy atom count.

###### 2. Physicochemical property matching

For each assay molecule, candidate decoys were evaluated using a physicochemical similarity score defined as a weighted sum of binary property-matching indicators. The descriptors included molecular weight, hydrogen-bond donor and acceptor counts, topological polar surface area, QED, number of aromatic rings, number of rotatable bonds, and occurrence counts for selected atom types. For each descriptor, a match was assigned when both the absolute and relative differences between the assay molecule and candidate decoy were within descriptor-specific thresholds. The individual matches were combined into a weighted similarity score, and candidates were retained if the score exceeded a size-dependent threshold. This threshold was relaxed for larger bioactivity-labeled molecules to allow greater descriptor variation with increasing molecular size.

###### 3. Topological dissimilarity filtering

Property-matched candidates were required to be structurally dissimilar to all molecules in the same assay. Structural similarity was computed using Morgan fingerprints equivalent to ECFP4 (radius=2, 2048 bits), and candidates were accepted only if their Tanimoto similarity to every assay molecule was below 0.28.

###### 4. Decoy selection

For each assay molecule, candidate molecules were sampled randomly from the candidate pool and evaluated against the physicochemical and topological filters. A sampled candidate was added to the decoy set if it passed both filters. Sampling continued until three decoys had been selected for the assay molecule. Selected decoys were not reused across molecules within the same assay.

The resulting decoy set therefore occupied the local physicochemical neighborhood of the assay molecules while remaining topologically distinct from them.

##### 8.2 Model architecture

DOScore’s architecture follows a hierarchical design that combines graph-based representation learning with transformer-based (*43*) contextual modeling for protein–ligand affinity prediction.

The graph neural network module consists of two parallel branches based on PPFConv (*41*) and GENConv (*42*) operators (Section 6.3). Both branches process the same protein–ligand graph and produce 256-dimensional node representations, which are combined through concatenation before being passed to a transformer encoder.

The 4-layer PPFConv branch, with 2 local and global MLP operators, incorporates explicit geometric information through pairwise distance and orientation descriptors. In parallel, the 4-layer GENConv branch uses softmax-based neighborhood aggregation to update atom representations.

To improve information flow between layers, both branches employ manifold-constrained hyperconnections (*67*) with 4 streams and 10 Sinkhorn iterations. They mitigate feature over-smoothing and preserve discriminative node representations.

The concatenated 512-dimensional node embeddings are subsequently processed by a transformer encoder where the self-attention mechanism incorporates pairwise structural biases directly into the attention score computation (Section 6.4). This modification enables the model to account for spatial relationships during global information exchange.

Following transformer processing, a ligand-specific pooling layer aggregates only ligand atom embeddings. A learnable query vector attends over the ligand nodes to produce a single representation, which is subsequently mapped to a scalar binding affinity prediction through a linear readout layer.

###### Input graph construction

The protein-ligand graph uses the atoms as nodes and adds edges based on atom type-specific distances. The graph includes all ligand atoms and all protein heavy atoms within 8Å distance from any ligand atom. We add edges between protein-ligand nodes if their distance is less than 8Å, and also add ligand-ligand and protein-protein edges if the distance is less than 4Å.

###### Graph node and edge featurization

For the input graph’s nodes we use 3 sets of features: a binary indicator whether the corresponding atom belongs to the ligand or the protein pocket, AEV representations (Section 6.2), and a learnable embedding based on the atom element type. The same atom type vocabulary is used for the AEV features and the learnable embeddings. These feature sets are concatenated and projected to a 256-dimensional representation using a linear layer, followed by layer normalization.

For the graph edges we use modified PPF features (*41*), where we replace the central node’s surface norm with the average vector of the position differences with neighboring nodes (Section 6.3).

###### GNNs with manifold-constrained hyper-connections

Following application of hyper-connections in GNNs, each node *i* is represented not by a single feature vector but by a multi-stream state *x*_*i*_ ∈ *R*^*n*×*d*^, where *x*^(*s*)^ ∈ *R*^*d*^ denotes the *s*-th stream and *n* is the stream expansion factor. An mHC-GNN layer combines two additive paths: a residual stream-mixing path and a message-passing path. The layer update is defined as

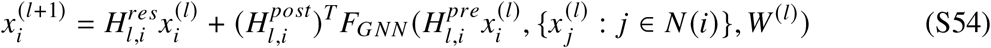

where *H*^*res*^ ∈ *R*^*n*×*n*^ is a doubly stochastic stream-mixing matrix, *H*^*pre*^ ∈ *R*^1×*n*^ aggregates the streams before message passing, *H*^*post*^ ∈ *R*^1×*n*^ expands the message-passing output back to the multi-stream space, and *F*_*GNN*_ is the underlying GNN operator.

Crucially, *H*^*res*^is constrained to the Birkhoff polytope, i.e., to the set of doubly stochastic matrices, by applying Sinkhorn-Knopp normalization to a learnable matrix. This constraint preserves feature means across streams and bounds signal propagation, which is the central mechanism used by the method to mitigate over-smoothing. Ablation results with or without hyper-connections can be found in Table S12.

###### Pooling and readout

The model operates on a joint graph representation of the protein pocket and ligand; however, the final prediction is conditioned solely on ligand-specific information. The ligand-only cross-attention pooling focuses the readout on the most relevant substructure for affinity prediction, as binding affinity is primarily determined by ligand-specific interactions conditioned on the pocket.

By restricting aggregation to ligand atoms, the model reduces noise from less informative protein regions while still leveraging protein context encoded during earlier joint processing, leading to more stable and discriminative representations.

The cross-attention pooler utilizes a learnable parameter vector *h̃*_cls_ ∈ *R*^*d*^ as a base query token. To condition this token on the input complex, we apply the adaptive layer normalization mechanism used in AlphaFold 3 (*68*). Specifically, we first compute a global context vector *s* ∈ *R*^*d*^ by averaging the transformer output embeddings of all *N* protein and ligand nodes:

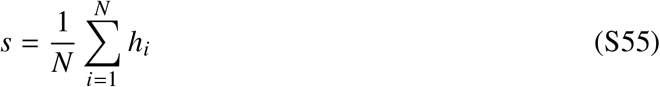

where *h*_*i*_∈ *R*^*d*^ represents the embedding of node *i*. The final conditioned query token *h*_cls_ is then computed by modulating the learnable query embedding *h̃*_cls_ using the context vector:

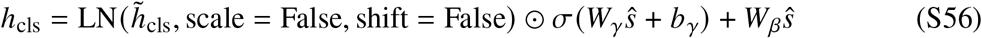

where *ŝ* = LN(*s*, shift = False) is the normalized context vector, *σ*(·) is the element-wise sigmoid function, and {*W*_*γ*_, *b*_*γ*_, *W*_*β*_} are the learnable linear projection parameters.

Let *H*_*L*_ ∈ *R*^*N*^^*L*×*d*^ be the set of ligand node embeddings after the transformer, where *N*_*L*_ is the number of ligand atoms. Cross-attention pooling is then applied as follows:

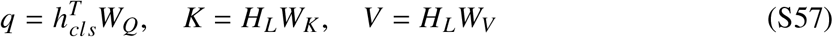

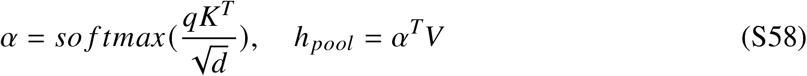

where *W*_*Q*_, *W*_*K*_, *W*_*V*_ ∈ *R*^*d*×*d*^ are learnable projection matrices. This results in a single pooled representation *h*_*pool*_ ∈ *R*^*d*^. The readout layer maps this representation to a scalar affinity prediction:

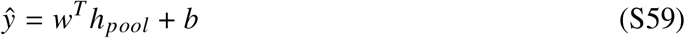

where *w* ∈ *R*^*d*^and *b* ∈ *R* are learnable parameters. The ablation results of pooling techniques can be found in Table S12.

##### 8.3 Training and model selection

We train DOScore using two different approaches: (i) for hit identification, (ii) for fine-grained scoring. Both variants share the same model architecture and general hyper-parameters, but employ different data sampling and optimization strategies.

We train all models for up to 10,000 steps using effective batch size of 256 protein-ligand complexes. During training, we maintain an Exponential Moving Average (EMA) of the network weights. To reduce variance and improve generalization, we save intermediate EMA checkpoints at 2,000, 4,000, 6,000, 8,000, and 10,000 steps. The final checkpoint for each training run is then produced by averaging the weights across these five saved snapshots.

The models are trained using a combination of AdamW (*69*) and Muon (*70*, *71*) optimizers: we apply Muon exclusively to multi-dimensional weight matrices where all dimensions are at least 128, while routing everything else to AdamW. The learning rate is 2 × 10^−4^, with a 1000-step warmup and a cosine annealing scheduler (*72*).

###### Hit identification

For training the hit identification variant of DOScore, we use MSE-based loss (Eq. S60, S61) where the decoy complexes are assigned a fixed affinity value equal to 3.0, but it zeroes out penalties for decoys predicted below this value. This design strictly punishes false positives while ignoring variations in how inactive a decoy is predicted to be.

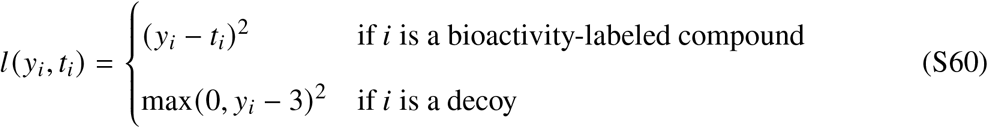

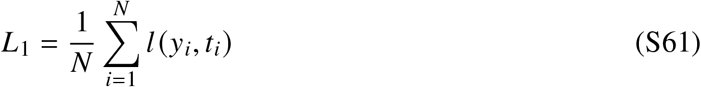

where *y*_*i*_ is the predicted score, *t*_*i*_ is the target value.

To construct training mini-batches, we employ a hierarchical sampling strategy that scales interassay representation logarithmically relative to individual assay size. This mitigates the training dominance of massive assays. Within each individual assay, we counteract severe class imbalances between bioactivity-labeled and decoy ligands by distributing weights across classes, with 25% allocated to labeled ligands and 75% to decoys.

###### Fine-grained scoring

Our scoring model is, by design, assay-independent: it predicts the affinity score without explicit information about the experimental setup. However, the training data combines measurements such as *K*_*i*_ and IC_50_ from diverse assays with different conditions and endpoints. Such variation introduces assay-specific biases, which add noise in standard regression setting. In order to train the fine-grained variant of DOScore, we mitigate this issue by combining an intra-assay mean-centered regression loss (Eq. S63) with a softplus pairwise ranking penalty evaluated strictly on bioactivity-labeled ligands (Eq. S64, S65).

Because *K*_*i*_ and IC_50_ are related through the Cheng–Prusoff equation (*73*), subtracting the assay-specific mean in log space effectively cancels the correction term, provided that substrate concentration [*S*] and Michaelis constant *K*_*m*_ remain constant within a given assay:

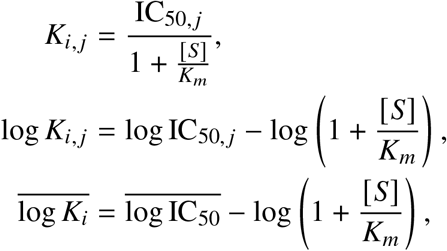

which directly implies:

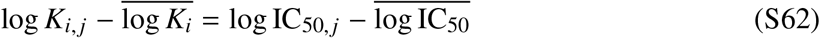

Translating this principle into our optimization objective, the regression loss centers predictions and targets within each assay to focus on relative intra-assay variations:

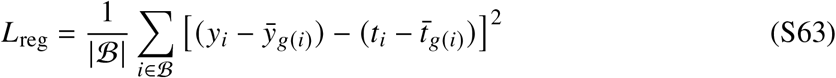

where B represents the current mini-batch of training compounds, where *y*_*i*_ and *t*_*i*_denote the model’s predicted affinity score and the log-transformed experimental target value for compound *i*, respectively. The function *g*(*i*) maps compound *i* to its specific assay group, forcing the predicted means (*ȳ*_*g*(*i*)_) and target means (*t̄*_*g*(*i*)_) to be computed locally using only the samples from that assay present within the mini-batch.

Alongside regression, the ranking loss component penalizes misordered pairs within the same assay whose true affinities differ by more than a set margin, directly optimizing local ranking metrics. This component applies a softplus margin penalty over the set of all valid intra-assay pairs P (Eq. S64). The ranking pair set is defined as P = {(*i*, *j*) ∈ B × B | *g*(*i*) = *g*(*j*) and *t*_*i*_ > *t* _*j*_ + *m*}. Here, *m* is the strictly enforced ranking margin (set to 0.1 in our runs).

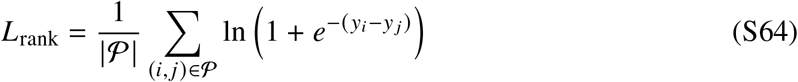

Finally, for this variant of the model, we combined the intra-assay mean-centered regression loss and the pairwise ranking penalty into a joint optimization objective *L*_2_:

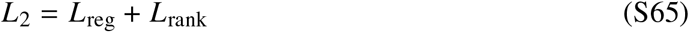

The mini-batch selection algorithm for the fine-grained scoring model also assigns assay-level sampling weights. It merges all structural clusters within each unique assay into a single, de-duplicated pool consisting exclusively of bioactivity-labeled ligands, assigning each assay a selection probability proportional to the square root of its total pool size. To construct a mini-batch, the sampler uses the precomputed weights to sample 16 unique assays without replacement and then draws up to 16 ligands uniformly at random from each assay’s pool. If the resulting batch is smaller than the target size of 256 — because some assays have fewer than 16 ligands — additional assays are sampled to fill the remainder.

#### 9 Experimental validation

##### 9.1 Prospective docking and X-ray validation of PCSK9 AZD-0780 complex

###### 9.1.1 Identification of the AZD-0780 Binding Site on the PCSK9 C-terminal Domain

AZD-0780 was docked into 16 publicly available PCSK9 structures containing an intact C-terminal domain (CTD), and the resulting binding modes are summarized in Figure S12. Docking converged to a consensus binding mode in 12 of the 16 structures. The remaining four structures either adopted a flipped ligand orientation (3SQO) or corresponded to engineered constructs (5VLP, 6U2F, and 6U3I) with distorted CTD conformations that occlude the binding pocket.

**Figure S12:**
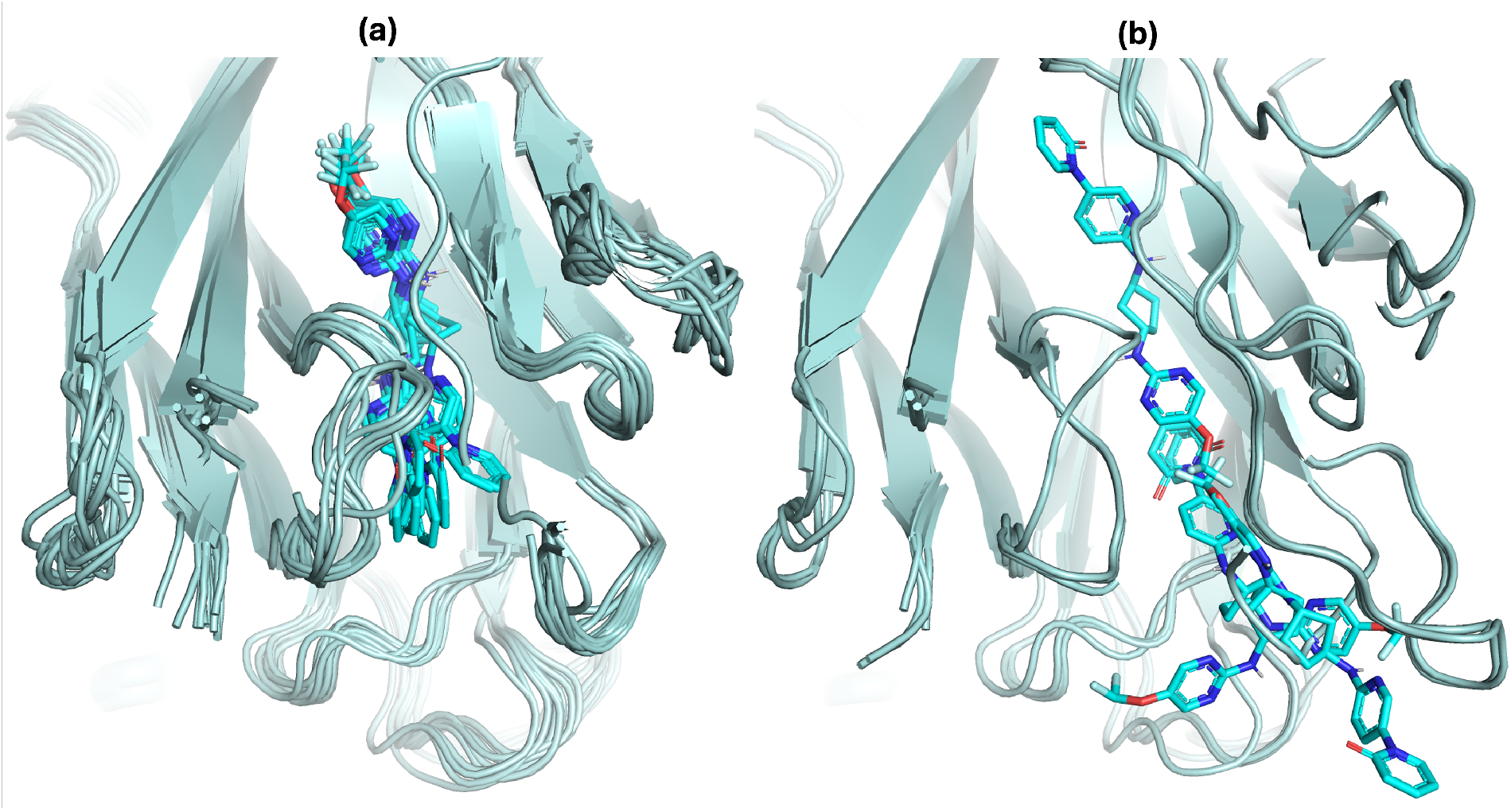
Ensemble docking across PCSK9 C-terminal domain structures. (a) Native human PCSK9 structures producing a consistent binding mode (2P4E (*74*), 2PMW (*75*), 3H42 (*76*), 3MOC (*77*), 3P5C (*78*), 4K8R (*79*), 4NE9 (*80*), 5OCA (*81*), 6U36, 6U38, 6U3X (*82*), 7ANQ (*83*)). (b) Structures producing inconsistent poses, including a flipped orientation in 3SQO (*84*) and distorted binding pockets in the engineered constructs 5VLP (*85*), 6U2F, and 6U3I (*86*).

###### 9.1.2 PCSK9 expression and purification

A human PCSK9 construct corresponding to residues Gln31–Gln692 of UniProt Q8NBP7 was expressed with an N-terminal honeybee melittin secretion signal and a C-terminal His_6_ tag. The construct used for structural studies contained the V474I and G670E substitutions. The expressed protein comprised the PCSK9 prodomain and catalytic/C-terminal domains and retained the C-terminal affinity tag.

Secreted C-terminally His-tagged PCSK9 was purified from insect-cell conditioned medium by immobilized metal-affinity chromatography followed by size-exclusion chromatography. The final purified protein was formulated in 20 mM Tris-HCl, pH 8.0, 300 mM NaCl, concentrated to 11 mg ml-1 by A_2_80, aliquoted, flash-frozen and stored at -80 °C.

Purity and homogeneity were assessed after one freeze–thaw cycle. Reducing SDS–PAGE showed >95% purity, with bands corresponding to the mature prodomain and catalytic/C-terminal domain. Analytical size-exclusion chromatography on a Superdex 200 Increase 10/300 column, equilibrated in the storage buffer, showed a monodisperse elution profile without evidence of aggregation, degradation or higher-order oligomerization. The purified protein was present as the mature non-covalent PCSK9 complex generated by autolytic cleavage at the expected site between residues 152 and 153. Protein identity and integrity were confirmed by intact-mass LC–MS and peptide mass fingerprinting. Intact-mass analysis detected the mature prodomain and catalytic/C-terminal domain and was consistent with partial phosphorylation of the prodomain and glycosylation of the catalytic domain.

###### 9.1.3 PCSK9 crystallization, data collection and structure determination

Human PCSK9 in complex with AZD0780 was crystallized using a construct comprising residues 31–692 and containing the V474I and G670E substitutions; the purification tag was left uncleaved. Purified PCSK9 at 11.0 mg ml^−1^ in 20 mM Tris-HCl, pH 8.0, 300 mM NaCl was pre-incubated for 1 h with AZD0780 at 1.5 mM, corresponding to an approximately tenfold molar excess of compound. Crystals were obtained at 20 °C by sitting-drop vapor diffusion by mixing protein–ligand complex with an equal volume of reservoir solution containing 0.10 M MD Buffer System 1, pH 5.8, 13% MD Precipitant Mix 2, 2.5% (v/v) MD Cryopolyols and 90 mM MD LiNaK Mix. Crystals suitable for diffraction were obtained after 7 d.

Crystals were cryoprotected by addition of ethylene glycol to a final concentration of 15% before flash-cooling. X-ray diffraction data were collected at the European Synchrotron Radiation Facility, Grenoble, France, on beamline ID23-1. Data were collected at a wavelength of 0.91508 Å from a crystal of PCSK9–AZD0780 in space group P2_1_22_1_, with unit-cell parameters a = 107.0 Å, b = 109.4 Å, c = 122.6 Å, *α* = *β* = *γ* = 90^◦^. The data were initially integrated and scaled using auto-PROC and SCALA/AIMLESS, giving a complete isotropic dataset to 3.13 Å resolution. Because the diffraction was anisotropic, the data were further processed using autoPROC/STARANISO, yielding anisotropic diffraction limits of 2.13, 3.76 and 2.60 Å along a*, b* and c*, respectively, and an effective high-resolution limit of 2.07 Å. The STARANISO-processed dataset had an overall ellipsoidal completeness of 92.5%, mean *I*/*σ*(*I*) of 4.2, multiplicity of 11.5 and CC1/2 of 0.982.

The structure was solved by molecular replacement using a published PCSK9 structure as the search model (PDB 2QTW). Iterative model building and refinement were performed using manual rebuilding and REFMAC5. Ligand restraints for AZD0780 were generated with ACEDRG from the CCP4 suite. Atomic displacement parameters were refined using one isotropic B factor per atom and TLS groups defined per protein chain. The final model was refined against the STARANISO-processed data to 2.07 Å resolution and contained 8,833 atoms, with an average B factor of 35.9 Å^2^. The final refinement statistics were *R*_*work*_ = 0.327 and *R* _*f* *ree*_ = 0.405. Model geometry was good, with root-mean-square deviations from ideality of 0.002 Å for bond lengths and 0.802° for bond angles. Ramachandran analysis showed 95.2% of residues in favored regions, 4.8% in allowed regions and no outliers.

The asymmetric unit contained two PCSK9 molecules, each comprising the inhibitory prodomain and the catalytic/serine protease plus C-terminal cysteine-rich domains. Electron density was best defined for chains A and B, whereas chains C and D were more weakly ordered. Clear Fo – Fc difference density was observed for AZD0780 in the binding site of chain B before ligand modelling, enabling placement of the full compound. The final model includes AZD0780 bound in the C-terminal cysteine-rich domain of PCSK9.

Table S16 summarizes the data collection and refinement statistics.

**Table S16:** PCSK9 crystallization data collection and refinement statistics. Data was collected from one crystal. Values in parentheses are for the highest-resolution shell. Completeness is reported as ellipsoidal completeness from STARANISO processing because the dataset was anisotropic. The ligand/ion atom count includes 30 ligand atoms and two calcium ions.

| **Data collection** |  |
| --- | --- |
| Space group | $P2_122_1$ |
| Cell dimensions |  |
| $a, b, c$ (Å) | 106.990, 109.447, 122.602 |
| $\alpha, \beta, \gamma$ (°) | 90.0, 90.0, 90.0 |
| Resolution (Å) | 109.45–2.07 (2.45–2.07) |
| $R_{\text{merge}}$ | 0.543 (1.710) |
| $I/\sigma I$ | 4.2 (1.6) |
| Completeness (%) | 92.5 (66.8) |
| Redundancy | 11.5 (11.2) |
| **Refinement** |  |
| Resolution (Å) | 109.45–2.07 |
| No. reflections | 33,926 |
| $R_{\text{work}} / R_{\text{free}}$ | 0.327 / 0.405 |
| No. atoms |  |
| Protein | 8,297 |
| Ligand/ion | 32 |
| Water | 504 |
| $B$-factors (Å<sup>2</sup>) | |
| Protein | 39.04 |
| Ligand/ion | 17.70 |
| Water | 24.94 |
| R.m.s. deviations |  |
| Bond lengths (Å) | 0.002 |
| Bond angles (°) | 0.802 |

###### 9.1.4 Validation of the AZD0780 binding pose

Although the global refinement residuals were comparatively high *R*_*work*_= 0.327 and *R* _*f* *ree*_ = 0.405, the placement of AZD0780 was supported by multiple local measures that were evaluated independently of the overall model statistics. The elevated global residuals were associated with pronounced diffraction anisotropy and weak electron density for the second PCSK9 molecule in the asymmetric unit, whereas the PCSK9 copy comprising chains A and B, including the AZD0780-binding site in chain B, was substantially better ordered.

Before inclusion of AZD0780 in the atomic model, refinement of the ligand-free structure produced contiguous positive (Fo-Fc) difference density at the binding site that encompassed the full length of the compound and permitted an unambiguous assignment of its orientation. Subsequent refinement yielded continuous (2Fo-Fc) density around the ligand and preserved the hydrogen-bonding and hydrophobic interactions expected for the refined pose. The ligand was modeled at an occupancy of 1.00, with a mean atomic B factor of 16.9 Å^2^, closely matching the mean B factor of 18.5 Å^2^ for protein atoms within 4 Åof the ligand. The similarity between ligand and local protein B factors argues against placement of a poorly ordered or substantially overfitted ligand.

The asymmetric unit contained two PCSK9 molecules; however, interpretable ligand density was observed only in the better ordered chain-B binding site. The corresponding region of chain D was globally more weakly defined and did not support reliable placement of a second AZD0780 molecule. The chain-D site was therefore left unoccupied rather than imposing a ligand model in the absence of sufficient experimental density. Consequently, the chain-B ligand pose is supported by the local experimental data but is not independently replicated by the second crystallographic copy.

The overall chain-B structure superimposed on the reference PCSK9 structure 2QTW with a *C*_*α*_ r.m.s.d. of 1.1 Åover 473 aligned residues. The V474I substitution is spatially separated from AZD0780, with a minimum ligand–residue distance of approximately 16.6 Å, and does not make direct contacts with the compound. Residues 660–670, including the G670E substitution, were disordered and were not included in the final model. Thus, the structure supports the conclusion that the resolved ligand-binding site is not detectably distorted relative to the reference PCSK9 fold, but it does not permit a direct structural assessment of the G670E side chain itself.

The atomic coordinates and structure factors have been deposited in the Protein Data Bank under accession code 37HF.

##### 9.2 Virtual screening

###### 9.2.1 Virtual screening workflow

DODock and DOScore were utilized to screen two chemical spaces: the Enamine REAL Space synthon-based library, comprising 70 billion compounds, and the Enamine REAL database, containing 9.6 billion fully enumerated compounds. To navigate the synthon-based library, we deployed a reinforcement learning-based sampler designed to optimize docking scores by selecting the most favorable synthons during molecule generation. Conversely, an active learning sampling strategy was applied to the enumerated database. For both virtual screening workflows, approximately 5 million molecules were docked and scored.

###### 9.2.2 CD73

The prospective virtual screening workflow is summarized in Figure 4. CD73 presents a particularly challenging target for structure-based virtual screening because the active site contains two catalytic *Zn*^2+^ ions and represents a phosphate-binding pocket, both of which complicate accurate scoring of ligand interactions while the active site undergoes substantial conformational changes during catalysis.

To account for the substantial conformational flexibility of the CD73 active site, particularly around the N186 and N499 gating regions, an ensemble docking strategy was developed using multiple experimentally determined crystal structures representing distinct active-site conformations, including 6XUE chain A (*87*), 6YE2 chain A (*88*), 6S7F chain A (*89*), and 7JV8 chain C (*90*). Prior to docking, loop modeling and pKa calculations were performed to optimize protein structures and assign appropriate amino acid protonation states. Docking calculations were performed in both the presence and absence of phosphate to capture ligand-induced changes in active-site architecture.

To guide ligand recognition, all available CD73 co-crystal structures together with literature-reported inhibitor complexes were structurally aligned to identify conserved binding interactions. This analysis revealed three recurring pharmacophore features shared by virtually all potent non-nucleotide inhibitors: (i) an aromatic ring positioned between F417 and F500, forming the characteristic *π*-stacking interaction observed in co-crystal structures, (ii) a hydrogen-bond acceptor interacting with the side chain of N390, and (iii) a hydrogen-bond donor interacting with D506. These conserved interaction motifs are consistent with crystallographic observations and structure-activity relationship studies of nucleotide and non-nucleotide CD73 inhibitors (*91*) and were incorporated as key pharmacophore constraints during virtual screening.

Structural alignment additionally revealed a highly conserved water-filled subpocket adjacent to L415, occupied by ordered water molecules across all available CD73 crystal structures irrespective of ligand or protein conformation. To preserve this conserved hydration site during virtual screening, four exclusion spheres (radius 1.5 Å) were introduced to represent the steric volume occupied by these water molecules. This design was motivated by the consistent structural observation that productive CD73 ligands preserve this hydration network and was further supported by published structure-activity relationship studies demonstrating that bulky substituents projecting toward this region are poorly tolerated (*87*).

Application of the virtual screening workflow generated approximately 100,000 virtual hits (Figure 4). Compounds were subsequently prioritized using binding affinity score, a proprietary machine learning-based aqueous solubility (LogS) prediction model, pharmacophore compliance, ligand strain energy assessment, and molecular dynamics-based pose stability. Candidate molecules were additionally counter-docked against the open-state CD73 structure (PDB 4H2F) to eliminate compounds predicted to preferentially bind nonproductive open conformations. Compounds exhibiting significant disagreement between predicted poses across open and closed receptor conformations (RMSD > 2.5 Å), poor molecular dynamics stability (MD RMSD > 4.0 Å), or unfavorable physicochemical properties were removed during post-processing.

Following automated prioritization, approximately 1,500 compounds were selected for detailed manual inspection by experienced computational chemists. Binding modes, interaction networks, synthetic accessibility, scaffold novelty, and overall medicinal chemistry attractiveness were evaluated. To maximize chemical diversity and avoid rediscovery of previously reported inhibitor classes, complementary clustering and novelty analyses were subsequently performed. Scaffold diversity was assessed by Bemis-Murcko scaffold clustering using ECFP4 fingerprints with a Tanimoto similarity threshold of 0.5, while whole-molecule diversity was evaluated using ECFP4 fingerprints with a Tanimoto similarity threshold of 0.4. Finally, all prioritized compounds were compared against the set of 1,247 curated CD73 inhibitors. Molecules exhibiting an ECFP4 Tanimoto coefficient below 0.35 relative to every known CD73 inhibitor were classified as putatively novel chemotypes and were preferentially advanced for experimental evaluation.

The combined computational and expert-driven triage process yielded 220 compounds for procurement, of which 183 compounds were successfully acquired and experimentally tested. As summarized in Figure 4, biochemical screening identified 56 compounds with *IC*_50_ values below 500*μM*, including 54 compounds with biochemical *IC*_50_ < 100*μM* and 9 compounds with biochemical *IC*_50_ < 10*μM*. Cellular profiling identified 14 compounds with cellular *IC*_50_ values below 100*μM*, including 8 compounds with cellular *IC*_50_ < 10*μM*, with one compound exhibiting 570 nM cellular potency. Notably, most confirmed hits displayed low structural similarity to previously reported CD73 inhibitors.

###### CD73 biochemical activity assay

CD73 activity was measured using a malachite-green phosphate detection assay. Recombinant human CD73 (5’-nucleotidase, NT5E) was obtained from R&D Systems (5795-EN-010), 5’-AMP disodium salt from Sigma-Aldrich (01930-5G), and BIOMOL Green reagent from Enzo Life Sciences (BML-AK111-1000). Reactions were performed in assay buffer containing 50 mM Tris-Cl, pH 7.5, 100 mM NaCl, 1 mM MgCl_2_, 1 mM CaCl_2_, 5*μM* ZnCl_2_ and 0.005% Brij-35.

Compounds were dispensed acoustically using an Echo dispenser and screened either in a two-dose format at 100*μM* and 10*μM* or, for selected positive hits, in a 10-point dose-response format. Dose-response experiments were typically performed over a final compound concentration range of 200 to 0.01*μM*, in an assay buffer containing 1% v/v DMSO. Reactions were performed in a final volume of 30*μl* containing 0.06*nM* CD73 and 8*μM* 5’-AMP. Compounds were preincubated with CD73 for 30 min at 23 °C before reactions were initiated by addition of 5’-AMP. After 60 min at 23 °C, reactions were quenched by addition of 30*μl* BIOMOL Green reagent and incubated for a further 60 min at 23 °C. Absorbance was measured at 620 nm using an EnVision plate reader equipped with a monochromator.

Positive- and negative-control wells included enzyme and substrate in the absence of inhibitor, or substrate without enzyme, respectively. PSB-12379 and CD73-IN-5 were used as CD73 inhibitor controls (MedChemExpress, HY-100747 and HY-145334, respectively). Percent inhibition was calculated relative to the positive and negative controls, and *IC*_50_ values were determined by fitting concentration–response data with a four-parameter variable-slope nonlinear regression model. Compounds were considered positive hits if *IC*_50_ was < 100*μM* and Max Inhibition > 30%.

###### Cell-based CD73 activity assay

Cell-surface CD73 activity was measured in U87MG cells using a malachite-green phosphate detection assay. U87MG cells, which express high levels of CD73, were detached with Accutase, washed, resuspended in assay buffer and plated at 1,500 cells per well in 80*μl* in 96-well V-bottom polypropylene plates (Corning, 3363). The assay buffer contained 20 mM HEPES, pH 7.5, 137 mM NaCl, 5.4 mM KCl, 1.3 mM CaCl_2_, 4.2 mM NaHCO_3_ and 0.10% glucose.

Compounds were dispensed directly from DMSO stocks using a Tecan D300e digital dispenser, with DMSO normalized to 1% v/v in all wells. Compounds were screened either in a two-dose format at 100*μM* and 10*μM* or, for selected positive hits, in a 10-point dose-response format. Dose-response experiments were typically performed from 200*μM* using threefold serial dilutions. Cells were incubated with compounds for 60 min before addition of 25*μM* AMP substrate (Sigma-Aldrich, 01930-5G). After substrate incubation, plates were centrifuged at 225 g for 5 min, and 80*μl* supernatant was transferred to assay plates (Costar, 3370). Inorganic phosphate was detected using the Phosphate Assay Kit–PiColorLock™ (Abcam, ab270004) and according to the manufacturers instructions. Briefly, PiColorLock reagent containing accelerator was added to each well, plates were incubated for 5 min at room temperature, stabilizer was added, and plates were shaken for 5 min before a further 25-min incubation in the dark. Absorbance was measured at 620 nm using an EnVision multilabel plate reader.

DMSO-treated cells incubated with or without AMP were used as the maximum- or minimum-signal controls, respectively. Wells containing no cells were used for background subtraction. PSB-12379 and CD73-IN-5 were used as positive control. Beatty compound 33 (*87*) was used as an internal low-potency QC control. Percent inhibition was calculated relative to the maximum- and minimum-signal controls after background subtraction, and *IC*_50_ values were determined by fitting concentration-response data with a four-parameter logistic model.

###### 9.2.3 IRAK4

The prospective virtual screening workflow is summarized in Figure S13. IRAK4 represents a challenging kinase target for structure-based virtual screening due to the high conservation of the ATP-binding site and the propensity for false-positive enrichment driven by generic hinge-binding chemotypes.

**Figure S13:**
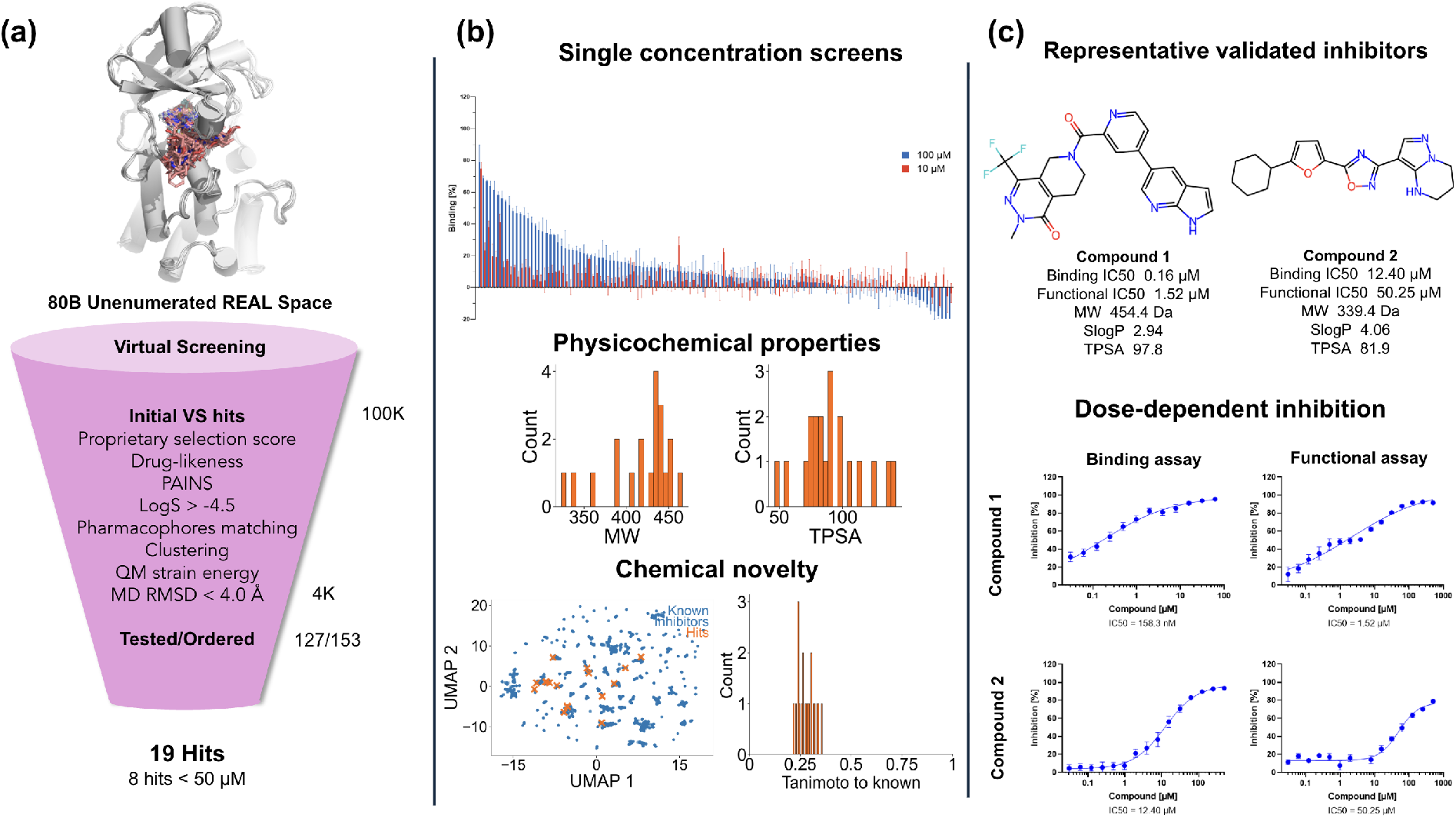
IRAK4 experimental validation results. **(a)** Prospective virtual screening workflow. Approximately 80 billion compounds from Enamine REAL Space were computationally screened and progressively prioritized to 153 compounds for procurement, of which 127 were experimentally evaluated. The number of compounds remaining after each filtering stage and the corresponding biochemical and cellular hit rates are shown. **(b)** Characterization of validated hits. Top, single-concentration biochemical inhibition measured at 100 and 10*μM*. Middle, molecular weight (MW) and topological polar surface area (TPSA) distributions of validated hits. Bottom left, UMAP projection of ECFP4 fingerprints comparing validated hits (orange) with previously reported IRAK4 inhibitors (blue). Bottom right, distribution of ECFP4 Tanimoto similarity between validated hits and the closest known IRAK4 inhibitor, illustrating the chemical novelty of the identified compounds. **(c)** Representative validated IRAK4 inhibitors. Chemical structures, biochemical and cellular *IC*50 values, physicochemical properties, and representative dose-response curves for two non-nucleotide inhibitors.

Structural analysis of available co-crystal structures revealed a conserved water molecule in the back pocket of the ATP-binding site, forming a stable hydrogen-bonding network with the side chains of E233 and D329. To preserve this interaction network, post-docking filtering was introduced to penalize or remove compounds that occupy this conserved water site. To guide ligand recognition, two pharmacophore constraints were defined for virtual screening: a hydrogen-bond acceptor interaction with the backbone of M265 in the hinge region, and an aromatic ring feature oriented toward the back pocket of the ATP-binding site.

To account for receptor flexibility, ensemble docking was performed using 4Y73_A (*92*), 8V2F_A (*93*), 6LXY_A (*94*), 8TVN_A (*95*), and 6O94_A (*96*) crystal structures, following loop modeling and pKa-based protonation state assignment. During virtual screening, the optimal receptor conformation was automatically selected per ligand based on docking score and pose consistency.

Following virtual screening, compounds were further prioritized using physicochemical property filters, quantum mechanics-based strain energy evaluation, and molecular dynamics-based pose stability. For chemical space characterization, the selected candidate set was compared against a curated set of approximately 320,000 kinase inhibitors with reported activities below 1*μM* (*IC*_50_, *K*_*i*_, *K*_*d*_, or *EC*_50_) from BindingDB. Across this reference set, the maximum ECFP4 Tanimoto similarity between screened compounds and known kinase inhibitors was 0.38, indicating that the prioritized compounds occupy a distinct region of kinase-relevant chemical space.

Experimental validation identified 19 structurally diverse chemotypes with TR-FRET *IC*_50_ values below 500*μM*, including 12 compounds with sub-100*μM* activity, 2 compounds with sub-10*μM* and one compound exhibiting 158*nM* potency. Notably, these hits span multiple scaffolds distinct from known IRAK4 inhibitor classes, confirming that the workflow enabled both enrichment of active compounds and expansion beyond the canonical kinase inhibitor chemical space.

###### IRAK4 TR-FRET competition binding assay

IRAK4 ligand binding was measured using a time-resolved fluorescence resonance energy transfer (TR-FRET) competition assay in which test compounds displaced fluorescent Kinase Tracer 222 (KT222, ThermoFisher Scientific, PV6121) from His-tagged IRAK4 (Sino Biological, 10735-H07B-50) and was observed by TR-FRET, through proximity of KT222 to anti-His Eu-W1024-labelled antibody (Revvity Health Sciences, AD0402).

Assays were performed in 384-well low-volume white flat-bottom NBS microplates (Corning, 3824). Reactions were carried out in LANCE Detection Buffer (Revvity Health Sciences) containing 10 nM His-IRAK4, 2 nM anti-His Eu-W1024 antibody, 50 nM KT222 and 2.5% v/v DMSO. For primary screening, compounds were tested in a two-dose format at 100*μM* and 10*μM*. Selected compounds were subsequently profiled in a 15-point concentration-response format using two-fold serial dilutions, with the highest final compound concentration reaching 500*μM*. Zimlovisertib (MedChemExpress, HY-19836) was used as a positive-control IRAK4 inhibitor.

Following overnight incubation at room temperature, TR-FRET was measured on a BioTek Synergy Neo2 plate reader in TRF-laser mode using 337 nm excitation and dual 620 nm/665 nm emission detection. Compound activity was calculated from the reduction in TR-FRET signal relative to DMSO and inhibitor control wells, and *IC*_50_ values for confirmed hits were determined by fitting concentration-response data with a four-parameter variable-slope nonlinear regression model. Compounds were considered positive hits if *IC*_50_ was < 500*μM* and Max Inhibition > 30%.

###### IRAK4 kinase activity assay

IRAK4 enzymatic activity was measured using the ADP-Glo Kinase Assay Kit (Promega), which quantifies ADP generated during the kinase reaction. Reactions were performed in 384-well low-volume white flat-bottom NBS microplates, containing 22.3 nM His-IRAK4, 100*ng*/*μL* Myelin Binding Protein, 200*μM* ATP and 2.5% v/v DMSO in a buffer containing 40 mM Tris-HCl, pH7.4, 20 mM MgCl2, 50*μM* DTT and 100*ng*/*μL* BSA. Briefly, His-IRAK4 and MBP were preincubated with compounds for 30 min at room temperature, followed by initiation of kinase reaction by addition of ATP and 1 hr incubation at room temperature. ADP production was detected using the ADP-Glo assay according to the manufacturer’s protocol, and luminescence was measured on a BioTek Synergy Neo2 plate reader.

Compounds were first screened in a two-dose format at 100*μM* and 10*μM*. Selected compounds were subsequently profiled in a 15-point concentration-response format using two-fold serial dilutions, with the highest final compound concentration reaching 500*μM*.

Compound activity was calculated from the reduction in ADP-Glo luminescence relative to DMSO and inhibitor-control wells. *IC*_50_ values for confirmed hits were determined by fitting concentration-response data with a four-parameter variable-slope nonlinear regression model.

###### 9.2.4 Factor XI

**Figure S14:**
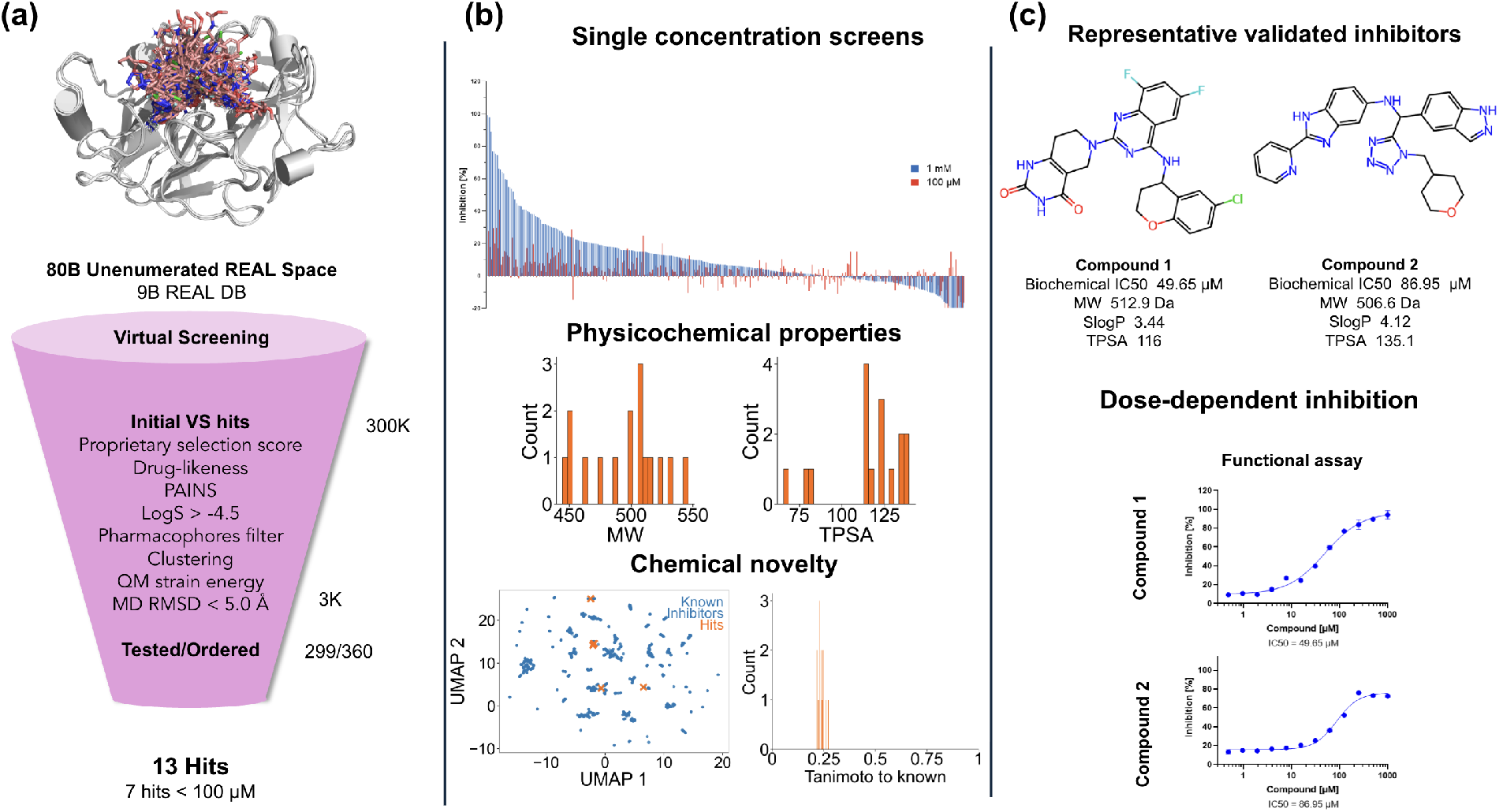
Factor XI experimental validation results. **(a)** Prospective virtual screening workflow. Approximately 80 billion compounds from Enamine REAL Space were computationally screened and progressively prioritized to 360 compounds for procurement, of which 299 were experimentally evaluated. The number of compounds remaining after each filtering stage and the corresponding biochemical hit rates are shown. **(b)** Characterization of validated hits. Top, single-concentration biochemical inhibition measured at 1*mM* and 100*μM*. Middle, molecular weight (MW) and topological polar surface area (TPSA) distributions of validated hits. Bottom left, UMAP projection of ECFP4 fingerprints comparing validated hits (orange) with previously reported FactorXIa inhibitors (blue). Bottom right, distribution of ECFP4 Tanimoto similarity between validated hits and the closest known FactorXIa inhibitor, illustrating the chemical novelty of the identified compounds. **(c)** Representative validated FactorXIa inhibitors. Chemical structures, biochemical *IC*_50_ values, physicochemical properties, and representative dose-response curves for two inhibitors.

The prospective virtual screening process is summarized in Figure S14. Factor XIa represents a challenging target for structure-based virtual screening because its active site is a shallow, solvent-exposed groove evolved to recognize an extended peptide substrate rather than a compact pocket, offering little enclosed volume for high-affinity ligand binding beyond the single S1 subsite.

First, 89 PDB structures with unique ligands bound to the active site were selected and clustered based on the geometry of the binding pocket. We found significant flexibility in the binding pocket outside the S1 cavity. However, we were able to select 4 representative structures with PDB ID 4NA7 (*97*), 4X6P (*98*), 7V17 (*99*) and 8BO7 (*100*) that captured the different pocket conformations well. When all 89 ligands were docked to these 4 selected structures in ensemble docking mode only 6 displayed RMSD larger than 2 Å. Thus, we adopted these structures for virtual screening.

From the resulting screen, 300,000 compounds were initially selected. The compounds then were filtered by their physico-chemical properties such as MW, LogS, LogP, TPSA, HBA, HBD, Fsp3, number of chiral centers and number of rotational bonds to ensure optimal starting properties for optimization and clustered to arrive at about 3000 compounds of interest. After removing compounds with poor molecular dynamics stability (MD RMSD > 5.0 Å) and high strain energy (> 25 kcal/mol) and manual curation 360 compounds were ordered, out of which 299 were successfully synthesised and sent for experimental testing. The kinetic fluorescence assay (see Methods) identified 13 novel hits (with max ECFP4 Tanimoto smaller than 0.22), 7 of which showing *IC*_50_ below 100*μM* with > 30% Max Inhibition.

###### Factor XIa functional activity assay

Factor XIa enzymatic activity was measured using a kinetic fluorescence assay based on cleavage of the fluorogenic peptide substrate IEGR-AMC. Human Factor XIa was obtained from Enzyme Research Laboratories (HFXIa 1111a), and IEGR-AMC was obtained from Bachem/MCE (4004961). Assays were performed in Greiner 384-well black small-volume non-binding plates (Greiner BioOne, 784900) in assay buffer containing 50 mM Tris, pH 7.5, 250 mM NaCl, 1 mM EDTA and 0.005% Tween-20.

Reactions contained 1 nM Factor XIa and 350*μM* IEGR-AMC, with compound and substrate stocks prepared in DMSO. Compounds were initially screened in a two-dose format at 1 mM and 100*μM*. Selected active compounds were subsequently profiled in a 9-point concentration-response format using three-fold serial dilutions. Milvexian, asundexian, frunexian and FD-IN-1 were used as Factor XIa inhibitor controls.

Substrate cleavage was monitored immediately after reaction initiation in kinetic mode over 2 h using excitation/emission wavelengths of 350 nm/445 nm. Fluorescence was measured every 5 min, and the slope of fluorescence increase over the linear range was used as a proxy of Factor XIa activity. Percent inhibition was calculated relative to positive- and negative-control wells, and *IC*_50_ values for confirmed hits were determined by fitting concentration-response data with a four-parameter variable-slope nonlinear regression model. Compounds were considered positive hits if *IC*_50_ was < 500*μM* and Max Inhibition > 30%.

###### 9.2.5 IL-17A

**Figure S15:**
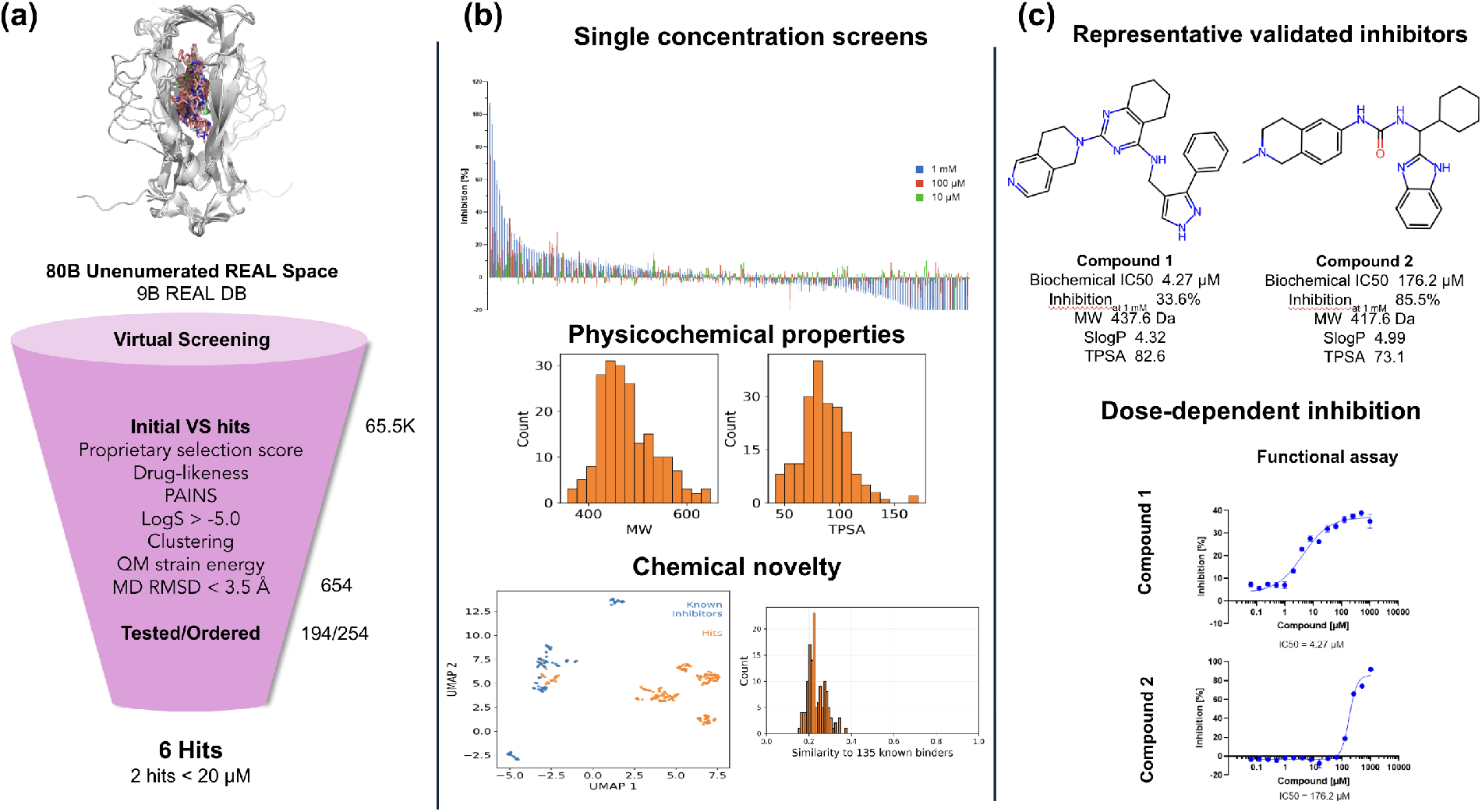
IL17A experimental validation results. **(a)** Prospective virtual screening workflow. Approximately 80 billion compounds from Enamine REAL Space were computationally screened and progressively prioritized to 254 compounds for procurement, of which 194 were experimentally evaluated. The number of compounds remaining after each filtering stage and the corresponding biochemical hit rates are shown. **(b)** Characterization of validated hits. Top, single-concentration biochemical inhibition measured in a three-dose format at 1 mM, 100 and 10 µM. Middle, molecular weight (MW) and topological polar surface area (TPSA) distributions of validated hits. Bottom left, UMAP projection of ECFP4 fingerprints comparing validated hits (orange) with previously reported IL-17A inhibitors (blue). Bottom right, distribution of ECFP4 Tanimoto similarity between validated hits and the closest known IL-17A inhibitor, illustrating the chemical novelty of the identified compounds. **(c)** Representative validated IL-17A inhibitors. Chemical structures, biochemical *IC*50 and maximum inhibition rate at 1 mM, physicochemical properties, and representative dose-response curves for two inhibitors.

The prospective virtual screening process is summarized in Figure S15. IL-17A represents a particularly challenging cytokine target for structure-based virtual screening because disruption of the IL-17A:IL-17RA complex is achieved indirectly by targeting a highly flexible allosteric pocket located at the IL-17A dimerization interface. Ligands binding to this site must induce or stabilize functionally relevant conformational changes that impair receptor recognition and prevent the formation of the IL-17A:IL-17RA complex.

To address the high conformational flexibility of the IL17A dimerization interface, an ensemble docking approach was developed. This method used multiple crystal structures of the IL17A homodimer, each representing different conformations of the dimerization interface, including the A and B chains from 7AMA (*101*), 8USR (*102*), 9FKX (*103*), 9H4D (*104*), and 9H4O (*104*). Before docking, all structures underwent loop modeling and pKa calculations to determine correct amino acid protonation states. Docking was performed in both the central and distal cavities of the dimerization interface to capture ligand-induced structural changes. To facilitate ligand recognition, all known IL17A co-crystal structures and literature-reported inhibitors were structurally aligned to identify conserved binding interactions.

Application of the virtual screening workflow generated approximately 65,500 virtual hits (Figure S15). These compounds were then clustered and prioritized based on docking scores, a proprietary machine learning-based aqueous solubility (LogS) prediction model, pharmacophore compliance, ligand strain energy assessment, and molecular dynamics-based pose stability. Compounds with poor molecular dynamics stability (MD RMSD > 3.5 Å) and high strain energy (> 25 kcal/mol) were excluded. Additionally, during post-processing, compounds were retained only if they showed no PAINS alerts, acceptable predicted solubility (LogS ≥ −5.0), lipophilicity within the predefined range (LogP or cLogP ≤ 7), and physicochemical properties, including 20-50 heavy atoms, ≤ 15 rotatable bonds, ≤ 5 hydrogen bond donors, ≤ 10 hydrogen bond acceptors, TPSA ≤ 180 Å^2^, ≤ 4 chiral centers, and FractionCsp^3^ ≥ 0.1. Additional elemental composition filters were applied to exclude compounds containing phosphorus, silicon, or iodine, while limiting the numbers of bromine, chlorine, fluorine, and sulfur atoms to ≤ 1, ≤ 2, ≤ 6, and ≤ 2, respectively.

Following automated prioritization, 654 compounds were chosen for detailed manual inspection by computational chemists. Binding modes, interaction networks, synthetic accessibility, scaffold novelty, and overall medicinal chemistry attractiveness were evaluated. To maximize chemical diversity and avoid rediscovering known inhibitor classes, additional clustering and novelty analyses were conducted. Scaffold diversity was measured using Bemis-Murcko scaffold clustering with ECFP4 fingerprints and a Tanimoto similarity threshold of 0.5, while whole-molecule diversity was evaluated with ECFP4 fingerprints and a Tanimoto similarity threshold of 0.4. All prioritized compounds were then compared to the set of 135 curated IL17A inhibitors. Molecules with an ECFP4 Tanimoto coefficient below 0.4 relative to all known IL17A inhibitors were identified as putatively novel chemotypes and preferred for experimental testing. This multi-stage prioritization strategy aimed to enhance predicted potency, structural diversity, and chemical novelty.

The combined computational and expert-driven triage process resulted in 254 compounds selected for procurement, of which 194 were successfully acquired and tested experimentally. As shown in Figure S15, biochemical screening identified 6 compounds with *IC*_50_ values below 500*μM* and more than 30% Max Inhibition of IL17A-IL17RA interaction, including 1 compound with a biochemical *IC*_50_ under 10*μM*. Notably, most confirmed hits had low structural similarity to previously reported IL17A inhibitors, indicating that the virtual screening effectively expanded beyond the limited scaffold space in earlier IL17A studies.

###### IL-17A/IL-17RA functional binding assay

Inhibition of IL-17A binding to IL-17RA was measured using a time-resolved fluorescence resonance energy transfer (TR-FRET) assay. Binding of biotinylated IL-17A (R&D System, BT7955B) His-tagged IL-17RA ECD (Sino Biological, 10895-H08H) generates a TR-FRET signal through proximity of Streptavidin–Alexa Fluor 647 (ThermoFisher Scientific, S32357) and anti-His Eu-W1024-labelled antibody. Test compounds reduced the TR-FRET signal by allosterically disrupting formation of the IL-17A:IL-17RA complex.

Assays were performed in white 384-well NBS microplates in LANCE Detection Buffer containing 5% v/v DMSO. Reactions contained 16 nM biotinylated IL-17A, 4 nM Streptavidin–Alexa Fluor 647, 16 nM His-IL-17RA ECD and 1nM anti-His Eu-W1024-labelled antibody. For primary screening, compounds were tested in a three-dose format at 1 mM, 100*μM* and 10*μM*. Selected active compounds were subsequently profiled in a 15-point concentration-response format using two-fold serial dilutions, with highest final compound concentration reaching 1 mM. LY3509754 (MedChemExpress, HY-139206) was used as a positive-control IL17A inhibitor.

Plates were incubated overnight at room temperature before TR-FRET readout. Compound activity was calculated from the reduction in TR-FRET signal relative to DMSO and assay control wells, and *IC*_50_ values for confirmed hits were determined by fitting concentration-response data with a four-parameter variable-slope nonlinear regression model. Compounds were considered positive hits if *IC*_50_ was < 500*μM* and Max Inhibition > 30%.

###### 9.2.6 Clustering of experimentally validated hits

Hit compounds were clustered manually by an experienced medicinal chemist following established medicinal chemistry practices. Most compounds were assigned using a hybrid approach that considered common synthetic routes, similarity of the core scaffold, and the orientation of exit vectors. Representative examples of such clusters include carbohydrates, 1,3-diaminopyrimidine derivatives, 1,2-disubstituted rigid-core scaffolds, compounds with flexible aliphatic core chains, star-shaped scaffolds, and 1,2,3-triazole derivatives together with their corresponding isosteres. In selected cases, functionally defined clusters were created based on shared chemical features rather than scaffold similarity. For example, compounds containing functional groups expected to coordinate or form ionic interactions with zinc ions were assigned to a dedicated metal-binding cluster.

The chemical structures for the hit clusters across all target proteins are provided in Section 3.1.

